# Global extent and change in human modification of terrestrial ecosystems from 1990 to 2022

**DOI:** 10.1101/2025.01.12.632633

**Authors:** David M. Theobald, James R. Oakleaf, Glenn Moncrieff, Maria Voigt, Joe Kiesecker, Christina M. Kennedy

## Abstract

Habitat loss and degradation associated with industrial development is the primary threat and dominant driver of biodiversity loss globally. Spatially-explicit datasets that estimate human pressures are essential to understand the extent and rate of anthropogenic impacts on ecosystems and are critical to inform conservation commitments and efforts under the Global Biodiversity Framework. We leveraged the human modification framework to generate comprehensive, consistent, detailed, robust, temporal, and contemporary datasets to map cumulative and individual threats associated with industrial human activities to terrestrial biodiversity and ecosystems from 1990 to 2022. In ∼2022, 43% of terrestrial lands had very low levels of modification, while 27%, 20%, and 10% had low, moderate, and high modification, respectively. Nearly ⅔ of biomes and ½ of ecoregions currently (∼2022) are moderately-modified, and 24% of terrestrial ecosystems (31 M km^2^) experienced increased modification from 1990 to 2020. About 29% of countries and 31% of ecoregions might also be particularly vulnerable to biodiversity loss given their above-average increased modification and less than 30% protection.

## Background & summary

Human activities and associated land uses have resulted in alteration, modification, fragmentation, degradation, and often conversion of natural habitats^1,2^, significantly contributing to the declines in biodiversity and nature’s benefits to people^3^. Habitat destruction and degradation is the primary threat to 88% of endangered and threatened species^4^ and is the dominant, direct driver of biodiversity loss globally (Jaureguiberry et al. 2022). To conserve biodiversity in the face of these challenges, we need to deepen our understanding of the global patterns and trends in these activities in order to guide conservation interventions that protect, connect, and restore ecosystems. Therefore, datasets that estimate human pressures have been developed and applied to understand the extent and/or change in the condition of natural systems^5,6,7,8,9,10,11^.

Although mapping human pressure does not directly measure realized ‘state’ of or ‘impacts’ on natural systems or their biodiversity, it provides a practical, surrogate measure of structural state and landscape characteristics^12,13,14^. Human pressure maps have been found to coincide with the loss, degradation, or impairment of ecosystems and its associated biodiversity^8,11,15^. Here we focus on global mapping of industrial pressures based on agriculture, forestry, transportation, mining, energy production, electrical infrastructure, dams, pollution and human accessibility^11^. We do not map traditional land use practices (e.g., shifting cultivation, agroforestry, hunting or harvesting of flora and fauna), which have had long history of use and modification by Indigenous Peoples and local communities and often been compatible with the maintenance of biodiversity and ecosystem services^3^, but are increasingly threatened by industrial development^16^.

Information from remotely-sensed data, especially land cover datasets, are central sources of information for human pressures mapping. Additional data from other sources (e.g., census, cartographic mapping), including geospatial analyses that generate key indicators, provide complementary information and are needed to provide robust and comprehensive understanding of human pressures^17^. This includes human activities that do not necessarily result in conversion (e.g., grazing^18^); have spectral characteristics that are difficult to distinguish from adjacent, un-modified areas (e.g., gravel/dirt roads that are constructed with similar materials to adjacent lands); are too small or thin to be detected at the resolution of the data source (e.g., small roads, trails, and fences^19^); are obfuscated by tree canopy (e.g., small roads, trails), are difficult to observe or are cryptic in nature (e.g., animal/plant collection or poaching^20^); or derive from characteristics of people, communities, and management activities^21^. Recent advances in remote sensing and machine learning are beginning to address some of these challenges (e.g., 1 m tree canopy models^22^), though they typically are founded on non-imagery georeferenced features, such as OpenStreetMap^23^. Yet, there remain practical limitations that preclude widespread use beyond more localized case studies (e.g., data not consistently available for the extent needed) and for conditions previous to the past few years.

Here we describe updated and refined global datasets on the degree of human modification of terrestrial ecosystems for contemporary (circa 2022) conditions (at 90 and 300 m) and from 1990 to 2020 (at 300 m) at 5-year increments (change maps). The human modification (HM) framework^6,8,10,24^ (Figure 1) and accompanying datasets quantify human activities that directly or indirectly alter natural systems and potentially cause negative stress or impairment to ecological systems. Generally, the HM framework proceeds by identifying datasets for each pressure (or threat – hereafter we use threat for consistency^25^); combining multiple datasets to calculate the spatial area (or footprint) for each threat; parameterizing the intensity value that is multiplied by the footprint to quantify human modification; and aggregating the threats into an overall measure of cumulative threat. Here, we also perform analyses to verify/validate the datasets and produce global HM datasets from 1990 to 2020 at 300-m and 90-m for 2022 and summarize patterns at country, realm, biome and ecoregional scales to support application of HM to a variety of uses.

**Figure 1.**
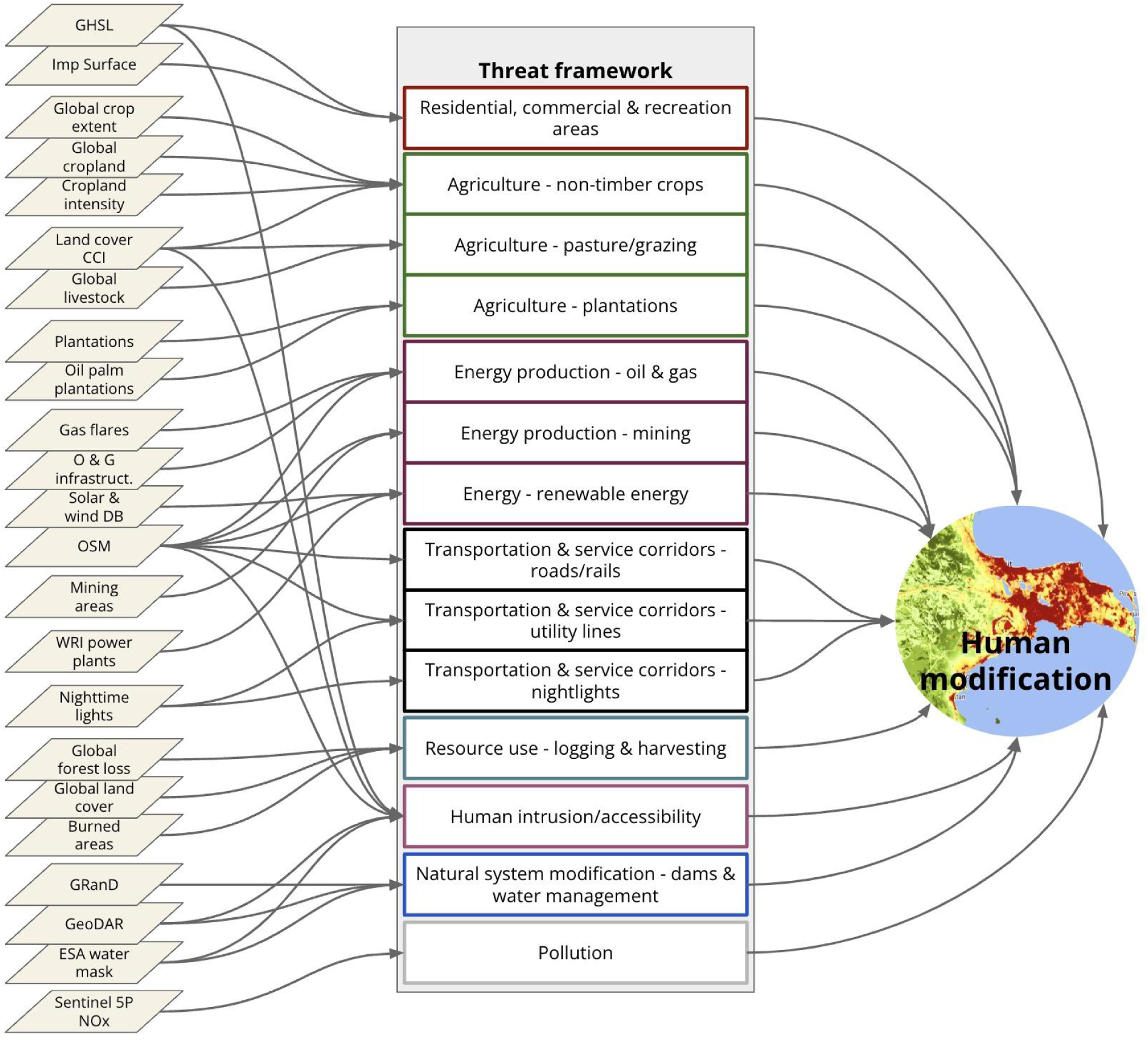
An overview of the global human modification datasets, following the structure of the IUCN threat framework (IUCN 2022). Acronyms: CCI= ESA Climate Change Initiative, DB=database; ESA=European Space Agency; GHSL=Global Human Settlement Layers; GRanD=Global Reservoir Database; GeoDAR= Georeferenced global Dams And Reservoirs; Global Reservoir and Dam database; O & G=oil and gas; OSM=Open Streetmap.

Key to the HM framework is the International Union for Conservation of Nature’s (IUCN) threat classification scheme (hereafter threat taxonomy^25,26,27^, which provides a comprehensive and parsimonious way to organize threats to be included in resulting maps and is foundational to the Convention on Biological Diversity (CBD) reporting procedures. The threat taxonomy is particularly valuable, because it provides a parsimonious list of threats to understand more specific (towards mechanistic) impacts on biodiversity and a basis to evaluate or better understand threshold values per threat. Practically, the taxonomy forms a foundation to guide decisions about what threats and representative datasets to include or exclude. The classifications are designed to be comprehensive, consistent, and exclusive for the first and second levels (Salafsky et al. 2008). We adopt this important distinction and call the first level of human threats the “threat class”, the second level “threat”, and finer-grained distinctions, typically represented by a single dataset, are placed into the relevant threat. For example, three datasets that represent urban areas such as buildings and population density would be considered to be a single threat (“housing and urban area”) within the “residential and commercial development” class.

We mapped direct threats^25^: “… the proximate human activities or processes that have caused, are causing, or may cause the destruction, degradation, and/or impairment of biodiversity targets (e.g., logging)”, but not threats, which are impaired attributes of a conservation target’s ecology that results directly or indirectly from human activities (e.g., fragmented habitat, impaired reproductive success, or degraded water quality). Similar to other efforts^5,7,8,28^, we augmented remotely sensed data with traditionally mapped cartographic features to provide a more comprehensive and complete dataset of global human threats.

Relative to previous versions of global HM datasets^8,10^, this version has: three additional threats with 13 new and 11 updated datasets; revised and standardized estimates of intensity values based on regional ecological feedback; refined methods to characterize pasture/grazing lands using an adaptive, land cover patch approach; and higher resolution (90 m) annual map for 2022 and an extended time period from1990 to 2020 at 5 year intervals.

The human modification datasets described here have significant advantages over previous efforts because they are:

- *comprehensive*, representing 16 threats as defined by the IUCN threat taxonomy;
- *globally consistent*, so that country, biome, ecoregional, and local evaluation and comparison are supported;
- *higher resolution,* with spatial data products used to generate the time-series data from 1990-2020 at 300 m (median resolution across data sets=30 m) and 90 m for 2022 (also median=30 m);
- *robust,* because threats are represented as fractional coverage within each pixel, with separate estimates of intensity, which are combined into a cumulative measure that minimizes confounding effects caused by double-counting similar datasets (i.e. does not assume independence among datasets and threats);
- *interpretable*, with HM values that are a “real” data type and consistent across the full gradient of modification, from unmodified or “wild” (0.0) to highly modified or developed (1.0); and
- *contemporary* (up-to-date) with the “latest-greatest” dataset represents nominally ∼2022 (median year of datasets =2022, to be updated on an annual basis).

For these reasons, these global human modification datasets are the most comprehensive, consistent, detailed, robust, and contemporary datasets currently available to map threats to terrestrial biodiversity and ecosystems.

## Methods

The new datasets presented here map 16 threats that are then organized by 8 threat classes (Table 1) and aggregated to map the cumulative degree to which human modification is present on global terrestrial lands (except Antarctica). Following previous work^6,8,10^, we calculated the degree of human modification value *H* for each threat *t* as the product of the spatial extent (or “footprint”) and the intensity of the land use:

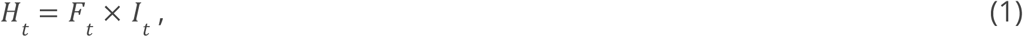

**Table 1.**
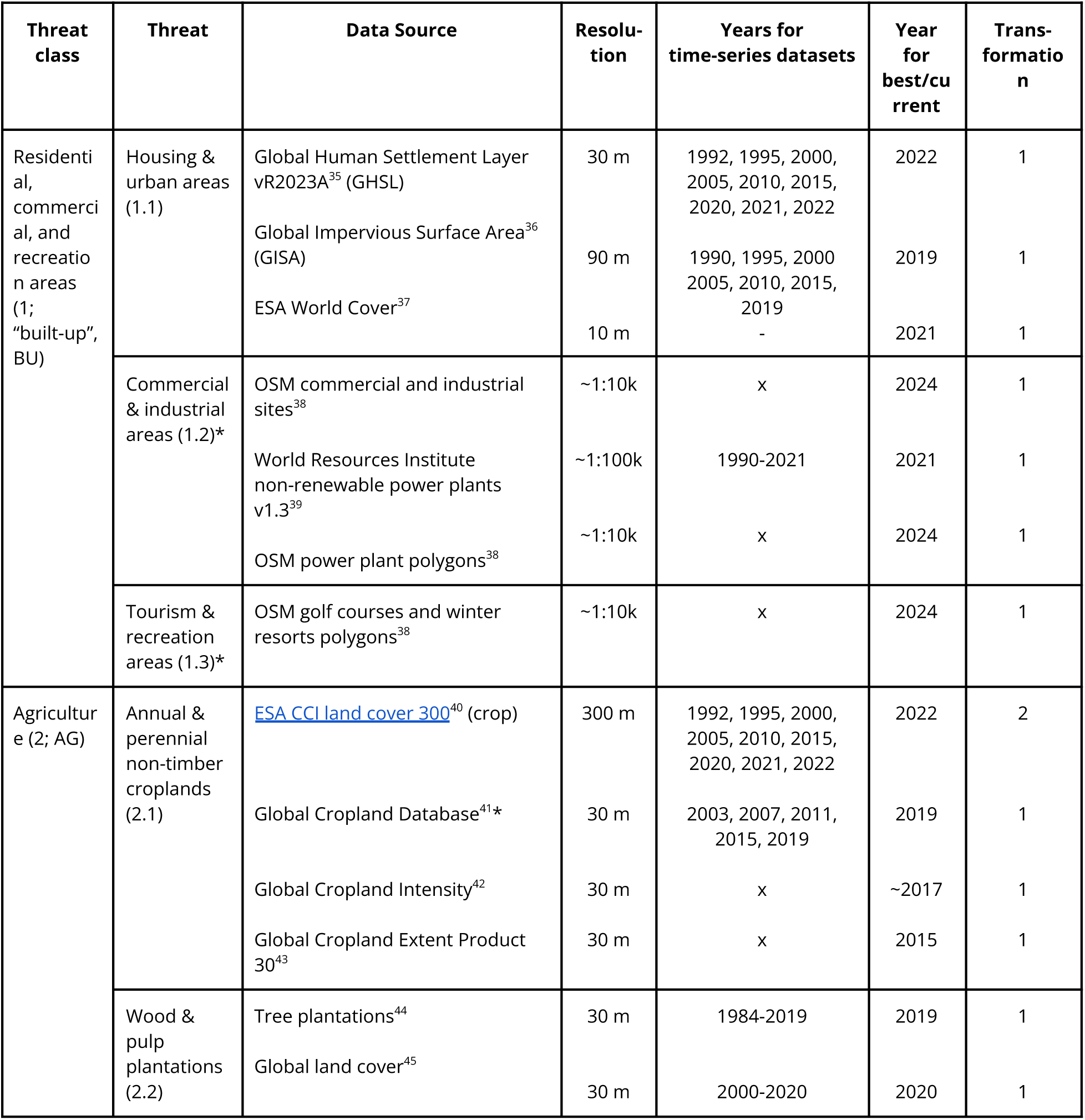
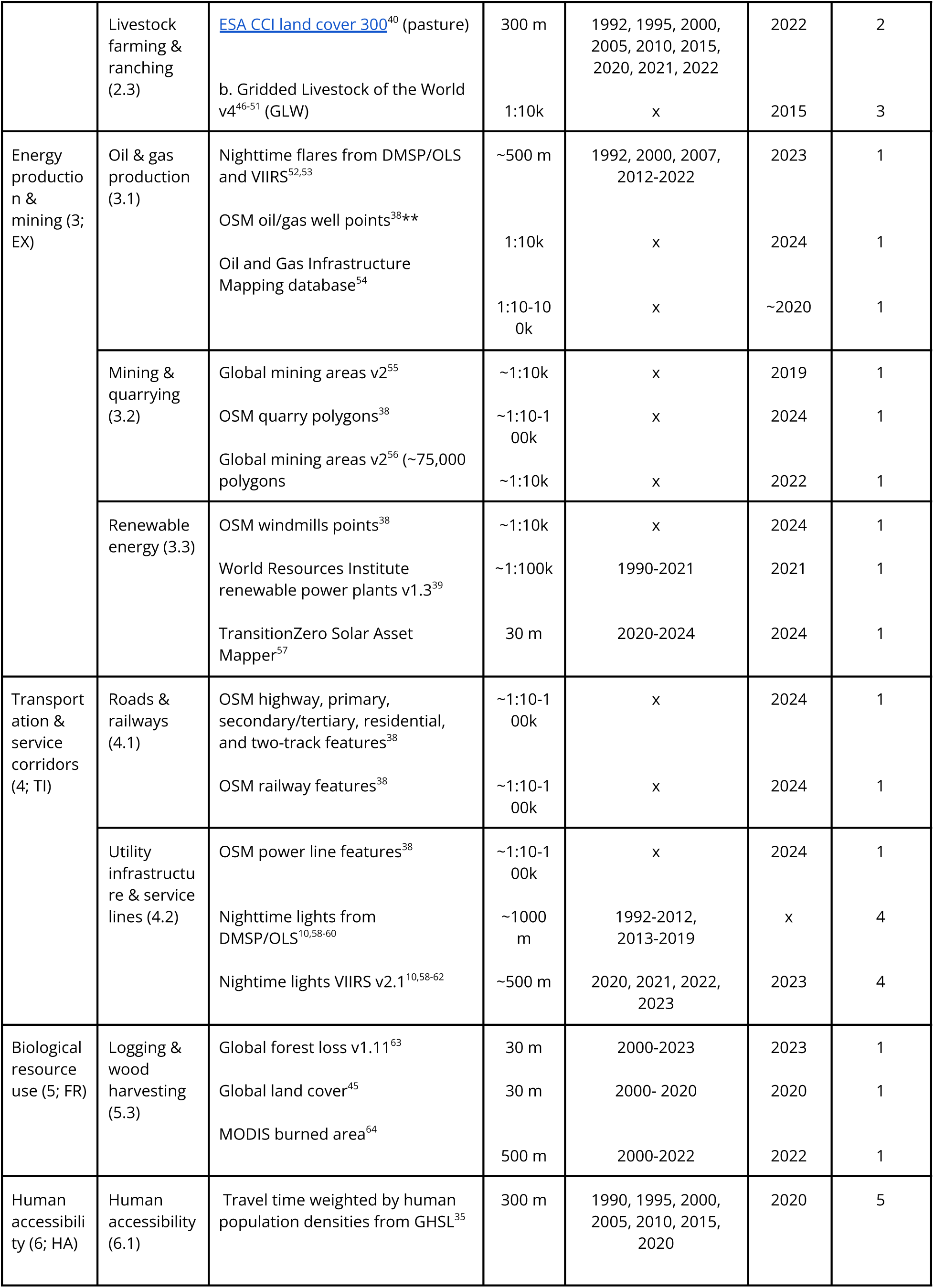
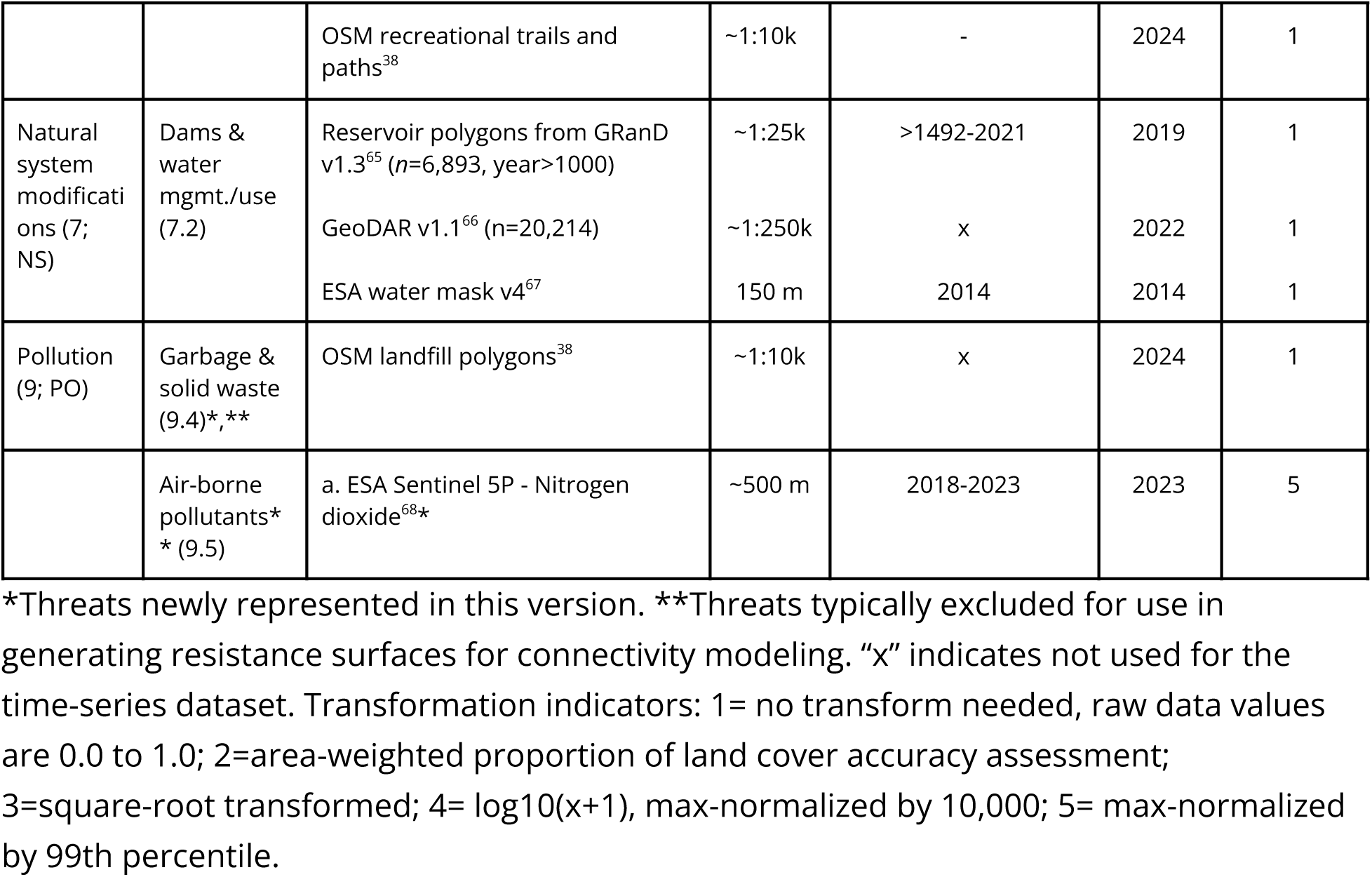
The classes and threats from the Direct Threats v2/IUCN Threats Classification v3.3 scheme, with related data sources used to create the global Human Modification datasets. This includes the spatial resolution, years of the data availability, the type of data about potential accuracy or uncertainty included, and the transformation formula to convert (i.e. normalize) the raw data values to range between 0.0-1.0. Threat classification levels in parentheses correspond to those within the Direct Threats Classification^25,26,27^, with abbreviations referencing threat class.

where *F_t_*is the proportion of a pixel occupied (i.e. the footprint) by threat *t*, and *I_t_* is the intensity – which is used to differentiate land uses that have varying impacts on terrestrial systems (e.g., grazing is less intensive than mining). Intensity values are based on a generalized land use coefficient, termed Landscape Development Intensity (LDI) that captures the per-unit amount of non-renewable energy required to maintain a human activity^30^, detailed for each threat in Table 2 and derivation of intensity values from LDI values (Supplementary Table 1). We note, however, that estimates of the intensity and negative impact are likely to be system- and region-specific, but with limited information to support such a tailored approach in a globally consistent way. However, the HM framework allows adjusting intensity weight and tailoring for different spatial regions based on regional expert input or from documented impacts where additional information is available and supported, particularly when downscaling to country or regional extents.

**Table 2.**
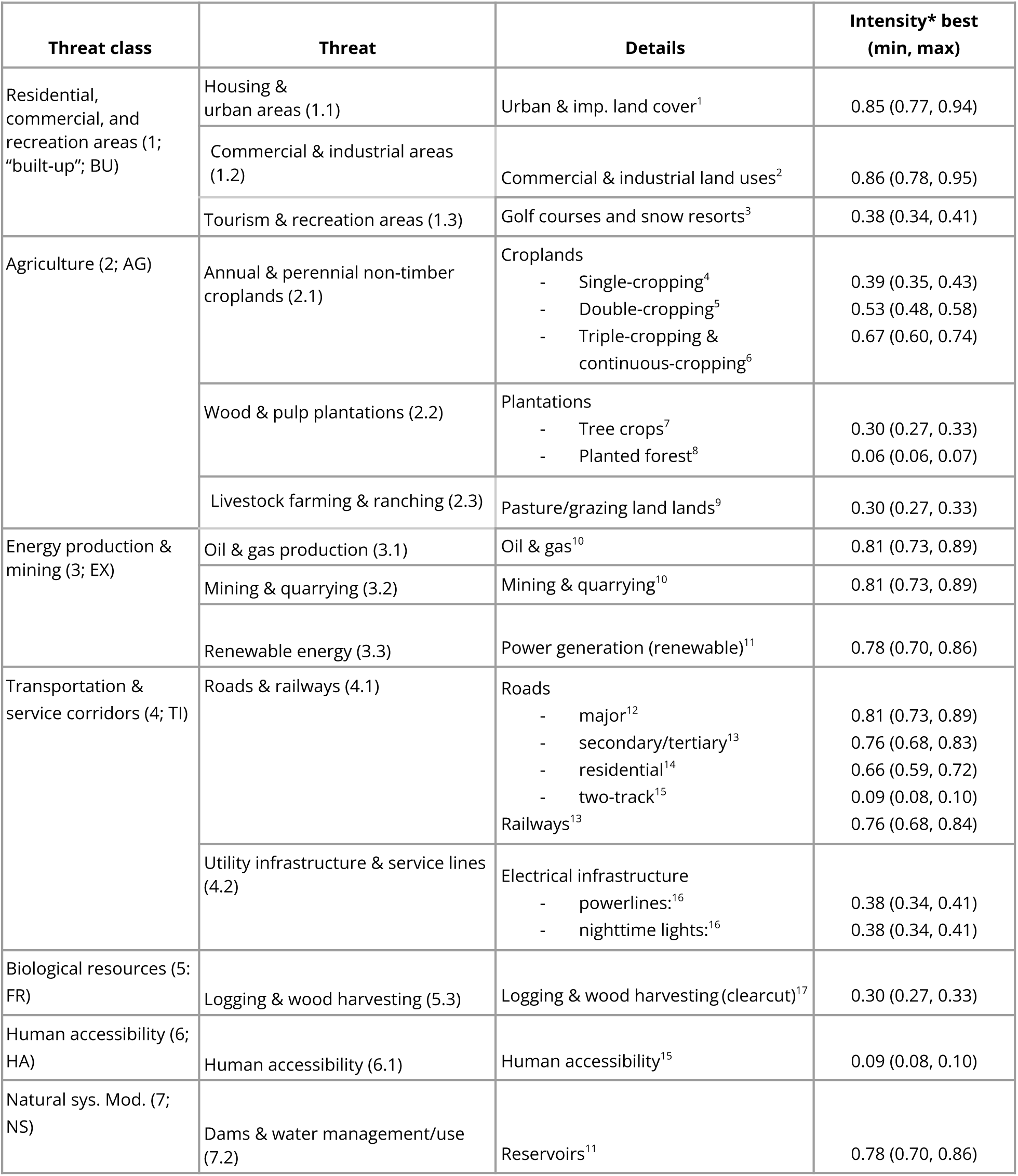
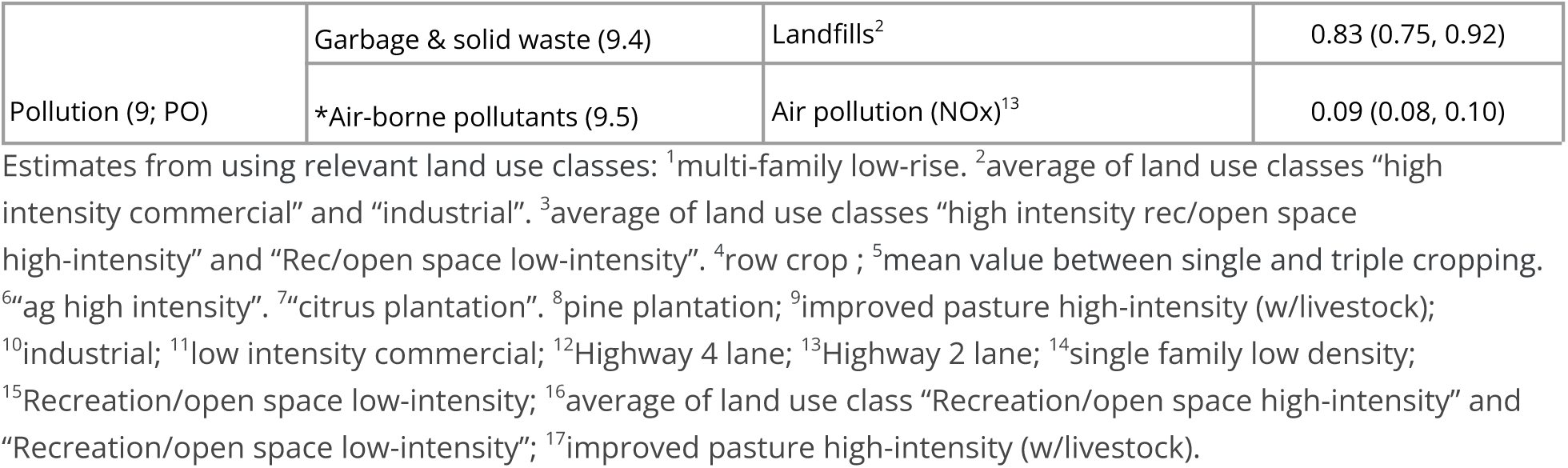
Estimated intensity values for each threat, organized by threat class. Intensity values shown are the “best” estimates and are bracketed by a minimum and maximum range, following the lowest-highest-best-estimate elicitation procedure to reduce bias^70^. Note: min and max values are specified as +/- 10% from best estimate, except when specific emergy-based range estimates were available^30,71^, see Supplementary Table 1.

To combine the degree of human modification for each individual threat *t*, we calculated *H* using a fuzzy sum^6,31,32^, (aka “increasive mean”):

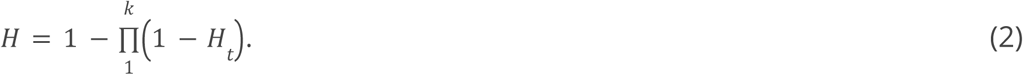

Because this formulation accounts for the partial correlation of multiple threats, it minimizes the bias associated with non-independent threats^6,28,32,33^, rather than implicitly assuming independence between threats made by simple additive calculations commonly used in human pressure modeling^7,17^. The increasive mean formula results in values of ratio data type (i.e., 0 to 1) that are less than the sum, equal to or greater than the maximum of any individual threat, and equals or exceeds the arithmetic mean.

To generate the human modification dataset, we reviewed for each threat readily-available global datasets and selected those that represented recent conditions (∼2015-2022), were mapped at the highest resolution possible (preferably ⪯300 m), provided multi-year data, and ideally included uncertainty estimates particularly for classification error, spatial precision, etc. That is, we selected datasets using the following criteria^34^: relevance and accuracy (e.g., does the dataset represent desired aspects in a clear way following standard definitions, datasets, and approaches and are robust accuracy and uncertainty statistics provided?); spatial scale (i.e. is the dataset at global extent and is it mapped at a high resolution (preferably <300 m) or even higher resolution?); temporal scale (e.g., can the data support spatially-explicit time-series datasets at time-step of 1-5 years, from 2000-2020 (at minimum), and current as well?); and licensing (e.g. are the data publicly accessible, available, and does licensing support open source publication and general use?).

We describe the specific datasets (Table 1), their data type and scale, and the equations used to calculate an updated measure of the degree of human modification globally from 1990 to 2022.

### Threat class 1: Residential, commercial, and recreation areas

We mapped the residential, commercial, and recreation area threats (i.e. built-up), typically associated with human settlement and impervious surface, commonly located in urban areas (Figure 2). We distinguished three primary built-up threats using OpenStreetMap land use polygons and identified areas dominated by: housing and residential land use (1.1), commercial and industrial (1.2), and concentrated tourism and recreation facilities (1.3). We used the built-up class from the ESA World Cover dataset^37^ (WC, 10 m resolution), the global human settlement layer^35^ (100 m;) and an impervious surface dataset Global Impervious Surface Area^36^ (GISA, 90 m) to identify the footprint associated with built-up land uses. The GHSL data provide information on structures only (buildings, etc.) while the WC and GISA maps also include non-building built-up areas, such as roads and parking lots. We combined the datasets by calculating the increasive mean to account for the partial dependency of the datasets within the threat, each weighted based on their classification accuracy.

**Figure 2.**
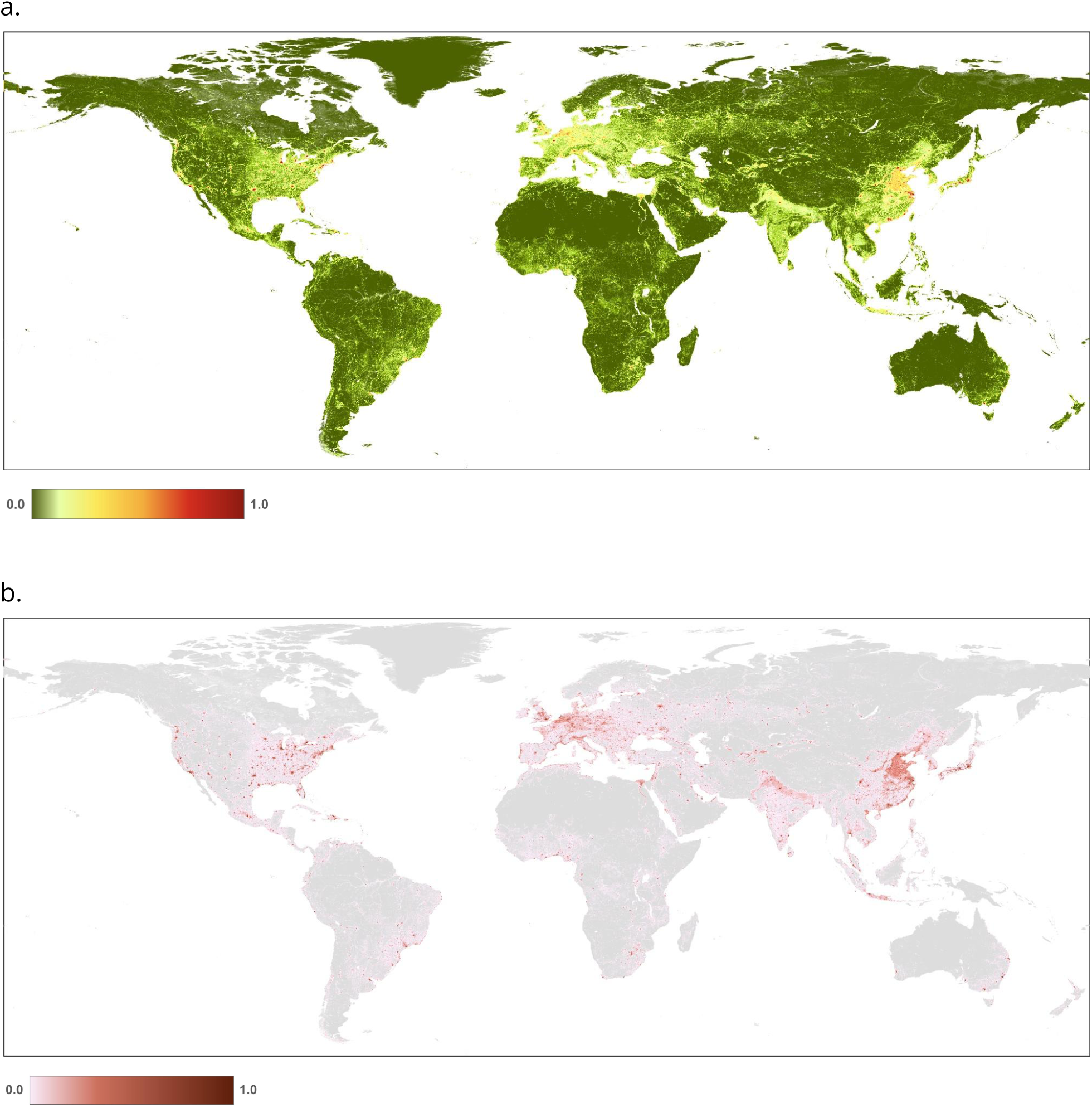
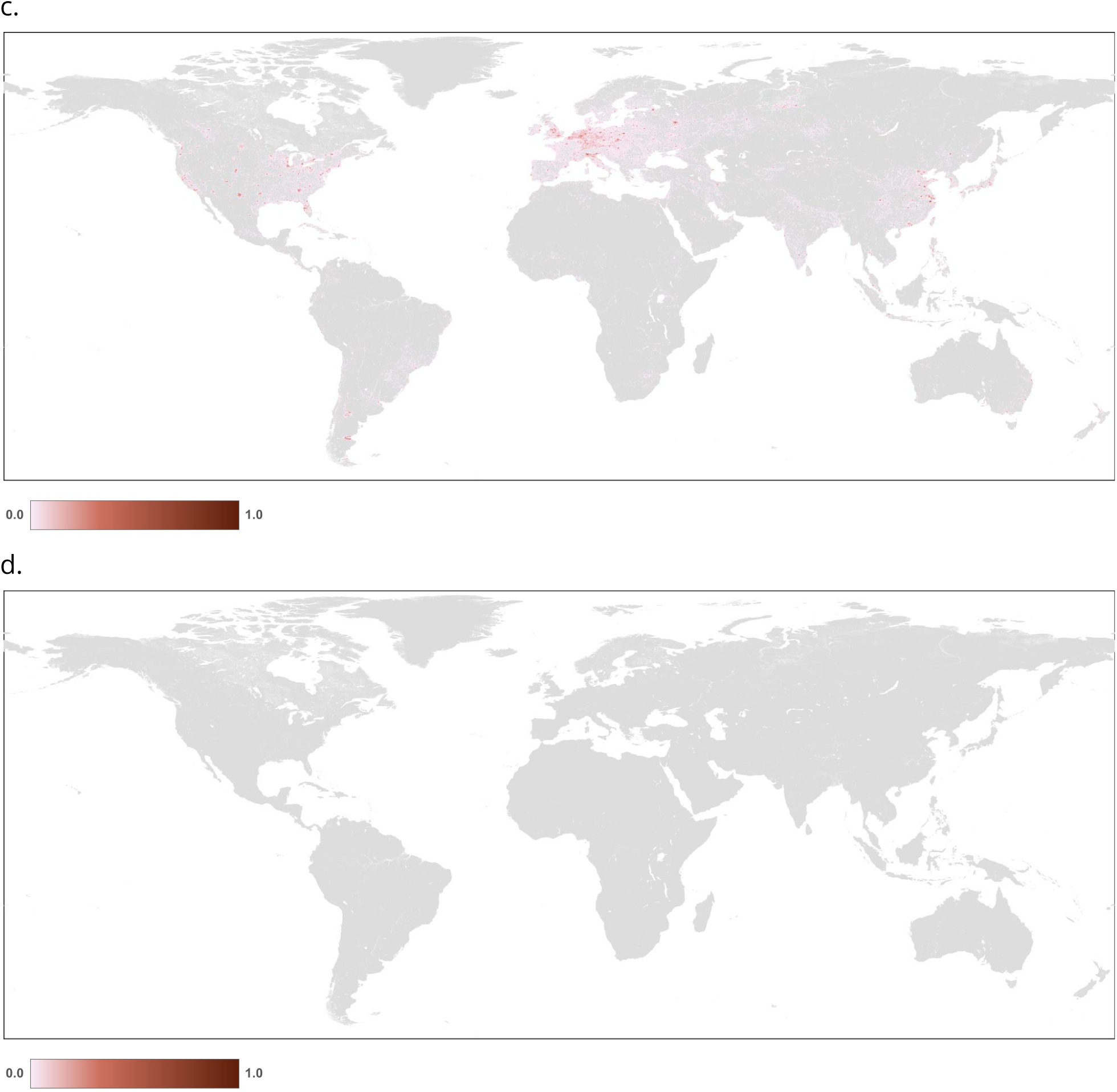
Maps of human modification values for: a. the residential & commercial development (built-up) threat class: b. housing & urban areas (1.1); c. commercial and industrial (1.2); and d. tourism and recreation areas (1.3).

We calculated the built-up footprint (*F_b_*) for circa 2022 (using the best and most recent datasets) as:

where:

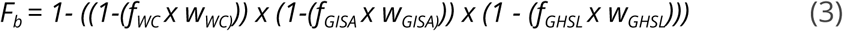

*f_WC_* = proportion of built-up land cover (aggregated to 90 m)

*f_GHSL_* = proportion of human settlement

*f_GISA_* = proportion of impervious surface

*w_WC_ = 0.9; w_GHSL_ = 0.8; w_GISA_ = 0.8*

For the datasets that support change analysis from 1990-2020, we calculated *F_b_*as:

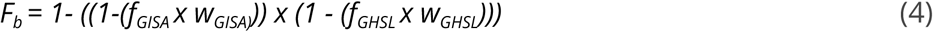

where:

*f_GHSL_* = proportion of human settlement

*f_GISA_* = proportion of impervious surface

*w_GHSL_ = 0.8; w_GISA_ = 0.8*

To calculate the footprint for the housing and residential threat (*F_br_*, 1.1), we excluded built-up areas identified by OSM polygons classed as commercial/industrial or recreation-tourism. To calculate the footprint for the commercial/industrial threat (*F_bc_*, 1.2), we included only built-up areas overlapped by OSM polygons (power = ‘plant’; *n*=38,475) and classed as commercial/industrial (1.2). Similarly, to calculate the footprint for the tourism/recreation threat (*F_bt_*, 1.2), we included only built-up areas overlapped by OSM polygons classed as tourism/recreation (1.3).

To calculate the human modification, *H*, for each threat, we multiplied the footprint by the intensity values specified in Table 2. For example, the human modification value for the housing/residential built-up threat is calculated as: *H_b_* = *F_b_* x *I_b_*. Also note that OSM datasets are only used when representing “current” conditions (for ∼2022), because OSM’s mapping process and data structure do not support temporal change analyses.

Note that we considered using other built-up datasets, especially temporally-consistent, but excluded them because they did not meet our spatial or temporal criteria and/or were direct replications of the more accepted datasets used in our analysis. We considered using the Microsoft building footprints^69^, but these data are already incorporated in the modeling of the GHSL. We considered but did not include, human population because we found that representing human use through the human intrusion/accessibility threat better represents potential impacts on ecosystems located nearby avenues of human movement (e.g., roads, trails) driven by accessibility (i.e. gravity models) to human populations rather than overly simple methods (i.e. buffering highways) that cause high rates of commission errors.

Commercial and industrial threats (1.2), although typically associated with urban areas, also occur in rural areas around land uses dominated by oil and gas infrastructure, factories and processing facilities, and non-renewable power production. To map these, we used OSM polygons (landuse = “commercial”’ n=492,224, “industrial”; *n*=1,717,542, or “retail”, *n*=337,811). We mapped non-renewable power production (i.e. power plants) using polygons from the OSM (power = ‘plant’; *n*=38,475) and points from the global power plant dataset (WRI; *n*=3,224). The WRI power plant dataset is used previous to 2020 because only for this timespan year built information is available.

We distinguished threats associated with recreation/tourism (1.3) from other built-up threats using OSM polygons that represent golf courses (landuse=”golf_course”, *n*=19,307) and winter resorts (where landuse=”winter sports”; *n*=3,055). We selected these land use types because they represent distinct areas of human activities (typically with boundaries and/or fences) and have negligible amounts of impervious surfaces and buildings within them that could have been captured within others threat maps. We examined but excluded polygons attributed with other OSM land use types that typically mapped very diffuse land uses represented by very large polygons (e.g., > ∼10 km^2^).

### Threat class 2: Agricultural

To account for impacts due to agricultural land uses, we mapped threats for non-timber croplands (2.1), wood & pulp plantations (2.2), and non-timber pasture/grazing (2.3). Note that activities associated with timber harvest (e.g., clear cut) are included in the biological resource use threat (see below).

#### Croplands

We calculated the the croplands footprint (*F_a_*) for ∼2022 conditions using the following datasets: croplands from Global Cropland Database^41^ (GCD; 30 m); extent of croplands from Global Cropland Intensity^42^ (GCI; 30 m); cropland class from land cover data ESA CCI 300 m^40^ (CCI; cover types: 10, 11, 12, and 20 at 300 m), and Global Cropland-Extent Product^43^ (GCEP30; 30 m).

We estimated the cropland footprint as the fractional cover of cropland within each output pixel (i.e. at 90 and 300 m), and excluded potential cropland pixels within urban areas (represented by a mask created CCI urban class buffered by 1 km) to minimize misclassifications (e.g., urban parks and golf courses).

Each of the input datasets represent croplands, but with slightly different purposes and satellite imagery source data. Therefore, we used a hybrid-fusion formula that accounts for datasets with partial dependency^72^, and combined them into the cropland threat using the increasing mean. We found the correlation (i.e. the proportion of overlap) of the GCD and GCI to be 0.7, and we decreased the weights to reflect the native resolution of each dataset where smaller pixels were given a higher weight. This approach leverages information from multiple data sources and represents the data from a probabilistic perspective. We determined the weights *w* by adjusting and comparing the cropland footprint to well-known and high-quality regional datasets: the Cropland Data Layer for the US^73^ and CORINE in Europe^74^, until the fit minimized the differences between these and the areal extent of our cropland footprint estimates (12% over estimation and 12% under-estimation, respectively). We confirmed that our estimate of the extent of the cropland footprint (1560 M ha) is within 1% of the FAO’s cropland estimate (1562 M ha). To address minor but widespread commission errors due to erroneous cropland pixels in the CCI dataset typically located in alpine tundra grasslands in high elevation mountainous landscapes, we removed cropland pixels from the GCD dataset that exceeded the 99th percentile of elevation for each continent (i.e. 2575 m for Africa, 2319 m for Euro-Asia-Oceania, 2365 m for North America, and 3407 m for South America).

We calculated (*F_ac_*) reflecting current conditions (∼2022) as:

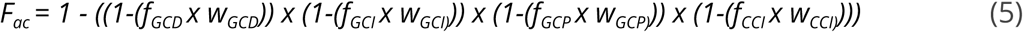

where:

*f_GCD_* = proportion of cropland pixels (30 m) in each footprint pixel (90, 300 m)

*f_GCP_* = proportion cropland (30 m) within a 300 m pixel

*f_CCI_* = proportion of cropland class from land cover pixels (300 m)

*f_GCI_* = proportion cropland intensity classes (30 m) used as an index to intensity estimate

*w_GCD_ = 0.5; w_GCP_ = 0.45; w_CCI_ = 0.35*

The cropland footprint (*F_ac_*) for the time-series datasets (1990-2020) was calculated as:

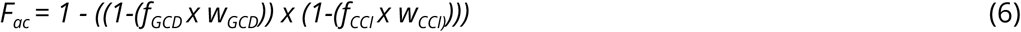

where:

*f_GCD_*, *f_GCI_* = proportion of cropland pixels (30 m) in each footprint pixel (300 m)

*f_CCI_* = proportion of cropland class land cover pixels (300 m) in each footprint pixel (300 m)

*w_GCD_ = 0.9; w_CCI_ = 0.5*.

To calculate *H_ac_*for cropland (Figure 3), we multiplied the cropland footprint value times the intensity values times: *H_ac_*= *F_ac_* x *I_ac_*. We determined spatially-explicit cropland *intensity* values by considering factors described in literature on the frequency of cultivation and amount of external inputs factors that cause impacts due to cropland systems^75,76^. These factors commonly relate to the number of harvest rotations, irrigation source, tillage, and chemical inputs (fertilizer, herbicides, etc.). We chose to represent intensity values based on harvest rotations because it is a primary factor that tends to be less dynamic inter-annually and because high resolution datasets are readily available and with coverage globally. To do this, we used the GCI dataset that represents single, double, and triple (and continuous) cropping and applied a specific estimated intensity value for each (see Supplementary Table 2). We filled *no data* values in the GCI dataset that appeared as latitudinal “cracks” using a modal filter with a 20 km moving window, then smoothed by a 2 pixel radius moving window (∼20 km) to reduce artifacts introduced by the relatively coarse resolution. We calculated *H_c_* for the cropland threat as the footprint times the intensity value associated with each cropland rotation class at each pixel.

**Figure 3.**
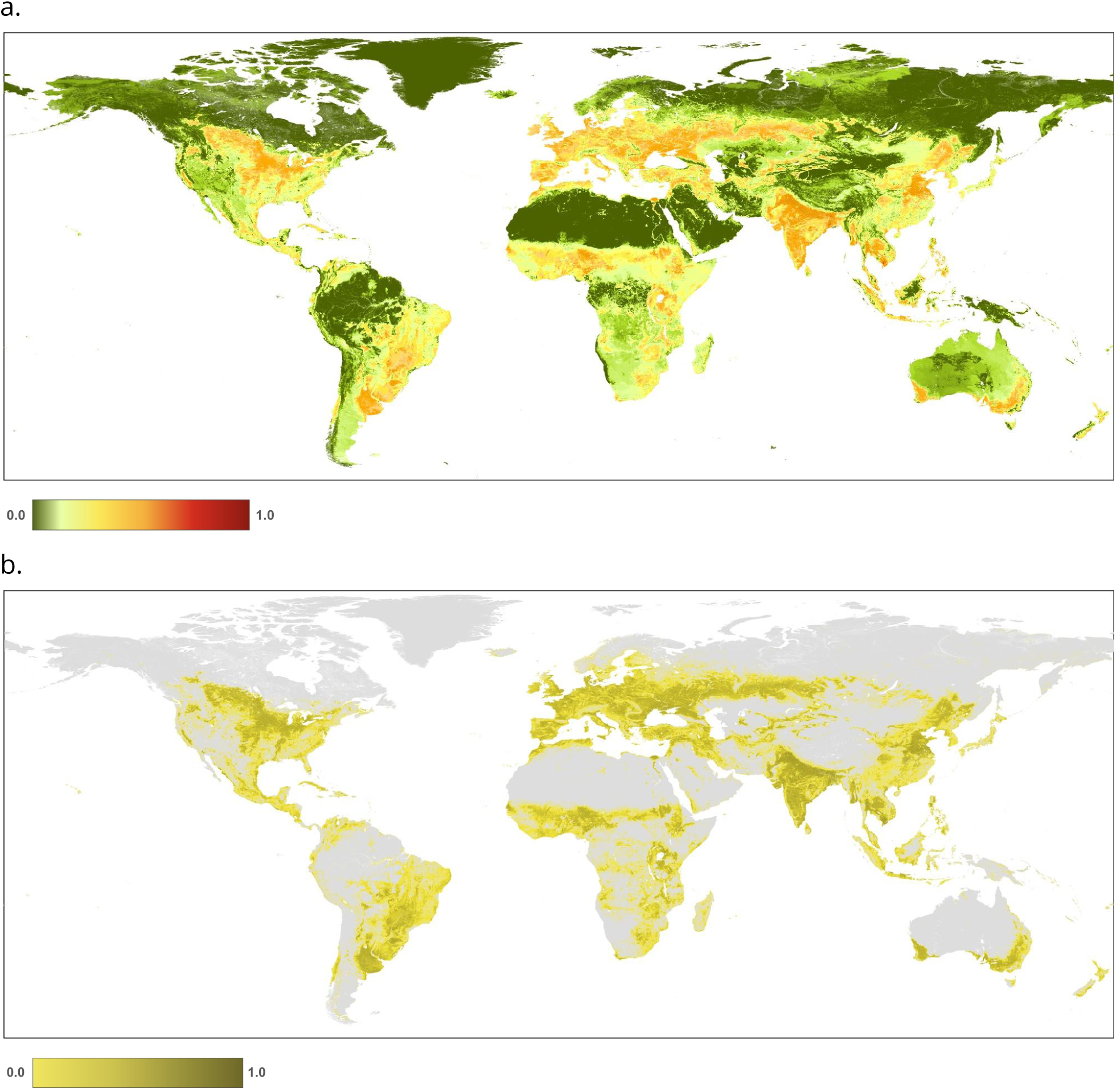
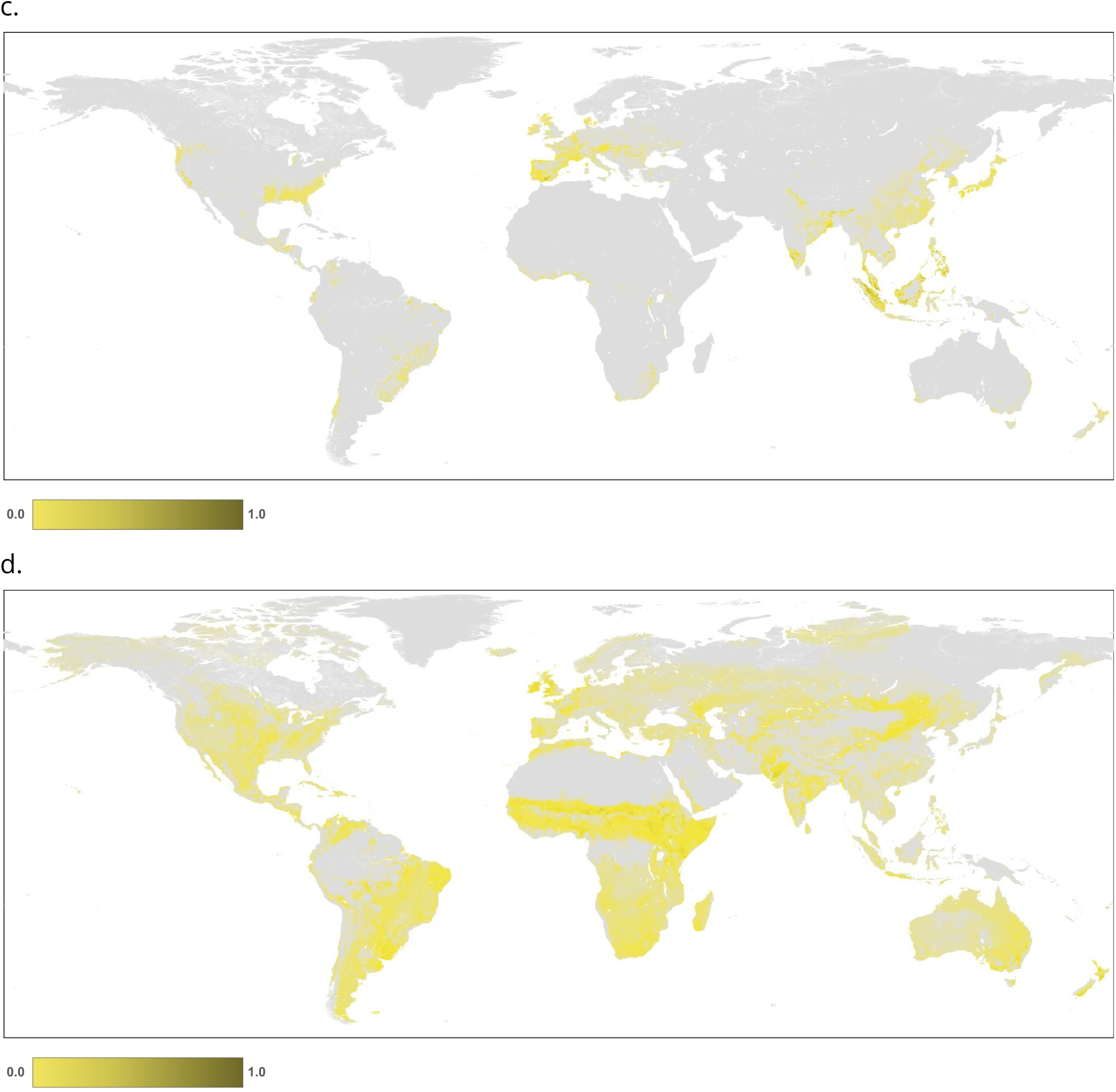
Maps of human modification values for: a. agriculture threat class; b. Annual & perennial non-timber croplands (2.1); c. wood & pulp plantations (2.2); and d. Livestock farming & ranching (2.3).

#### Wood & pulp plantations

The footprint of wood and pulp plantations was determined using data^44^ that maps planting years globally at 30 m from 1984 to 2020. We mapped the plantation footprint as those pixels currently classified as a plantation (i.e. we did not distinguish the number of years prior to the current year, which would require assumptions about the accumulation of effects over time), multiplied times the per-pixel proportion of forest canopy cover in 2010^77^.

To calculate the HM intensity values for plantations, we grouped the 190 species/systems into two basic types: stands of perennial *tree crops* (e.g., oil palm, coffee, orchards) vs. *planted forests* grown for wood, fiber, or sheltering from erosive processes). We then used estimated intensity values for wood & pulp plantations (Table 2) to calculate *H_ap_* for plantations.

#### Pasture/grazing lands

To calculate the pasture/grazing lands *footprint*, we used two datasets: (a) land cover data from ESA CCI using the mosaic-croplands classes and other potential pasture/grazing types, including mosaic herbaceous, shrublands, grassland, sparse shrub and grass, sparse forest, and shrub/herbaceous flooded types (note cropland is excluded); and (b) livestock grazing densities Gridded Livestock of the World v4 for 2015^46–51^.

We selected these datasets because they provide the best available land cover data representing possible cover types consistent with pasture/grazing lands and mapped at multiple time steps. We calculated the pasture/grazing lands footprint by first mapping land cover types consistent with grazing, and then adjusting by livestock density estimates For this, we use the CCI land cover types, that were biophysically were very likely to be providing forage resources (i.e., excluding types like water, snow/ice, barren)^78^. We added sparse grasslands, shrublands, and forest pixels to this footprint where grazing densities were within the 95% distribution of grazing density values of grazed land cover classes (Supplementary Table 2). We spatially “sharpened” the livestock density map by calculating the mean value for each contiguous region (i.e. “patch”) found within each land cover type, with pixels intersecting major roads (highways) removed, and adjusted the footprint to account for livestock grazing density (as suggested by^18^). Livestock density was calculated by establishing a maximum threshold of 1000 livestock units/km^2^ based on the break-point between grazing and industrial livestock systems^79^. We applied a square-root transform of the density values rather than a log10(x+1) used in ^33^, following detailed inspection and feedback from expert reviewers to reflect presumed lower footprint values.

The livestock farming & ranching class (pasture/grazing) land footprint (*F_p_*) was calculated as:

where:

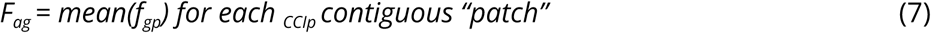

*f_CCIp_* = the region for each pastureland class land cover defined by 8 neighbors; *f_gp_*= the footprint of the livestock density, calculated as the animal unit density (au/km^2^) of bovines, goats, and sheep, and then normalized by 1000 then square-root transformed to minimize the effects of outliers.

To calculate *H_ag_*, we multiplied the pasture/grazing footprint value times the intensity values times: *H_ag_* = *F_ag_* x *I_ag_*. Note that we explored but did not use the Farming the Planet v1 pasture datasets^18^, because it focuses more on pasture than grazing, was relatively coarse resolution (we used higher resolution land cover data and livestock data – both primary inputs to the FTP modeled dataset), was static and represented relatively old conditions (year 2000), and had strong artifacts of abrupt edges associated with census data compiled within various administrative boundaries. We also considered but did not use the approach by Piipponen et al. (2022), because it focussed on potential net productivity and represented only 2010 and 2015 conditions.

We calculated the *H* values for the agricultural threat class as the pixel-wise maximum of *H* from the datasets for cropland, plantations, and pasture/grazing land: *H_a_*=max(*H_ac_*, *H_ap_*, *H_ag_*), because cropland, pasture/grazing lands, and plantation footprints may have some spatial overlap, in particular as a consequence of combining different resolution datasets.

To calculate *H_a_*(Figure 3), we multiplied the footprint value times the intensity value.

### Threat class 3: Energy production & mining

The primary component threats in the energy production & mining threat class are oil & gas extraction (3.1), mining & quarrying; (3.2), and renewable-energy (power plant production; 3.3).

#### Oil & gas extraction

We used three data sources to estimate the human modification associated with oil & gas extraction *H_eo_*. Nighttime flare locations are derived from DMSP and VIIRS nighttime light datasets and 90% of gas flares occur at locations where oil and gas are extracted^53^. For each flare, we thus approximated a footprint of 0.057 km^2^ per well-head^80^ using a cone with a linear decay to 0 at a radius of 233 m. Because the infrastructure needed associated with gas flares would remain on the landscape after de-commisioning (e.g., as plugged wells, well pads, access roads, etc.), we assumed that the footprint of these threats could only increase over time. These data on nighttime gas flares were used for both the 1990-2020 time-series and for ∼2022. We also mapped oil and gas wells using data from recent OSM data (downloaded July 16, 2024, using points with the attribute man_made = ‘oil_well’, ‘gas_well’, or ‘petroleum_well’; *n*=2,915,936), and from the Oil and Gas Infrastructure Mapping database^54^ (*n*=2,560,399). We considered mapping pipelines as well, but did not include these features because they were generally drawn too spatially imprecise and lacked completeness in the OSM data.

#### Mining & quarrying

To estimate the human modification associated with mining & quarrying, *H_em_* we combined polygon data from 3 datasets: OSM landuse (‘quarry’, *n*=203,715), ‘industrial’, *n*=1,224,318), the global mining database v2^55^ (*n*=∼44,000), and from global mining areas^56^ (*n*=∼74,548). These data were used for just the HM dataset representing 2022 conditions.

#### Renewable energy

To estimate the human modification associated with renewable-energy production *H_er_*, we mapped features from the global power plant dataset (WRI; *n*=23,364), solar energy facilities^57^ (polygons *n*=63,616), and wind turbine locations from OSM (man_made = ‘windmill’; *n*=6,612) with a footprint of 0.0014 km^2^ from ^81^. These data were used for the ∼2022 dataset, and the WRI power plant dataset was used as well for the 1990-2020 time-series.

To calculate *H_e_*(Figure 4), we multiplied the footprint value times the intensity value.

**Figure 4.**
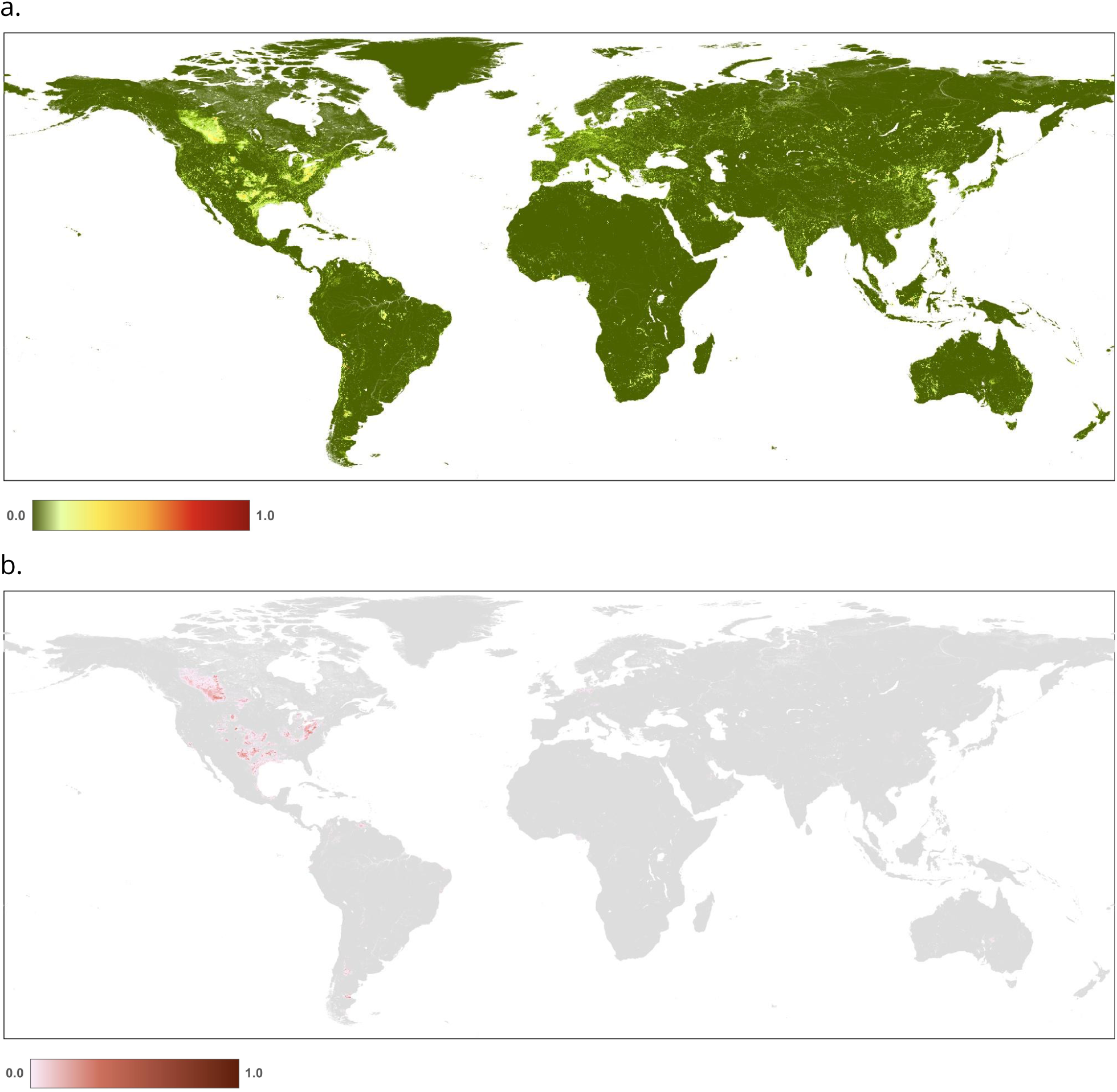
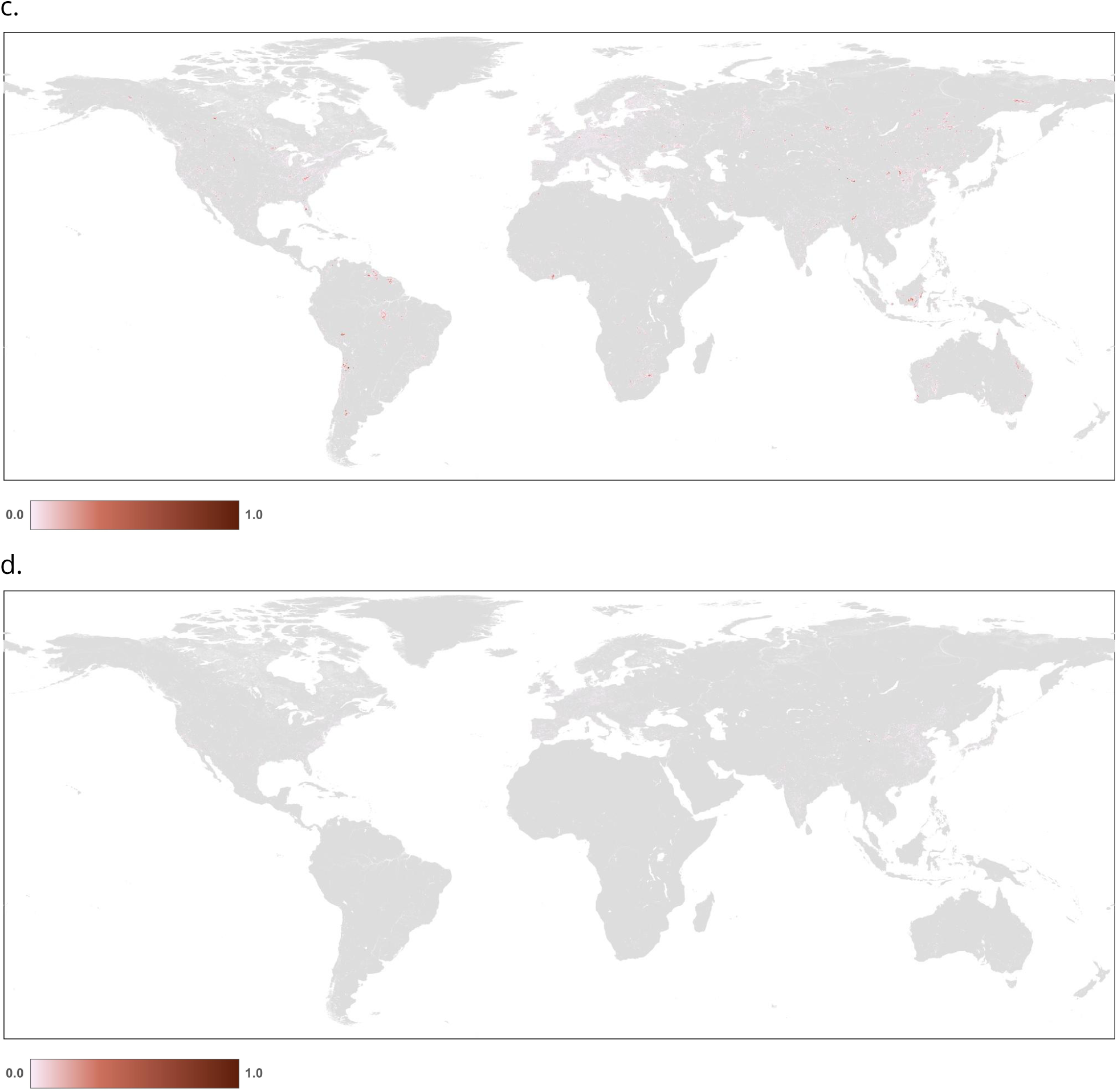
Maps of human modification values for: a. energy production & mining threat class; b. oil & gas production (3.1); c. mining & quarrying (3.2); and d. renewable energy production (3.3). (Note: because the location of threats in c & d are limited in area, they are difficult to see on a global extent).

### Threat class 4: Transportation & service corridors

We mapped two threats associated with the transportation & service corridors class (4), including roads & railways threat (4.1) and utility infrastructure & service lines (4.2).

#### Roads & railways

We mapped the footprint of roads and railways as the fractional cover of a pixel^10^ occupied by the road-way, the road bed, and any median if present. Road and railway features from <u>OSM</u> types were differentiated into four major types that have separate HM footprint and intensities ascribed: highways (*f_t1_*) included “motorway”, “trunk”, “primary”, “motorway_link”, or “primary_link”; secondary/tertiary (*f_t2_*) included “secondary”, “tertiary”, “secondary_link”, “tertiary_link”; residential (*f_t3_*) included “residential,” “road”, or “street”); two-track (*f_t4_*) included “track”, “unclassified”, or “service usage”; and railways (*f_t5_*) included “rail”, “narrow_gauge”, “light_rail”, “abandoned”, “disused”, or “tram”). To remove errors of commission caused by roads mis-attributed as residential, but likely to be “tracks” because of their remote locations, we restricted residential roads to be within ∼5 km of settlement locations (specified by GHSL settlement data used in the built-up threat class).

For the transportation footprints, the width was estimated to include the road surface and immediately adjacent land where cover is significantly modified and associated with the road and was calculated by dividing the per-road type estimate of road width (m) by the pixel width (e.g., 100 m). That is, the footprint of a highway (approximately 40 m wide) at 90 m resolution would be 4/9 or 0.45, not 1.0 if calculated simply as a road touching a pixel. The estimated widths (*w*) were 40, 20, 15, 5, and 20 m (highways, secondary, residential, track, rail). Note that the footprint is calculated for both lines of a divided highway, if represented by distinct lines. Because pixels that have different road types intersect will have different footprint values, we added the transportation-class-specific H values at each pixel.

The transportation footprint (*F_tr_*) was calculated as:

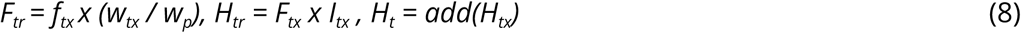

where *tx* indicates the type of transportation features (highways, secondary, residential, track, and rail, respectively) and *w_1_=40, w_2_=20, w_3_=15, w_4_=5, w_5_=20, w_p_=300*.

To calculate *H_tr_* for transportation threats (Figure 5), we multiplied the footprint value times the intensity values. Note that the human modification approach purposely does not represent presumed ecological “edge-effects” – in particular those associated with roads that are typically modeled with simple buffers extending up to 15 km (e.g. ^7^) and applied uniformly to both urban and remote sections of roads. Instead, we model adjacent/nearby “use” effects directly in the human accessibility threat, because human activities and land use depend on numerous factors other than the physical footprint, including the traffic volume, proximity to and population of nearby cities and towns, and off-road/trail characteristics that influence access and human activities such as land cover type, topography, and altitude^82,83^. This allows the human modification dataset to be used subsequently in modeling, including intactness, species distribution and use, and connectivity) to apply the desired spatial process on the HM dataset without confounding effects. Data on just highways and railways for 2024 were used in the time-series dataset to represent the core transportation infrastructure, but data on all roads was included in just the ∼2022 dataset.

**Figure 5.**
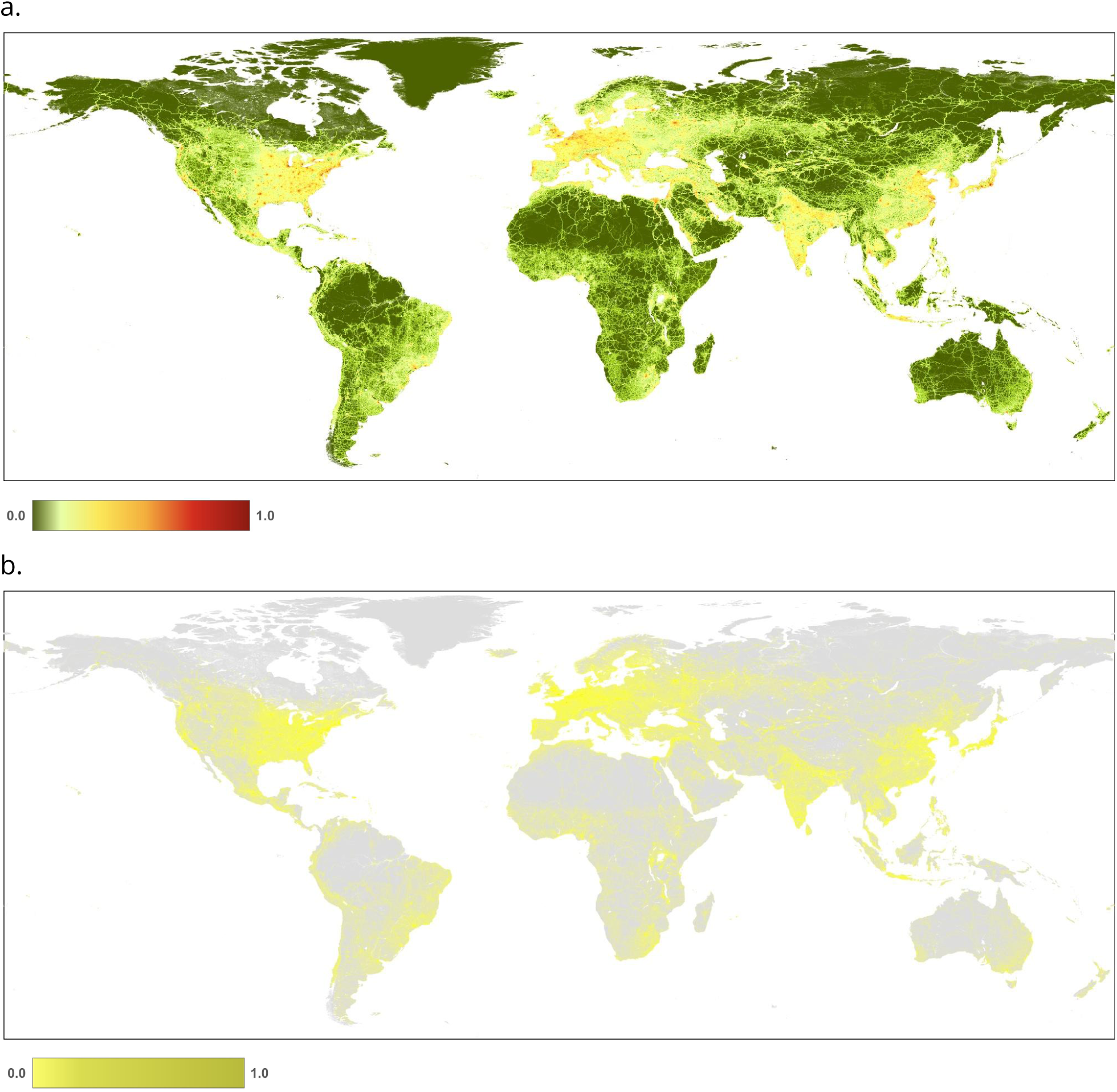
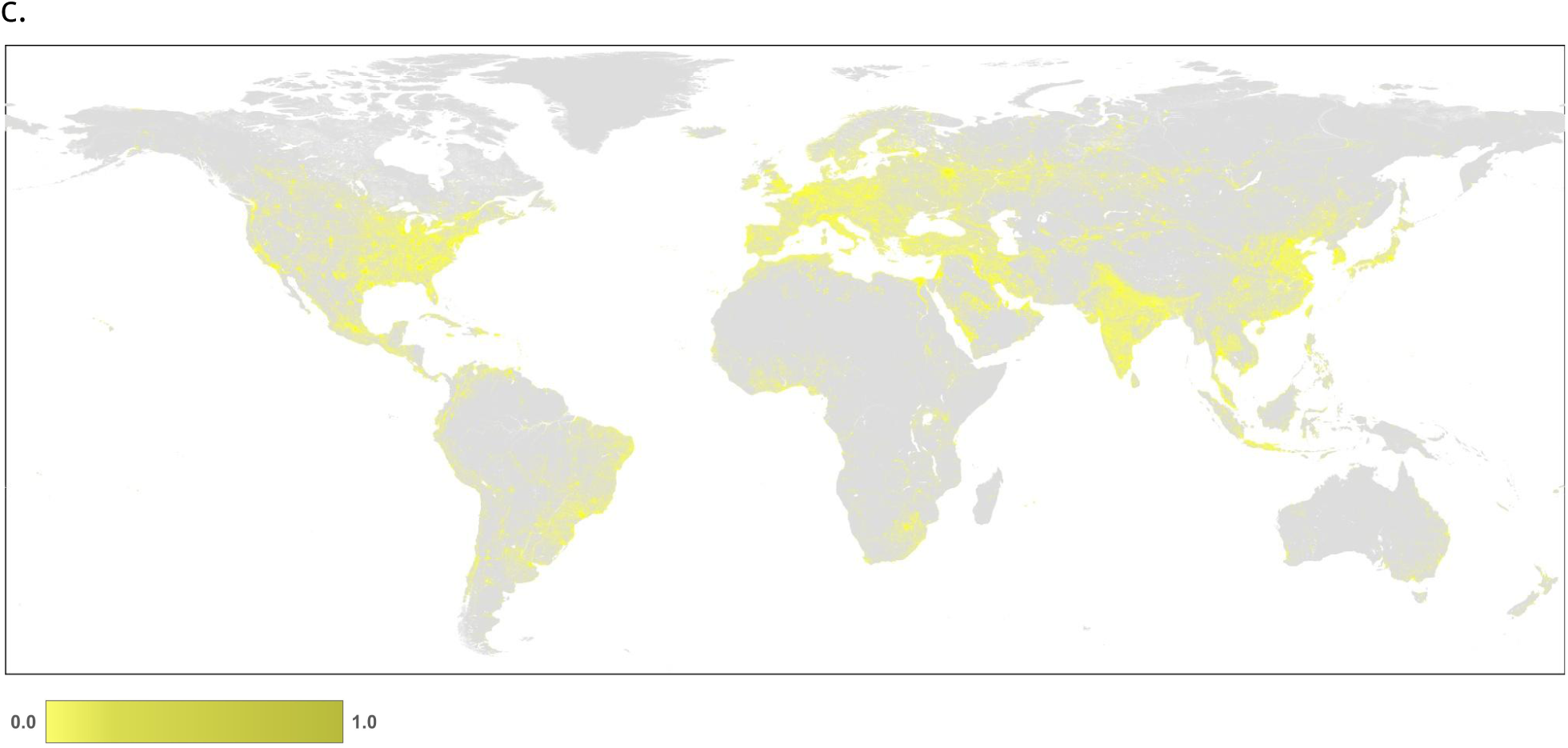
Maps of human modification values for: a. Transportation & service corridors threat class; b. roads & railways (4.1); and c. utility infrastructure and service lines (4.2).

#### Utility infrastructure

To map threats associated with utility infrastructure & service lines *H_tl_*, we used above-ground power lines and annual-average nighttime lights data. For powerlines, we compiled data from OSM linear features (power = “line”; *n*=968,651), and calculated the footprint using a kernel width of 250 m, max-normalized by 0.25 to reflect spatial uncertainty of the data. For nightlights data for the HM time-series datasets (1990-2020), we used a dataset^61,62^ (DVNL) developed specifically for harmonizing trends across the DMSP (1993-2013) and VIIRS (2013-2023) satellite platforms. DVNL is estimated as a “DMSP-like” nightlight value that leverages VIIRS satellite data for 2013-2019. We assumed that nightlights could only increase over time because effects of infrastructure associated with activities that were captured by the nightlights data typically remain present. Because the VIIRS data reduced the “blooming” artifacts present in the DMSP data^10^, we constrained the 1992-2013 series of DMSP using the DVNL 2014 maximum value (note that 2013 which had an error in northerly latitudes). The footprint is calculated as max-normalized by the maximum value reported in DMSP (x=63). For the HM product for 2020-2022, we used the raw VIIRS datasets, add 1 and log_10_ transformed, and then max-normalized by the 95^th^ percentile globally for 2022 (1.05). Data on powerlines mapped in 2024 were used just for the ∼2022 dataset, but data on nighttime lights was used in the time-series datasets as well.

To calculate *H_t_*(Figure 5), we multiplied the footprint value times the intensity value.

### Threat class 5: Biological resource use

To represent the biological resource use threat, we focused on the forest driver^84^ “intensive deforestation loss” (e.g., clear-cutting). Specifically, the footprint of timber harvesting was calculated as those pixels that had been designated as a) deforested in any year prior to the mapping year from the global forest loss dataset^63^; b) had been at least 10 m in height in 2000 to limit possible errors of commission of loss^45^; and c) was not deforested in the year that a wildfire occurred^64^. We simplified the calculation to ignore forest regrowth, because data were available only since 2001 and because of the broad variation of growth rates. We recognize that tree height does not necessarily account for the structural complexity, species diversity, biomass, etc. of forest ecosystems in general, but it is a practical surrogate for potential natural recovery and is conservatively estimated. We note that if deforested pixel was converted to another land cover/use (e.g., cropland or pasture/grazing), then the footprint will also be mapped for the resulting threat.

The biological resource use (timber harvest) footprint (*F_f_*) was calculated as:

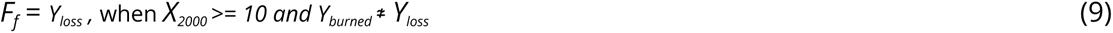

*X_2000_* = forest height (m) in 2000

*Y_loss_* = year forest lost

*Y_burned_* = year a pixel was burned by wildfire

To calculate *H_f_*(Figure 6), we multiplied the footprint value times the intensity value.

**Figure 6.**
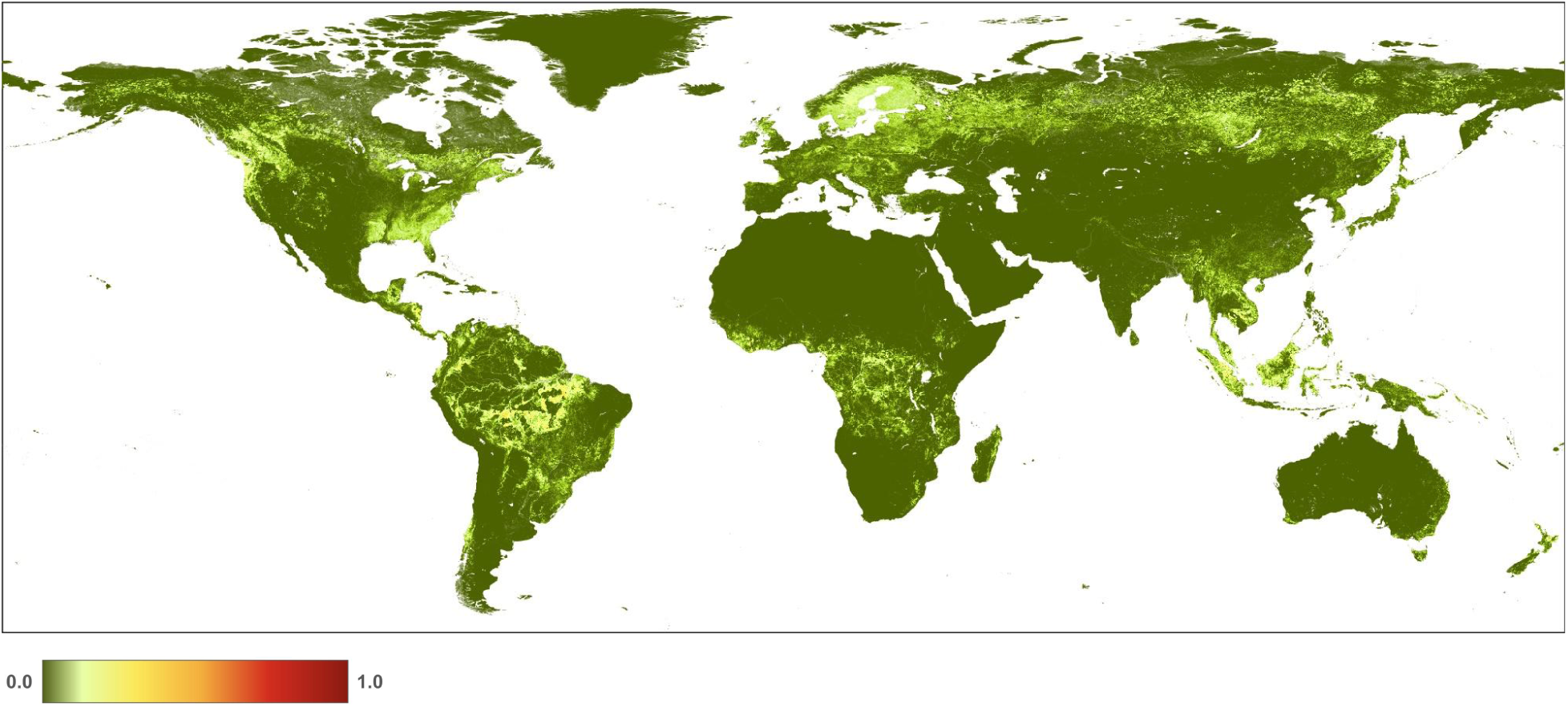
Map of biological resource use (timber harvesting) threat class (5.3).

We examined datasets and ways to map *selective* harvesting (another major driver of deforestation^84^), but did not include it as a threat because selective harvesting occurs at very fine spatial scales (<10 m) and locations with “opened” canopy in tropical regions often closes quickly (<1 year) so these areas are challenging to monitor in annual datasets. RADD datasets^85^ offer promising potential to map this type of deforestation, though a noted disadvantage is that these data are only available since 2020. Measuring very fine-scale (∼tree level) disturbances is challenging, especially for those locations where it is an incursion into unconverted forest. Deforestation due to annual & perennial non-forest crops are estimated in the agricultural threat; areas of forest loss due to wildfire are excluded, but note that loss due to other natural disturbances such as insect, disease, and hurricanes are not included because data that map these are not readily available, and forest loss due to urbanization is captured in the built-up threat. Data on selective harvest of forest resources was used in the time-series from 2005 to 2020, as well as in the ∼2022 dataset.

### Threat class 6: Human accessibility

Human activities (termed “intrusions” in the threat taxonomy) themselves are an important threat on biological diversity that include various human activities and uses that do not result in land cover transformation, yet are associated with or are surrogates for a variety of ecological impacts^86^. These activities include: use of roads (traffic) and associated noise, lights, and other types of pollution, activities adjacent to roads and trails such as collecting/foraging for plants and animals or use for “backcountry” recreation, or surrogates for invasive species.

To represent the human accessibility threat (6), we estimate the threat associated with human activities using accessibility modeling and human populations based on central place theory implemented using a gravity model^82^. Notably, we explicitly estimate both the amount of pressure associated with human activities using a surrogate of the population of towns and cities, and the accessibility based on estimated travel time, both on transportation infrastructure, such as roads and on all off-road lands and waters (including rivers, lakes and coastlines). This approach provides significant improvements to overly simple geospatial methods (i.e. uniform buffering of roads and approximating continuous processes like distance decay with arbitrary classes) that ignore basic human geography and landscape ecology principles.

Accessibility is estimated as a function of a “resistance” surface, where each pixel is assigned a travel time or “cost” to traverse. Different costs or weights are assigned depending on road types for off-road on slope and dominant land cover types, including rivers, lakes, and coastlines^10^. Accessibility was calculated using estimates of travel time along roads and rails, as well as off-road through different features of the landscape, using established travel time factors^87–89^: highways=19, secondary=48, residential=96, track=167, rail=169, trails=200, and water=167 (minutes per km). This included incorporating trails from OSM, effects of international borders following ^89^, and accessibility to lands from travel across water bodies (including coastlines).

Travel time is calculated in minutes away from locations with concentrations of human populations (i.e. urban areas, towns, villages, etc.) across the resistance surface that estimates travel time. Data on population density (people per km^2^) were obtained from the ESA’s Global Human Settlement POP dataset^90^ (100 m resolution). We identified population density classes (*C*) of development as the integer values obtained from a natural-log transformation of the mean population density (*D_2000m_*) within a 2 km Gaussian kernel, resulting in 15 levels. That is:

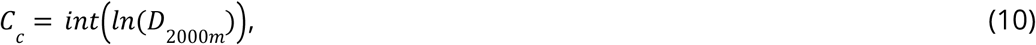

when *D_2000m_* > 1.0. The population (number of people) was estimated within each population density “patch” defined from the contiguous pixels for each class *c*. The estimated population was also obtained as the integer values (*z*) of *ln*() transformed population estimates. For each population class *z*, travel time (*T*) is calculated using least-cost distance calculations using the accessibility surface as cost-weights, from all pixels of class *z,* and assumed to decline by half for each 30 minutes traveled. Population density (i.e. the number of people assumed to travel) is used to weight the accessibility layers. The resulting estimates were then summed per-pixel for all density classes to estimate the total population that could access any pixel:

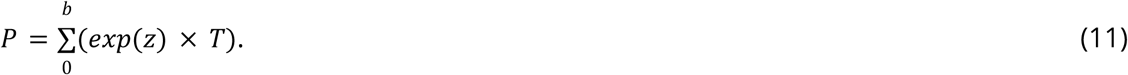

To calculate the footprint of human accessibility (*F_h_*), we normalized *P* by its value at the ∼90th percentile – roughly approximating the equivalent of a town of 100,000 people:

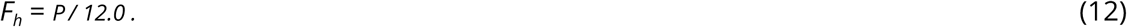

To represent increased footprints associated with human accessibility along and nearby trails, we added the footprint from representing trails mapped in the OSM dataset (highway=”path”; n-8,476,902) with a linear decay to 0 at a radius of 233 m.

Year-specific data on human accessibility were used for both the time-series dataset, and the current ∼2022 dataset. To calculate *H_h_*for the human accessibility threat (Figure 7), we multiplied the footprint value times the intensity values.

**Figure 7.**
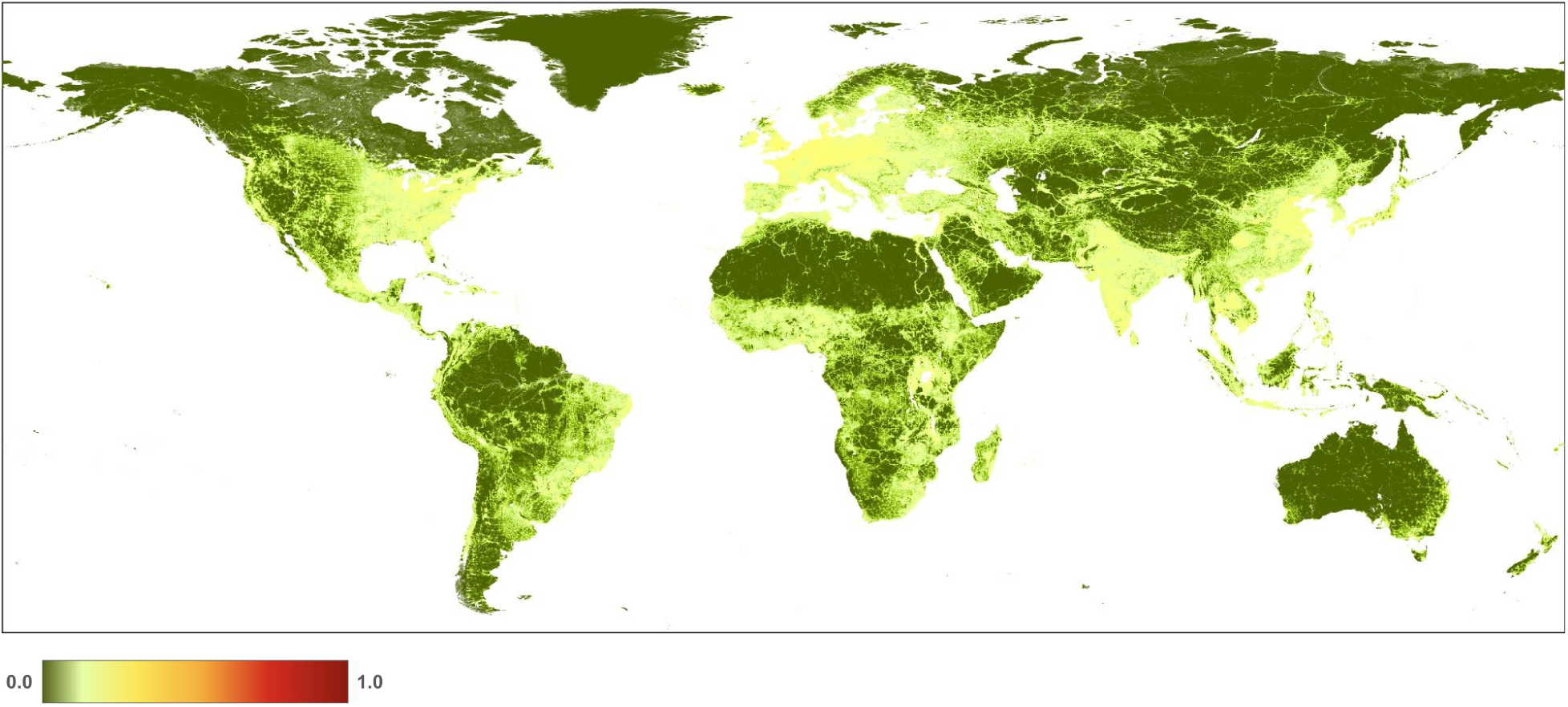
Maps of human modification values for the human accessibility threat class and threat (6).

### Threat class 7: Natural system modification

Dams and their associated reservoirs flood landscapes and strongly modify natural flow regimes of the adjacent rivers^91^. To represent natural system modification (albeit associated only with dams/reservoirs) threat, we mapped the footprint of reservoirs from the GRanD v1.3^65^ for the change datasets and GeoDAR v1.1^66^ for the 2020-2022 output. We buffered the reservoir polygons by 1 km and then masked them using the ESA watermask v4^67^ to reduce artifacts at the edges caused by varying water levels and different scale representations of water bodies. We did not include Lake Victoria (Africa) as a reservoir based on expert regional feedback. Data on reservoirs were used in both the time-series and ∼2022 datasets.

To calculate *H_n_*for the natural system modification threat (Figure 8), we multiplied the footprint value times the intensity values.

**Figure 8.**
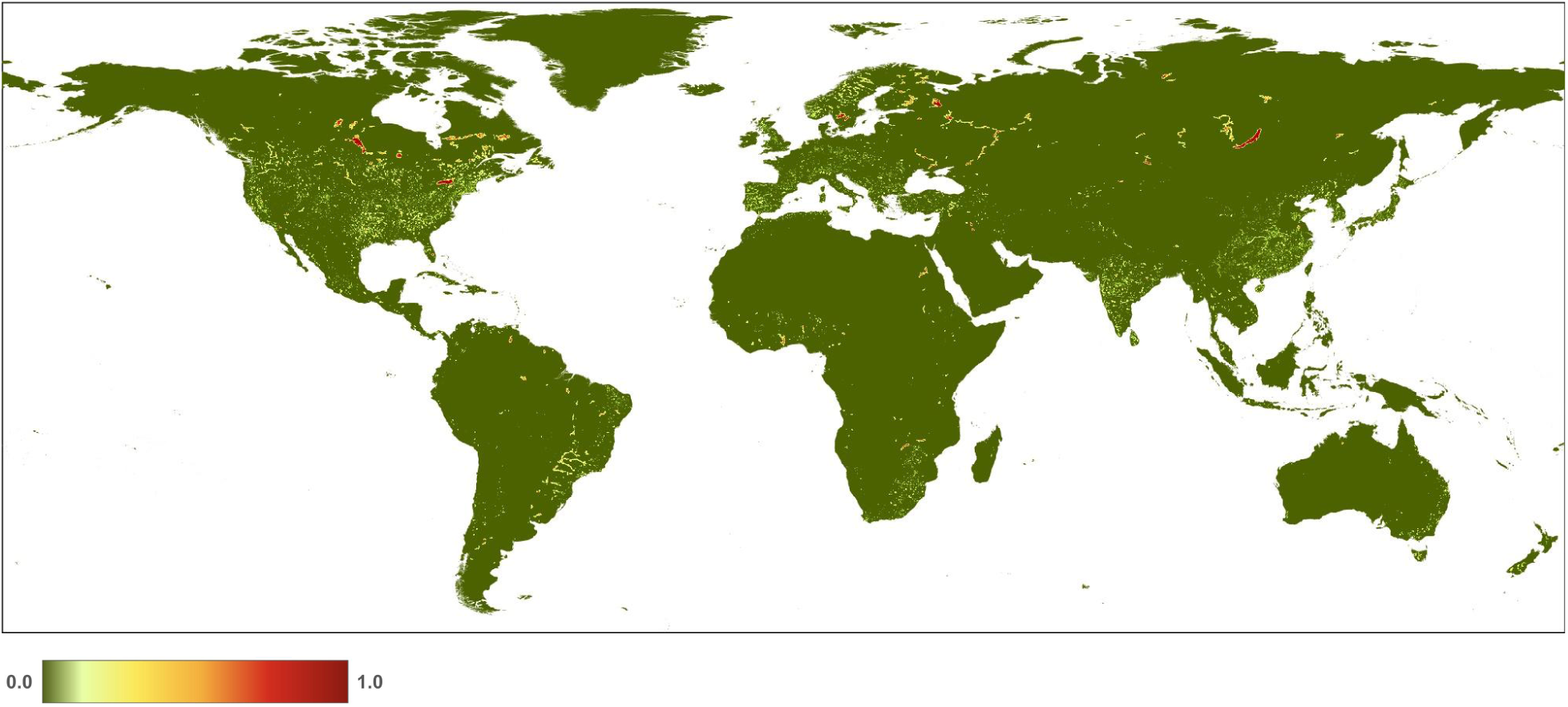
Map of human modification values for the natural system modification (focusing on dams and reservoirs) threat class and threat (7.2).

### Threat class 9: Pollution

To map garbage and solid-waste (i.e. landfills) and associated pollutants, we used OSM polygons (landuse=”landfills”, *n*=53,236). We mapped pollution using datasets of nitrous oxides because atmospheric levels of NO_x_ are a strong contributor to acid rain and fog and tropospheric ozone^92^ with impacts to biodiversity^93,94^, and is primarily generated by human land use activities. We estimated the footprint of air pollution by using data on nitrogen dioxide (NO_2_) from the Sentinel-5P satellite data collection (TROPOMI 2023, “tropospheric NO2 column number density”). We max-normalized using the 99th percentile value (0.0003). Data on garbage/solid waste was used only in the ∼2022 dataset, while air pollution was used in both the time-series and ∼2022 datasets.

To calculate *H* for the pollution threat (Figure 9), we multiplied the footprint value times the intensity value.

**Figure 9.**
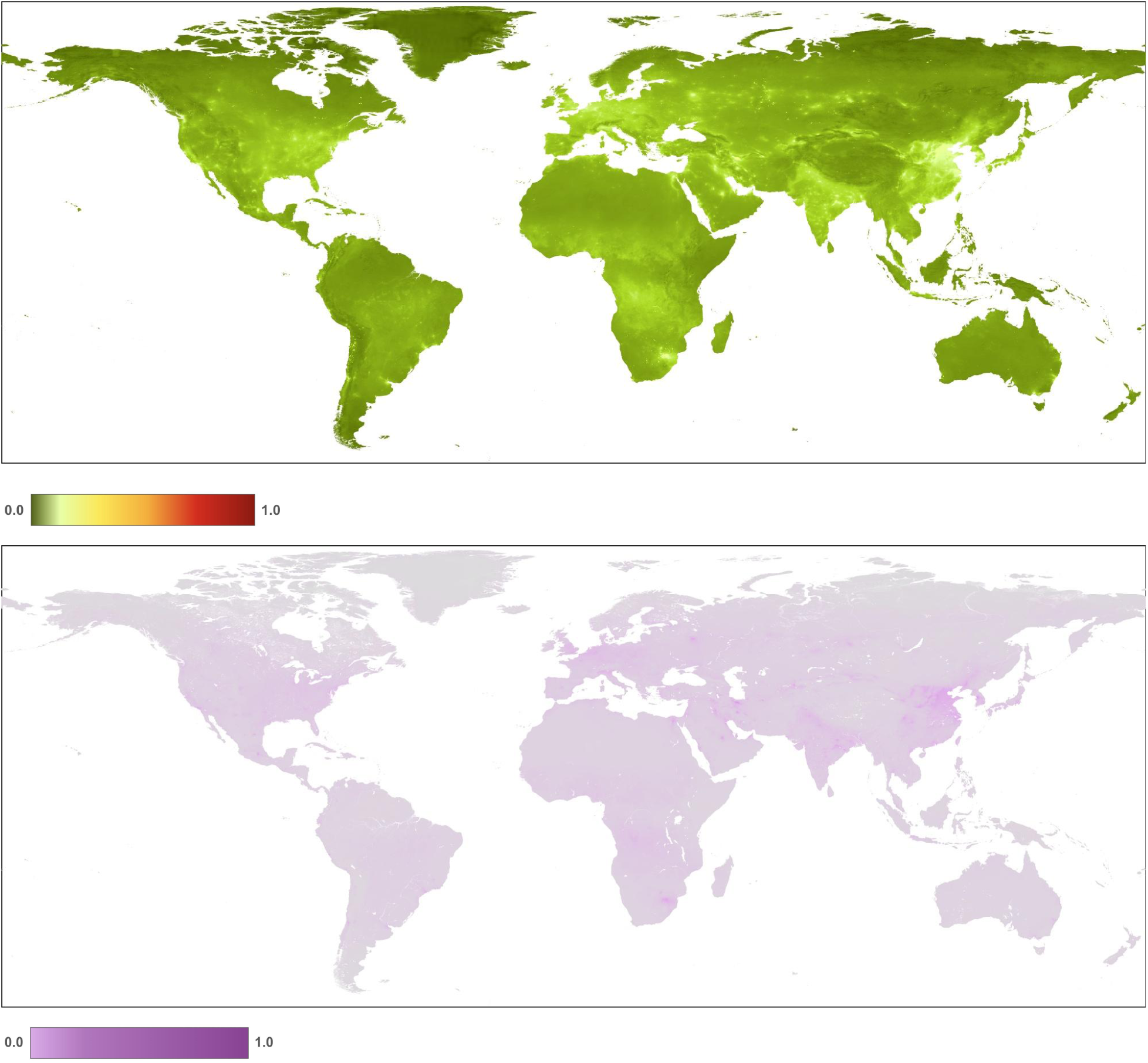
Maps of human modification values for pollution threat class (a) and (b) for air-borne pollutants (9.5). Note that for the garbage and solid-waste threat (9.4), patterns at the global extent are not discernable.

#### Combining threats

We then used the increasive mean function (Eq. 2) to calculate the overall degree of human modification *H* for 2022 combining all 16 threats: *H_ac_*, *H_ap_*, *H_ag_*, *H_br_*, *H_bc_*, *H_bt_*, *H_eo_*, *H_er_*, *H_ee_*, *H_tr_*, *H_tl_*, *H_f_*, *H_h_*, *H_n_*, *H_pg_*, and *H_pa_*. The built-up tourism & recreation (1.3), extractive mining (3.2), transportation - roads/railways (4.1), and pollution - solid waste/garbage (9.4; i.e., *H_bt_*, *H_em_*, *H_tr_*, *H_pg_*) threat classes were excluded from the 1990-2020 time-series datasets because no time-series data were available.

We found that the mean value of *H* in 2022 at 300 m resolution for global terrestrial lands is 0.1120 (±0.212; Figure 10, Table 3). Across the seven ecological realms, mean *H* values ranged from 0.0579 (Australasia) to 0.2930 (Indomalayan). For the 15 biomes, mean *H* values ranged from 0.0046 (Tundra) to 0.2848 (Temperate Broadleaf & Mixed Forests). Of the 822 ecoregions overlapping the HM datasets, mean *H* values ranged from 0.0001 to 0.9179. We note that estimates of average values of human modification are different from the area of natural or developed lands, as the latter requires additional assumptions about threshold values. For example, using the human modification classes^33^, 43% globally was at very low (*H*<0.01), 27% at low (0.01-0.1), 20% at moderate (0.1-0.4), 8% at high, and 2% at very high modification (*H*>0.7). The mean *H* value for the higher resolution HM (90 m) was 0.1078, slightly lower than the 300 m mean of 0.1120 (Table 3). This follows a typical pattern when calculating arithmetic means at a coarser resolution causes slight spatial smoothing of values and patterns.

**Figure 10.**
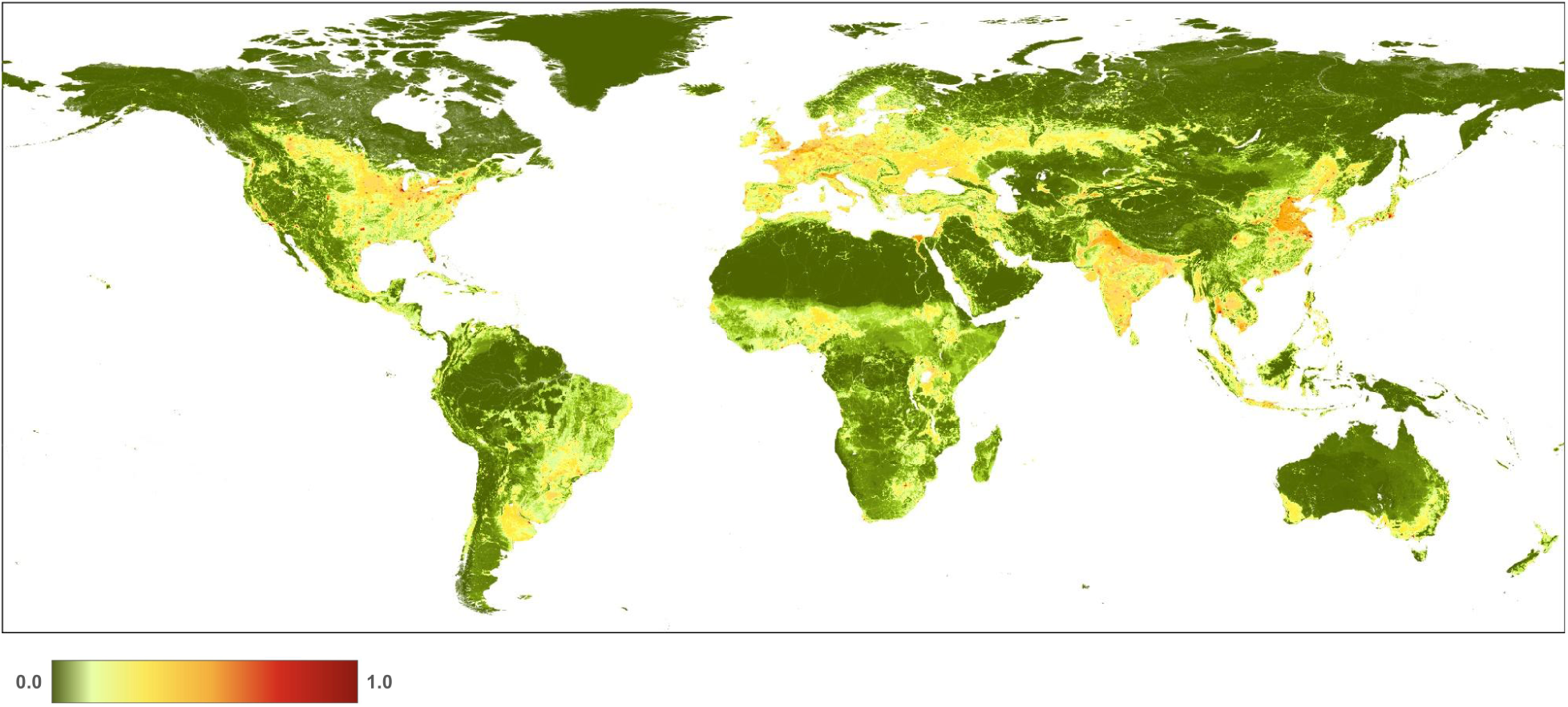
A map of the overall human modification (or cumulative threats) data layer for 2022.

**Table 3.**
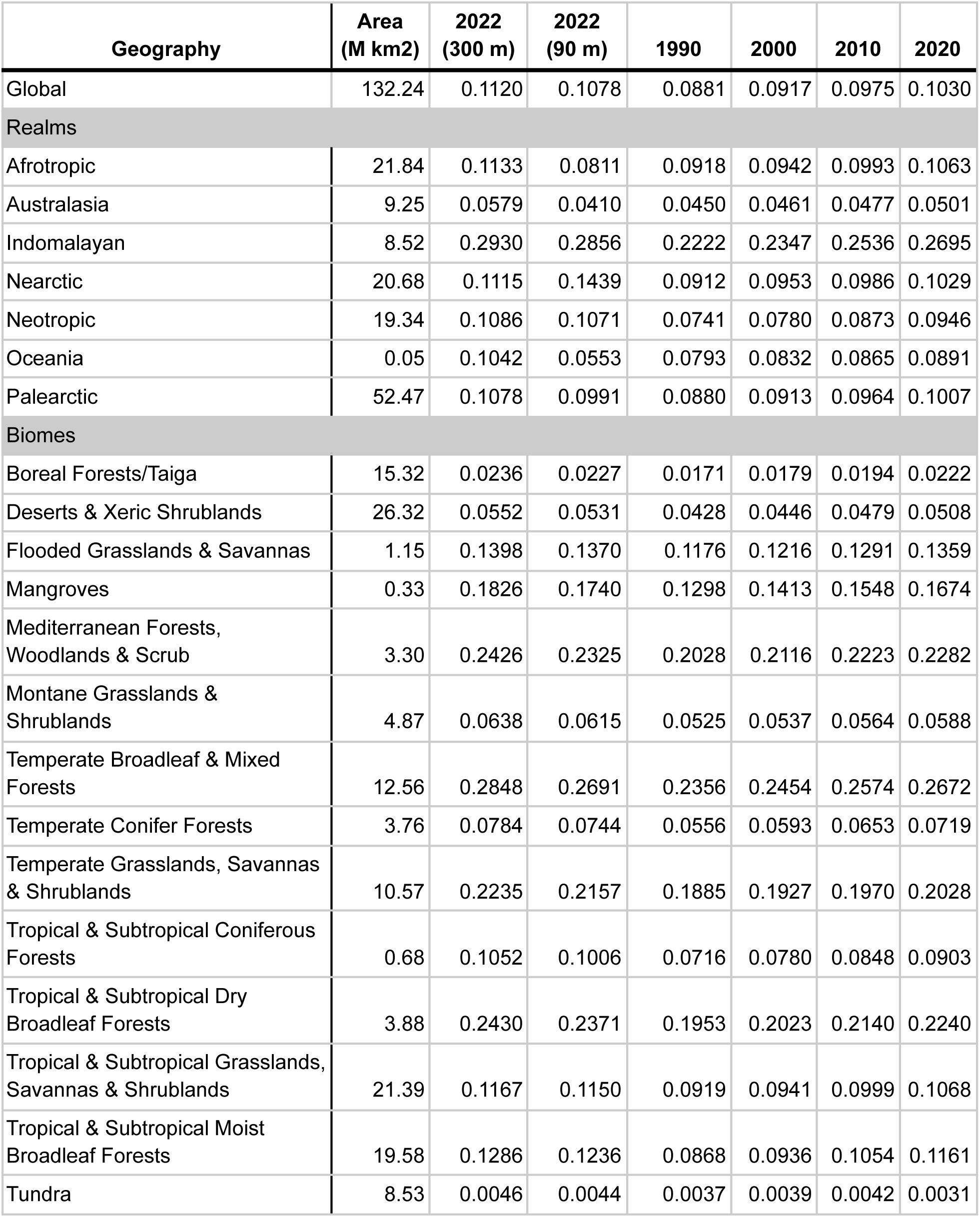
Mean human modification values for the cumulative threat and threat classes for 2022 and 1990-2020, globally, by terrestrial realm and biome.

| **Geography** | **Area**<br>**(M km<sup>2</sup>)** | **2022**<br>**(300 m)** | **2022**<br>**(90 m)** | **1990** | **2000** | **2010** | **2020** |
| --- | --- | --- | --- | --- | --- | --- | --- |
| Global | 132.24 | 0.1120 | 0.1078 | 0.0881 | 0.0917 | 0.0975 | 0.1030 |
| **Realms** |  |  |  |  |  |  |  |
| Afrotropic | 21.84 | 0.1133 | 0.0811 | 0.0918 | 0.0942 | 0.0993 | 0.1063 |
| Australasia | 9.25 | 0.0579 | 0.0410 | 0.0450 | 0.0461 | 0.0477 | 0.0501 |
| Indomalayan | 8.52 | 0.2930 | 0.2856 | 0.2222 | 0.2347 | 0.2536 | 0.2695 |
| Nearctic | 20.68 | 0.1115 | 0.1439 | 0.0912 | 0.0953 | 0.0986 | 0.1029 |
| Neotropic | 19.34 | 0.1086 | 0.1071 | 0.0741 | 0.0780 | 0.0873 | 0.0946 |
| Oceania | 0.05 | 0.1042 | 0.0553 | 0.0793 | 0.0832 | 0.0865 | 0.0891 |
| Palearctic | 52.47 | 0.1078 | 0.0991 | 0.0880 | 0.0913 | 0.0964 | 0.1007 |
| **Biomes** |  |  |  |  |  |  |  |
| Boreal Forests/Taiga | 15.32 | 0.0236 | 0.0227 | 0.0171 | 0.0179 | 0.0194 | 0.0222 |
| Deserts & Xeric Shrublands | 26.32 | 0.0552 | 0.0531 | 0.0428 | 0.0446 | 0.0479 | 0.0508 |
| Flooded Grasslands & Savannas | 1.15 | 0.1398 | 0.1370 | 0.1176 | 0.1216 | 0.1291 | 0.1359 |
| Mangroves | 0.33 | 0.1826 | 0.1740 | 0.1298 | 0.1413 | 0.1548 | 0.1674 |
| Mediterranean Forests,<br>Woodlands & Scrub | 3.30 | 0.2426 | 0.2325 | 0.2028 | 0.2116 | 0.2223 | 0.2282 |
| Montane Grasslands &<br>Shrublands | 4.87 | 0.0638 | 0.0615 | 0.0525 | 0.0537 | 0.0564 | 0.0588 |
| Temperate Broadleaf & Mixed<br>Forests | 12.56 | 0.2848 | 0.2691 | 0.2356 | 0.2454 | 0.2574 | 0.2672 |
| Temperate Conifer Forests | 3.76 | 0.0784 | 0.0744 | 0.0556 | 0.0593 | 0.0653 | 0.0719 |
| Temperate Grasslands, Savannas<br>& Shrublands | 10.57 | 0.2235 | 0.2157 | 0.1885 | 0.1927 | 0.1970 | 0.2028 |
| Tropical & Subtropical Coniferous<br>Forests | 0.68 | 0.1052 | 0.1006 | 0.0716 | 0.0780 | 0.0848 | 0.0903 |
| Tropical & Subtropical Dry<br>Broadleaf Forests | 3.88 | 0.2430 | 0.2371 | 0.1953 | 0.2023 | 0.2140 | 0.2240 |
| Tropical & Subtropical Grasslands,<br>Savannas & Shrublands | 21.39 | 0.1167 | 0.1150 | 0.0919 | 0.0941 | 0.0999 | 0.1068 |
| Tropical & Subtropical Moist<br>Broadleaf Forests | 19.58 | 0.1286 | 0.1236 | 0.0868 | 0.0936 | 0.1054 | 0.1161 |
| Tundra | 8.53 | 0.0046 | 0.0044 | 0.0037 | 0.0039 | 0.0042 | 0.0031 |

<sup>1</sup>Biome name: "Mediterranean forests, woodlands & scrub; <sup>2</sup>Biome name: "Temperate Grasslands, Savannahs, & Shrub"

From 1990-2020 (see Table 3; Figure 11), globally the mean *H* increased from 0.0881 in 1990 to 0.1030 in 2020, and the annual rate of increase for the overall *H* was 0.57%. Table 4 shows that the agricultural threat class had the highest global mean value of 0.0607, followed by transportation & service corridors (0.0241), while energy production & mining had the lowest mean value (0.00002). The annual rate of increase by threat class was 0.24% for agriculture, 4.20% for built-up, 0.19% for human accessibility, 0.53% for pollution, and 1.17% for transportation & service corridors. By threat class, energy production had the lowest mean *H* value, largely because of the small areal extent of oil & gas wells, and renewable energy facilities, though they are likely under-represented in the datasets available.

**Figure 11.**
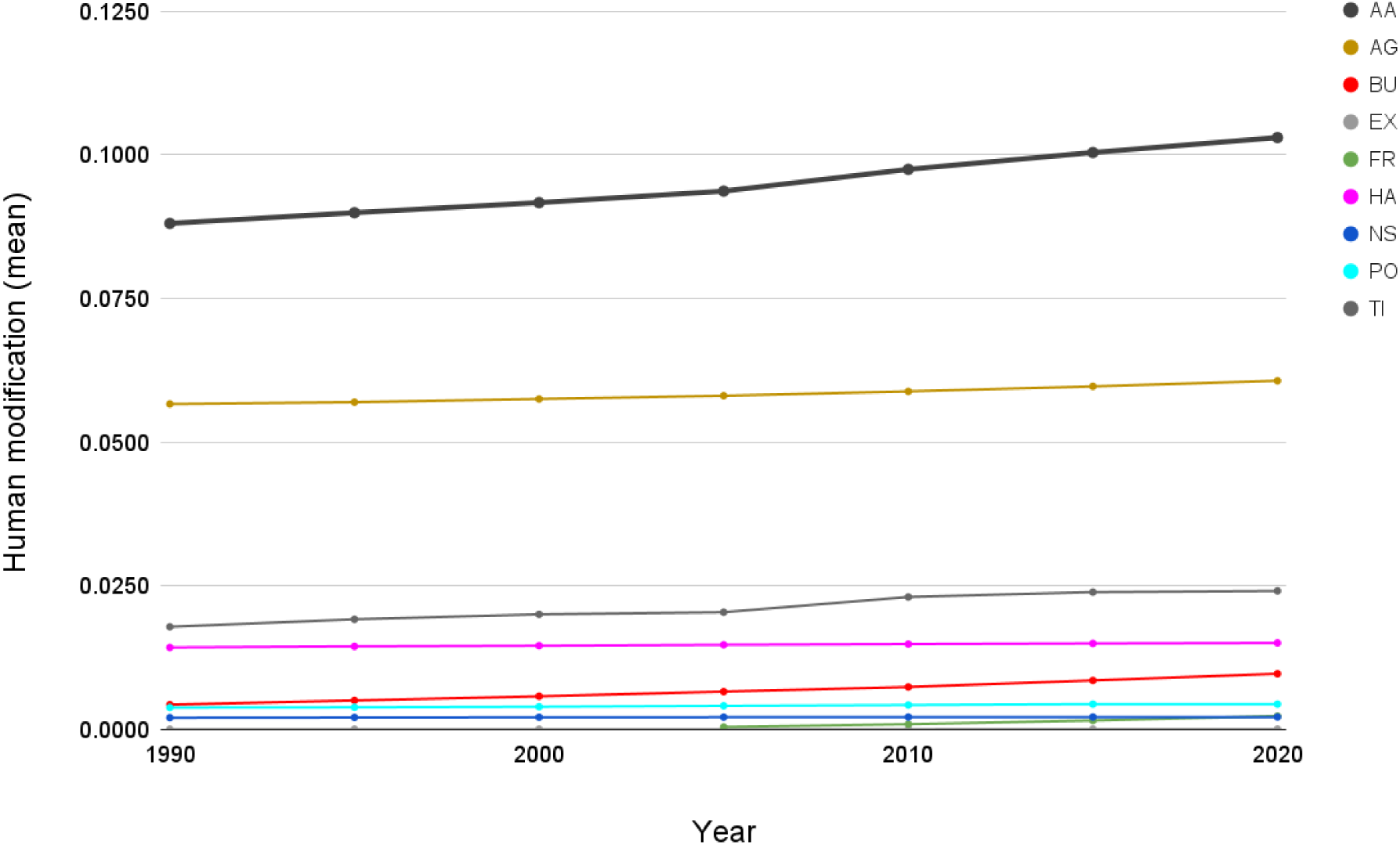
Trends in the mean terrestrial global human modification from 1990 to 2020. AA is the overall (cumulative threat) human modification; AG: agriculture (croplands, grazing, plantations); BU: residential & commercial development; EX: energy production & mining; FR: Biological resource use (logging & wood harvesting); HA: human accessibility; NS: natural systems modification (reservoirs), PO: pollution, and TI: transportation and service corridors.

**Table 4.** The mean human modification values for the 2022 cumulative threat and threat classes and 1990-2020, globally. “-” indicates data were not available for those time periods.

|  | 2022 | 2022<br>(90 m) | 1990 | 1995 | 2000 | 2005 | 2010 | 2015 | 2020 |
| --- | --- | --- | --- | --- | --- | --- | --- | --- | --- |
| Cumulative threat (AA) | 0.1120 | 0.1078 | 0.0881 | 0.0899 | 0.0917 | 0.0937 | 0.0975 | 0.1004 | 0.1030 |
| Agriculture (AG) | 0.0671 | 0.0665 | 0.0566 | 0.0569 | 0.0575 | 0.0581 | 0.0588 | 0.0597 | 0.0607 |
| Residential & commercial development (BU) | 0.0157 | 0.0130 | 0.0043 | 0.0050 | 0.0057 | 0.0065 | 0.0074 | 0.0085 | 0.0097 |
| Energy production & mining (EX) | 0.0015 | 0.0015 | 0.00000 | 0.00000 | 0.00000 | 0.00000 | 0.00000 | 0.00001 | 0.00002 |
| Biological resource use (FR) | 0.0023 | 0.0027 | - | - | - | 0.0004 | 0.0009 | 0.0015 | 0.0023 |
| Human accessibility (HA) | 0.0151 | 0.0152 | 0.0142 | 0.0144 | 0.0145 | 0.0147 | 0.0148 | 0.0149 | 0.0150 |
| Natural systems modification (NS) | 0.0023 | 0.0023 | 0.0020 | 0.0021 | 0.0021 | 0.0021 | 0.0021 | 0.0021 | 0.0021 |
| Pollution (PO) | 0.0044 | 0.0044 | 0.0038 | 0.0038 | 0.0039 | 0.0041 | 0.0042 | 0.0044 | 0.0044 |
| Transportation & service corridors (TI) | 0.0231 | 0.0196 | 0.0178 | 0.0191 | 0.0200 | 0.0204 | 0.0230 | 0.0239 | 0.0241 |

#### Dominant threat

We calculated the dominant threat class as the highest threat value per pixel (where *H* > 0.0). The threat class that caused the largest proportion of land conversion was agricultural (46.9%), followed by human intrusion (5.1%), transportation and electrical infrastructure (4.9%), and built-up (1.6%; Table 11, Supplemental Figure 12; Supplemental Table 4). Identifying the dominant threat is valuable for understanding potential conservation actions, such as protection, management, and/or restoration. Moreover, the detailed representation of different dominant threats provides critical and refined information to understand simplified classes of modification (e.g., the three conditions^95^).

Dominant human threats also vary with geographic distributions, though the majority of the land surface is subject to multiple threats – on average nearly 3 threats are present at each pixel– thus reinforcing the need to address the cumulative effects of industrial development on ecosystems.

The human modification datasets and supporting input data can be viewed interactively^96^, without GIS software, at https://hm-30x30.projects.earthengine.app/view/hm-v3.

#### Technical validation

We assessed the technical validity of our datasets by validating them against reference data (so-called “truth”) observations obtained using a robust sampling and response design, following well-established guidance^97^; and conducted sensitivity analyses of estimated model parameters and classification errors in land cover datasets. We augmented these technical validation analyses with visual and descriptive statistical comparisons with related datasets; obtained expert reviews through organized, structured by global and regional ecologists and data scientists; and compared results to authoritative estimates of land cover and ecoregional summaries.

We assessed the accuracy of our maps using the Global Land Use Emergent Dataset (GLUED) validation dataset following validation procedures^33,98^ to test the contemporary conditions of human modification (2022 map) that included all threats. We used an independent validation dataset from^33^ that quantified the degree of human modification from visual interpretation of high-resolution aerial or satellite imagery across the world (for ∼2015-2017), based on a spatially-balanced, probability-based random sampling design^98^. The randomization was stratified on a five-class rural-to-urban gradient using stable nighttime lights 2013 imagery^58^. For 1,000 plots of roughly 1 km^2^, we selected 10 simple random locations to capture rare features and the spatial heterogeneity in land use and land cover (for a total of 10,000 subplots), which were separated by a minimum distance of 100 m. The spatially balanced nature of the design maximizes statistical information extracted from each plot, because it increases the number of samples in relatively rare areas that are likely of interest (in contrast to simple random sampling) – especially for urbanized and growing cities.

Using GLUED as the validation dataset, the root-mean squared error (RMSE) for the HM v3 dataset was 0.180 and 0.178 (300 and 90 m), a slight improvement compared to previous versions of HM (v2 = 0.213 and v1=0.180).

Although accuracy assessments provide valuable insight into the quality of a dataset, other factors regarding the quality of spatial datasets remain important, especially mapping realistic and ecologically meaningful patterns that are difficult to assess statistically but are readily observable. Moreover, accuracy assessments themselves often have significant errors depending on the quality of the “truth” data, as well as the sampling balance among class categories^100^.

## Sensitivity analyses

Like other indices of human influence, the HM is sensitive to the error in input datasets and the chosen intensity values for each threat^101^. We evaluated the sensitivity of our findings to two primary sources of uncertainty: intensity estimates and land cover classification errors. To understand the sensitivity of our results to our estimates of intensity by threat^33^, we recalculated *H* by replacing “best estimate” values with values generated by drawing random values (*n*=100) between the minimum and maximum intensity values (Table 2), and then calculated the per-pixel mean and standard deviation for the randomizations at 1 km^2^ resolution for computational efficiency and provided corresponding maps. The mean *H* due to potential intensity errors across the 30 iterations was 0.1075, with uncertainty measured as SD=0.0038. Thus, the global overall *H* mean of 0.1078 obtained using our best-estimate intensity values was in line with our uncertainty results.

Next, we quantified the sensitivity of the results of the overall HM and component threats to potential errors in input datasets. Although most if not all datasets likely have some level of error^96^, we examined the likely primary contributors to error – land use and cover datasets. We also pragmatically restricted our analyses to datasets with readily-available confusion matrices. Specifically, we quantified sensitivity for the agricultural - cropland (2.1) threat class, for roughly 2020 conditions at 1 km resolution, from confusion matrices of the ESA CCI land cover (300 m), GCI cropping intensity (250 m), and global cropland extent (30 m) datasets. We focused only on this class as it is the largest contributor to overall HM at the global scale^16^, and the challenge of quantifying sensitivity to non-landcover-based datasets due to lack of confusion matrices.

For each dataset, we performed a Monte Carlo simulation, with the probability of sampling a value for a particular pixel determined by the accuracy of the associated dataset. We sampled 100 realizations for every input dataset to create 100 realizations of the agricultural components of the overall HM. For example, if the predicted class from the CCI land cover dataset is cropland, we sampled a new land cover class for that pixel in each of the 100 realizations. The probability of each possible land cover class was determined by the frequency of true land cover classes when a prediction of cropland is made, as reported in the confusion matrix. Because these confusion matrices provide overall, but not spatially explicit, estimates of uncertainty, we summarized uncertainty by 14 terrestrial biomes^102^ (Table 5). Mean biome-level average *H* associated with agriculture - croplands calculated from the mean of 100 realizations of input datasets and the actual input datasets were very similar. For most biomes, *H* slightly increased when sampling from realizations of input datasets. This follows from non-cropland pixels having low, but non-zero probabilities, of being sampled as cropland in our Monte Carlo analysis. The large areas of most biomes mapped as non-cropland then contribute a low, but non-zero, *H* value when averaging over many simulations. The standard deviation of mean biome-level cropland HM was very low for all biomes. Given the random, spatially unstructured sampling of error used, it is unsurprising that means converge when sampling over large areas composed of many pixels. These results show that if we assume spatially random error, misclassification in input datasets does not substantially alter regional cropland-related HM, which we assume also applies to other threat classes. Spatially unstructured error is a strong assumption likely to be violated in reality, leading to regional under- and overestimations of *H*, which we were unable to account for based on the available information.

**Table 5.** Uncertainty of the human modification values (*H*) associated with the agriculture - cropland threat class due to misclassifications of land cover datasets, summarized for 14 biomes. Mean and SD of agriculture-cropland *H* represents the mean and standard deviation of mean biome-level HM *H* 100 simulations in which land cover and cropland datasets were sampled based on their reported error matrices.

| Biome name | Mean | SD | Mean (original HM) | Area (million km<sup>2</sup>) |
| --- | --- | --- | --- | --- |
| Boreal Forests/Taiga | 0.020 | 0.000002 | 0.000 | 15.32 |
| Deserts & Xeric Shrublands | 0.040 | 0.000004 | 0.030 | 26.32 |
| Flooded Grasslands & Savannas | 0.110 | 0.000006 | 0.100 | 1.15 |
| Mangroves | 0.100 | 0.000024 | 0.090 | 0.33 |
| Mediterranean Forests, Woodlands & Scrub | 0.150 | 0.000010 | 0.150 | 3.30 |
| Montane Grasslands & Shrublands | 0.050 | 0.000003 | 0.040 | 4.87 |
| Temperate Broadleaf & Mixed Forests | 0.150 | 0.000004 | 0.150 | 12.56 |
| Temperate Conifer Forests | 0.040 | 0.000009 | 0.020 | 3.76 |
| Temperate Grasslands, Savannas & Shrublands | 0.170 | 0.000003 | 0.160 | 10.57 |
| Tropical & Subtropical Coniferous Forests | 0.050 | 0.000018 | 0.040 | 0.68 |
| Tropical & Subtropical Dry Broadleaf Forests | 0.170 | 0.000006 | 0.160 | 3.88 |
| Tropical & Subtropical Grasslands, Savannas & Shrublands | 0.100 | 0.000004 | 0.090 | 21.39 |
| Tropical & Subtropical Moist Broadleaf Forests | 0.080 | 0.000003 | 0.070 | 19.58 |
| Tundra | 0.010 | 0.000003 | 0.000 | 8.53 |

We recognize that we provide an incomplete characterization of the uncertainty in the input datasets, given our assumption that prediction errors are spatially independent and consistent within a class. However, a more rigorous uncertainty analysis would require the input dataset to provide predictions in the form of calibrated class probabilities that account for spatial structure in errors, which is rarely done^103^. Furthermore, the biome-level summaries we provide are based on simple pixel averaging. It is well known that pixel counting can lead to biased estimates of averages and areas when summarizing remote-sensing based datasets that are estimated with error^99^. The biome-level summaries and standard deviations we report should therefore not be interpreted as reflecting the true mean and variation, but rather simply the sensitivity to uncertainty in input datasets.

## Visual and statistical comparison

Throughout our modeling process, we inspected the source, intermediate, output, and validation datasets, using a dynamic map browser that allowed for coauthors and an external advisory committee to visually interpret and query map values and provide feedback. Our advisory committee consisted of two-dozen external scientists and conservation practitioners with both global and regional ecological expertise, particularly in the regions of: Andes, Angola, Argentina, Australia, Brazil, Canada, Colombia, Chile, China, Indonesia, Kenya, Mexico, Patagonia, South Africa, and United States. We obtained feedback through online workshops and via dynamic map viewers that allowed experts to examine and provide their feedback on areas they were knowledgeable about. This feedback contributed to refining aspects such as formulation of threat equations, intensity values, selection of input datasets, in particular related to forest and timber logging, pasture, and cropland threats.

We also compared the results of this work (HM v3) to other related data sources, specifically HM v1^16,33^, HM v2^10^, and the HFP 2009^7^, HFP 2013^104^. The mean value of HM v3 (0.1120) is slightly higher than HM v2 (0.1003) but lower than HM v1 (0.179; Table 6). The lower values of v2 & 3 are partly due to higher resolution, but also because updated livestock density data and model refinements resulted in a reduced spatial extent compared to v1. The reduced overall values (compared to v1) found here occur particularly in the Afrotropic, but minimally in Australasia. The ranks of the biomes with the highest modification values identified for the different HM and HF versions are fairly consistent – notably the top three biomes with the highest modification values in all datasets were: temperate broadleaf & mixed forests, tropical & subtropical dry broadleaf forests, and Mediterranean forests, woodlands & scrub.

**Table 6.** The mean human modification (HM) values by terrestrial biome for 2022, with comparisons to previous versions of HM and the human footprint (HF). Datasets are at 300 m resolution unless otherwise noted. (Note: the “rock & ice” biome in Antarctica and interior Greenland are not included here.) Datasets are from: HM v3=this work; HM v2^10^; HM v1^16^; HF 2013^104^, and HF 2009^7^.

|  | Area<br>(M km<sup>2</sup>) | HM v3<br>(2022) | HM v3<br>(2022, 90 m) | HM v2<br>(2017) | HM v1<br>(2016)* | HF<br>(2013)* | HF<br>(2009)* |
| --- | --- | --- | --- | --- | --- | --- | --- |
| Global | 132.24 | 0.112 | 0.1078 | 0.1003 | 0.179 | 0.137 | 0.113 |
| Realms |  |  |  |  |  |  |  |
| Afrotropic | 21.84 | 0.1133 | 0.1116 | 0.0811 | 0.2325 | 0.1617 | 0.1277 |
| Australasia | 9.25 | 0.0579 | 0.0558 | 0.0410 | 0.0749 | 0.0811 | 0.0779 |
| Indomalayan | 8.52 | 0.2930 | 0.2818 | 0.2856 | 0.4352 | 0.2869 | 0.2619 |
| Nearctic | 20.68 | 0.1115 | 0.1055 | 0.1439 | 0.1267 | 0.1271 | 0.0790 |
| Neotropic | 19.34 | 0.1086 | 0.1061 | 0.1071 | 0.1815 | 0.1206 | 0.1074 |
| Oceania | 0.05 | 0.1042 | 0.0986 | 0.0553 | 0.1858 | 0.1809 | 0.1946 |
| Palearctic | 52.47 | 0.1078 | 0.1030 | 0.0991 | 0.1778 | 0.1421 | 0.1183 |
| Biomes |  |  |  |  |  |  |  |
| Boreal Forests/Taiga | 15.32 | 0.0236 | 0.0227 | 0.0235 | 0.041 | 0.046 | 0.029 |
| Deserts & Xeric Shrublands | 26.32 | 0.0552 | 0.0531 | 0.0381 | 0.104 | 0.098 | 0.082 |
| Flooded Grasslands & Savannas | 1.15 | 0.1398 | 0.1370 | 0.1102 | 0.243 | 0.172 | 0.142 |
| Mangroves | 0.33 | 0.1826 | 0.1740 | 0.1700 | 0.301 | 0.228 | 0.199 |
| Mediterranean Forests,<br>Woodlands & Scrub | 3.30 | 0.2426 | 0.2325 | 0.2293 | 0.332 | 0.271 | 0.216 |
| Montane Grasslands &<br>Shrublands | 4.87 | 0.0638 | 0.0615 | 0.0422 | 0.153 | 0.118 | 0.108 |
| Temperate Broadleaf & Mixed<br>Forests | 12.56 | 0.2848 | 0.2691 | 0.3089 | 0.396 | 0.311 | 0.249 |
| Temperate Conifer Forests | 3.76 | 0.0784 | 0.0744 | 0.0881 | 0.139 | 0.140 | 0.099 |
| Temperate Grasslands, Savannas<br>& Shrublands | 10.57 | 0.2235 | 0.2157 | 0.1955 | 0.298 | 0.227 | 0.167 |
| Tropical & Subtropical Coniferous<br>Forests | 0.68 | 0.1052 | 0.1006 | 0.0992 | 0.267 | 0.173 | 0.157 |
| Tropical & Subtropical Dry<br>Broadleaf Forests | 3.88 | 0.2430 | 0.2371 | 0.2199 | 0.377 | 0.249 | 0.226 |
| Tropical & Subtropical Grasslands,<br>Savannas & Shrublands | 21.39 | 0.1167 | 0.1150 | 0.0857 | 0.210 | 0.148 | 0.121 |
| Tropical & Subtropical Moist Broadleaf Forests | 19.58 | 0.1286 | 0.1236 | 0.1412 | 0.231 | 0.152 | 0.139 |
| Tundra | 8.53 | 0.0046 | 0.0044 | 0.0028 | 0.009 | 0.010 | 0.007 |

\*1-km resolution

**Table 7.** Comparison of mean threat values for areas within 100 km radius, centered on Asuncion, Canada; Calgary, Canada; Qing Yuan, China and Timbuktu, Mali. Note that these global datasets can be visualized and explored further at: https://hm-30x30.projects.earthengine.app/view/hm-v3.

|  | Year | Resolution (m) | Asuncion | Calgary | Qing Yuan | Timbuktu | Global |
| --- | --- | --- | --- | --- | --- | --- | --- |
| HM v3 | 2022 | 300 | 0.1391 | 0.2859 | 0.3549 | 0.0611 | 0.1120 |
| HM v3 90 m | 2022 | 90 | 0.1345 | 0.2769 | 0.3203 | 0.0608 | 0.1078 |
| HM v2 | 2017 | 300 | 0.1119 | 0.4418 | 0.4307 | 0.0082 | 0.1003 |
| HM v1 | 2015 | 1000 | 0.2906 | 0.4415 | 0.5044 | 0.1407 | 0.1787 |
| HFP 2009 | 2009 | 1000 | 0.1982 | 0.1883 | 0.3306 | 0.0854 | 0.1125 |
| HFP 2013 | 2013 | 300? | 0.2313 | 0.3108 | 0.4259 | 0.0937 | 0.1374 |
| HFP Mu | 2020 | 1000 | 0.2483 | 0.3466 | 0.4476 | 0.1167 | 0.1534 |
| CCI 300 | 2020 | 300 | 0.0606 | 0.6318 | 0.2171 | 0.0161 | 0.1296 |
| Not natural | 2020 | 30-300 | 0.0408 | 0.5224 | 0.8281 | 0.0319 | 0.2296 |

\*Max-normalized to scale 0 to 1.
\*\*Assuming urban/built-up and cropland cover types equal 1.0, other classes = 0.0.

In addition to comparing mean *H* values across realms and biomes (Tables 3 & 4) and to values from earlier versions (Table 6), we also provide maps and summaries for four example locations selected across four continents and placed within different urban to rural landscape settings (Supplementary Figures 1-11; Table 4), to highlight varying geographical contexts, but also to illustrate the expected greater variability of values obtained for smaller spatial areas. We also included a few additional relevant datasets for comparison among the example regions: HFP 2020^105^, and HFP 2021^106^ at 100 m, the ESA CCI developed classes^40^, and the natural/not-natural dataset^34^.

Briefly, values from HM v3 were slightly higher globally and the four regional examples (Asuncion, Calgary, Qing Yuan, and Timbuktu) than v2, and lower than v1. It is not surprising that each of the regional examples (∼31,400 km^2^) exhibited more variability amongst the different datasets than the global mean, but the large range in values (e.g., for Asuncion from 0.04-0.29) suggest that results differ substantially among these datasets. Relative to human footprint (HF) datasets (see Supplemental Table 4), HM v3 values are lower, despite HM representing 2-13 years more recent conditions and maps more threats. This is likely because HM explicitly does not represent assumed spatial processes such as edge effects along roads and railroads, while HF applies a distance-decay buffer for all roads, up to 14 km away. Also, the datasets generated through land cover classes (i.e. CCI 300 and natural/not-natural) vary more widely than the other datasets, and are sensitive to landscape context (e.g., they are the minimal values for Asunsino and Timbuktu but among the highest for Calgary and Qing Yuan). We provide a comparison of the data and methods across the different human pressure maps (Supplemental Table 5).

### Usage notes

#### Use in the Global Biodiversity Framework

We anticipate these HM datasets will provide foundational information to monitor and report on the CBD’s Global Biodiversity Framework (GBF). The majority of the roughly two-dozen indicators proposed to evaluate ecological integrity by the GBF rely on human pressure data in their calculations. For example, the headline indicator for Goal A is the “Extent of natural ecosystems”; and HM provides a fit-for-purpose, operational estimation of “natural” defined as the inverse of human modified^6,16^. Because HM represents the full gradient of unmodified or “wild” to developed conditions in a spatially-explicit, robust and comprehensive way, it extends beyond “natural” (or wild) categorization and covers the gradient of semi-natural ecosystems that retain substantial biodiversity and are thus important for conservation^98^. Moreover, biodiversity-inclusive spatial planning, as required by Target 1 of GBF, encompasses the range of actions from protection to restoration to address land change and abate threats^107^. The extent of current modification and their rates of change can inform strategic conservation prioritization and planning^108^.

Overall, we found that 24% of terrestrial ecosystems (31 M km^2^) experienced increases in *H* values (*H_2020-1990_* > 0.01) from 1990 to 2020. Further, we found that 29% of countries^109^ (*n=*181 member states, *n*=246 total) and 31% of ecoregions^105^ (*n*=775, *n=*813 total) might also be particularly vulnerable to biodiversity loss given that they have above-average rates of modification and less than 30% of lands protected (i.e. Target 3; country mean=0.0318, ecoregion mean=0.0235; Supplemental Figures 13 & 14).

We found that roughly ⅔ of biomes and ½ of ecoregions currently (∼2022) are moderately-modified (0.1 < *H* < 0.7). Moderately-modified lands experienced increases in areas developed (*H_2020-1990_*> 0.01) from 1990 to 2020: 36% of the area developed was from low-modified lands (*H*<0.1) in 1990, 45% in moderately-modified (0.1-0.4), 18% in moderately-high modified (0.4-0.7), and 1% in highly modified (H>0.7). Moderately-modified ecosystems may warrant elevated conservation attention if they fall within critical ecological or habitat thresholds^16,33^ to prevent precipitous biodiversity losses and impairments to ecosystem functioning^110^. The global mean H in 2022 for protected areas was substantially lower than in non-protected areas: 0.0437 (SD=0.0454) vs. 0.1252, respectively.

By using the IUCN Threat taxonomy, our maps can be directly linked to the IUCN Red List of Species assessments, for which we mapped 16 of the 24 threats identified in the level two taxonomy. The 16 threats mapped in the human modification datasets are the dominant threats for 89.1% of Red List species (*n*=49,291; i.e. species listed as categories as critically endangered, endangered, vulnerable, or near threatened; https://www.iucnredlist.org/search).

Although many indicators can benefit from directly incorporating HM maps as produced, we recommend minor adjustments if used as inputs for connectivity modeling, such as using least-cost distance, circuit theory^111^, or related approaches. Because narrow, linear features such as transportation are important causes of fragmentation, it may be reasonable to adjust the intensity values – for those specific transportation pixels – to reflect a higher value approaching 1.0. This is particularly important when upscaling HM datasets to a coarser resolution to produce a resistance surface for connectivity modeling, which commonly occurs to reduce the file size and improve computational performance of connectivity models. Approaches that might be appropriate include: setting pixels that intersect “large” roads (e.g,. highways) to higher intensity values (such as 0.81); calculating H’=H^0.5 for road cells which retains the pattern but increases them substantially; and incorporating additional information such as traffic volume, speed, or presences of wildlife-proof fencing^83,112^. Also, some threats may be removed that are less relevant to movement, such as air pollution, or ones with base data layers that cause artificial or abrupt differences in connectivity modeling outputs such as pasture maps with sharply contrasting values at administrative boundaries^18^.

#### Data caveats, limitations, and appropriate use

Mean *H* values are a primary way to summarize results across varying geographies such as realms, biomes, ecoregions, and countries. To make comparisons to other area-based statistics, such as the number of square kilometers of built-up lands derived from a land cover map, we recommend converting the raw *H* values to a measure of area, which requires specifying a value to threshold *H* values to classes, such as we have done using human modification class thresholds^10,16^ (e.g., low human modification *H*<0.1). Because estimates of area are likely sensitive to the threshold values, we recommend using sensitivity analyses to better understand and convey the potential change in area estimated. We caution against calling a simple classification of human pressure datasets “intactness”, because the spatial structure associated with the neighborhood surrounding each pixel needs to be explicitly addressed. Results from the 2022 data should not be compared to the previous 1990-2020 data series. Because the 2022 contains additional datasets that represent only “current” conditions, the *H* values for 2022 will be ≥ in 1990-2020 (e.g., 0.1120 vs. 0.1014 globally).

Although our HM datasets provide the most comprehensive cumulative threat maps to date, they do not include all human threats^15^. Important data gaps to fill in future work include data needed to map threats associated with invasive non-native species (8.1), fishing and harvesting aquatic resources (5.4), and agricultural and forestry effluents (9.3). Based on the IUCN Red List Species assessment, filling these data gaps would contribute better understanding to 5.5%, 2.4%, and 1.4% additional species (excluding climate change, fire & fire suppression, and geological hazard threats).

Data representing roads (e.g., OpenStreetmap) are limited due to a lack of temporal information to support rigorous temporal change calculations, and many omissions can occur for small, narrow roads in dense canopy forests^113^ (e.g., Congo forests) as well as dispersed and remote roads and trails due to resource exploration^114,115^ (e.g., seismic lines).

HM datasets generated here are intended to support global to regional scale analyses, and we caution applying these data at more local exents. By following HM framework and methodologies, users can downscale HM by applying more local spatial data and/or intensity values specific to their area of focus. We believe however the global HM products provide a starting point to initiate potential downscaling.

We also recognize that measures of human threats, including HM, do not directly measure ecosystem condition, though they indicate locations where human threats associated with industrial human pressures^11^ that cause land conversion or degradation (e.g., human settlement, cropland, plantations, transportation, mining, energy production, etc.). Datasets on human pressures are often used as a practical, surrogate measure of structural state and landscape characteristics^12–14^. In practice, high quality datasets to estimate the characteristics of condition, such as compositional state or functional state characteristics, like percent invasive species, are generally unavailable even at regional or local-project scales^116,117^. As such, we make the assumption that surrogate variables, such as where human use of ecosystems are located, provide some complementary information to ecological condition, particularly where dispersed and low-intensity human activities associated in particular with industrial economies, that likely have negative impacts on biodiversity occur (e.g., grazing, pollution, human accessibility), but do not necessarily result in measurable land cover conversion. For most ecosystems and many species, lands with lower anthropogenic pressure cause less impacts because less habitat has been converted or modified, and natural ecological processes are allowed to proceed. In addition, information about the land use context and primary threats at a given location is key to inform appropriate and useful conservation strategies, as well as being valuable to engage stakeholders within a project area. However, at a project or operational scale, more specific and detailed information on human threats and potentially on direct aspects of ecological condition should be used in place of the global HM datasets, particularly when “downscaling” human pressure maps.

#### Robust mapping of human threats

Because habitat destruction and degradation associated with industrial development and activities is the primary threat and dominant driver of biodiversity loss globally^118^, datasets that estimate human threats are critically valuable to understand the extent and/or change in the condition of natural systems. Our updated and refined datasets on human modification described here will provide comprehensive, consistent, detailed, robust, and contemporary datasets to map threats associated with industrial human activities to terrestrial biodiversity and ecosystems.

Previous versions of the HM framework and datasets have been widely used at local to national to global scales, including to map the loss of natural lands in the United States^6,28,119^, extinction risk of global terrestrial mammals^120^, the ecological integrity of the sagebrush biome^121^, ecosystem loss and fragmentation globally^16^, connectivity of protected areas in the US^122,123^, climate adaptation in North America^124^ and in China^125^, climate corridors in California^112^, protected area effectiveness^126^, functional connectivity^127^, global ecosystem integrity^128^, impacts on forest structural density^129^, hot spots of threatened terrestrial vertebrates^130^, and global conversion pressure^131^.

## Acknowledgements

This work was supported by The Nature Conservancy – Global Protect 30 × 30 Strategy and Global Science Program. We thank members of our expert and advisory teams for their valuable review and feedback on our modeling approach and outputs, including: A. Trainor, K. Hall, E. Poor, D. Terasaki Hart, J. Anderson, P. Ellis, N. Robinson, T. Chapman, C. Schloss, A. Jones, J. Evans, M. Clark, F.T. Brum, J. Alvarez-Romero, J. Fitzsimmons, J. Shyvers, F. Reyna Sáenz, H. Cardenas Rodriguez, M. Nunez Regueiro, Manuel Agra, M. Ibanez, A. Gulielmetti, J. Burbano-Giron, W. Longzhu, M. Rosenfield, M. Barroso, R. Drever, and T. Boucher. We also thank S. Roy and the Awesome GEE community catalog for making datasets available to the Google Earth Engine community. We greatly appreciate the contributors to OpenStreetMap. OpenStreetMap is copyrighted and data and contributors are available from https://www.openstreetmap.org. We thank the ESA CCI Land Cover project for providing land cover datasets and the Global Human Settlement Layers.

## Author Contributions

D.T., C.K., J.O., and J.K. conceived and designed the study; D.T., C.K., J.O. designed the methods; DT applied all methods and produced the datasets; D.T. and G.M. conducted validation and sensitivity analyses; D.T., C.K., J.O., G.M., M.V., and J.K. drafted or revised the manuscript; all authors reviewed analyses and the manuscript.

