## Supplementary Information for "Global extent and change in human modification of terrestrial ecosystems from 1990 to 2022"

Supplemental Figure 1. Global human modification for 2022, with example cities: A=Asuncion, Paraguay, B=Calgary, Canada; C=Qing Yuan, China; and D: Timbuktu, Mali. Note: datasets can be explored at: https://hm-30x30.projects.earthengine.app/view/hm-v3.)

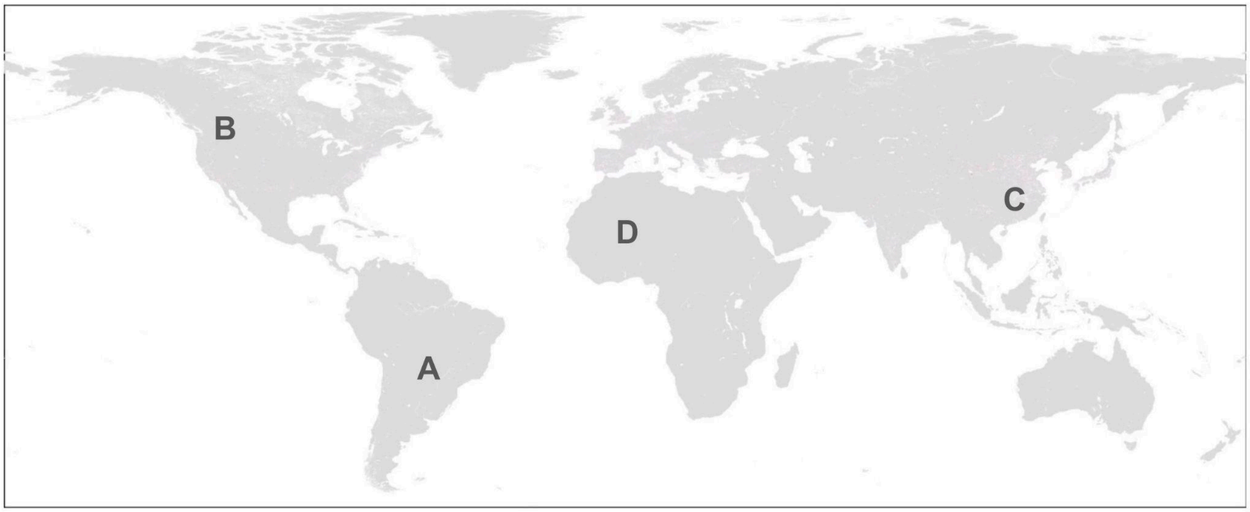

Supplemental Figure 2. Maps of human modification at 300 m for four example landscapes and their associated H for 2022 within 100 km radius of: (a, b) Asuncion, Paraguay; (c, d) Calgary, Canada; (e, f) Qing Yuan, China; and (g, h) Timbuktu, Mali. (Note: datasets can be visualized at: [https://hm-30x30.projects.earthengine.app/view/hm-v3](https://hm-30x30.projects.earthengine.app/view/hm-v3).

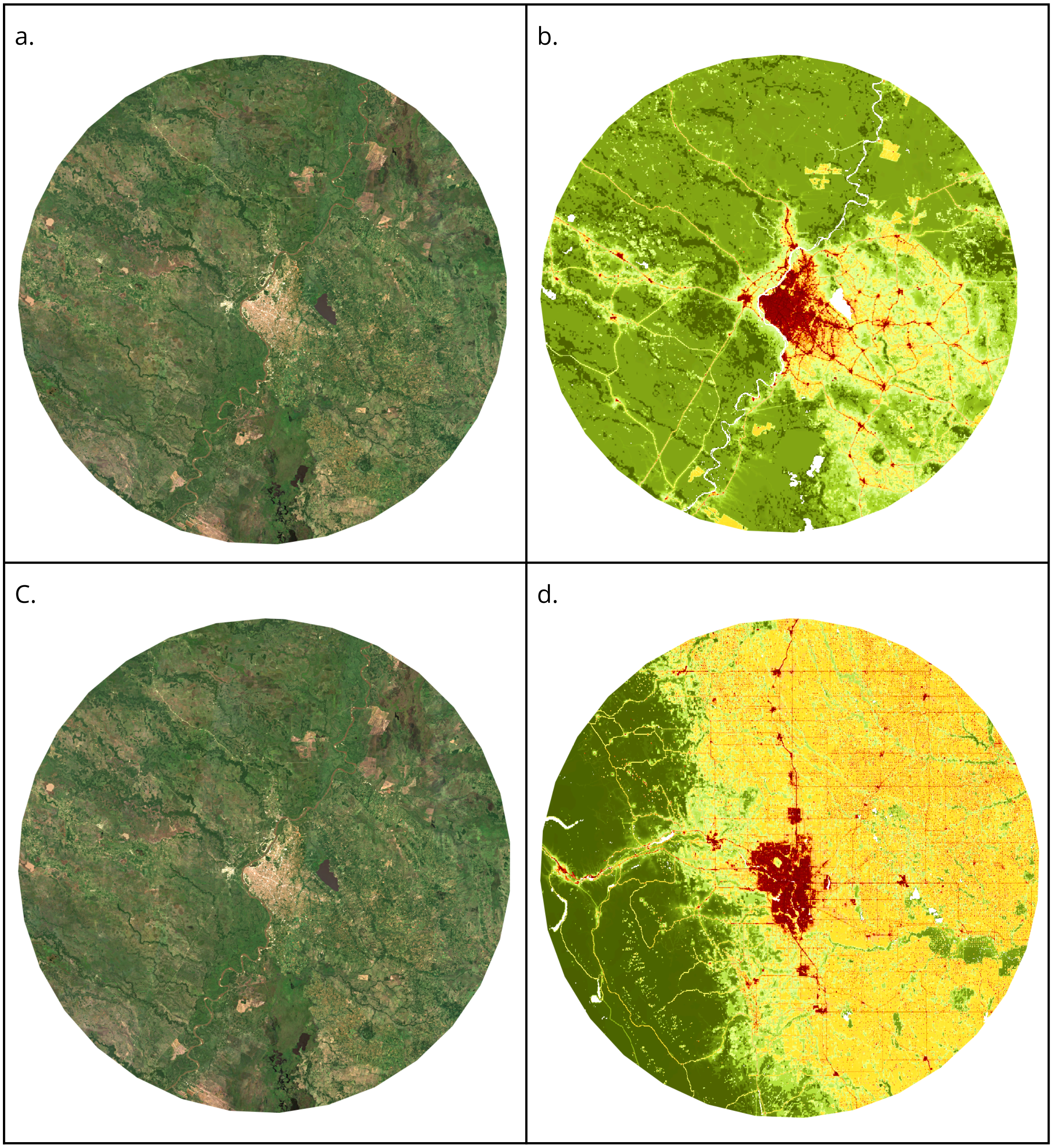

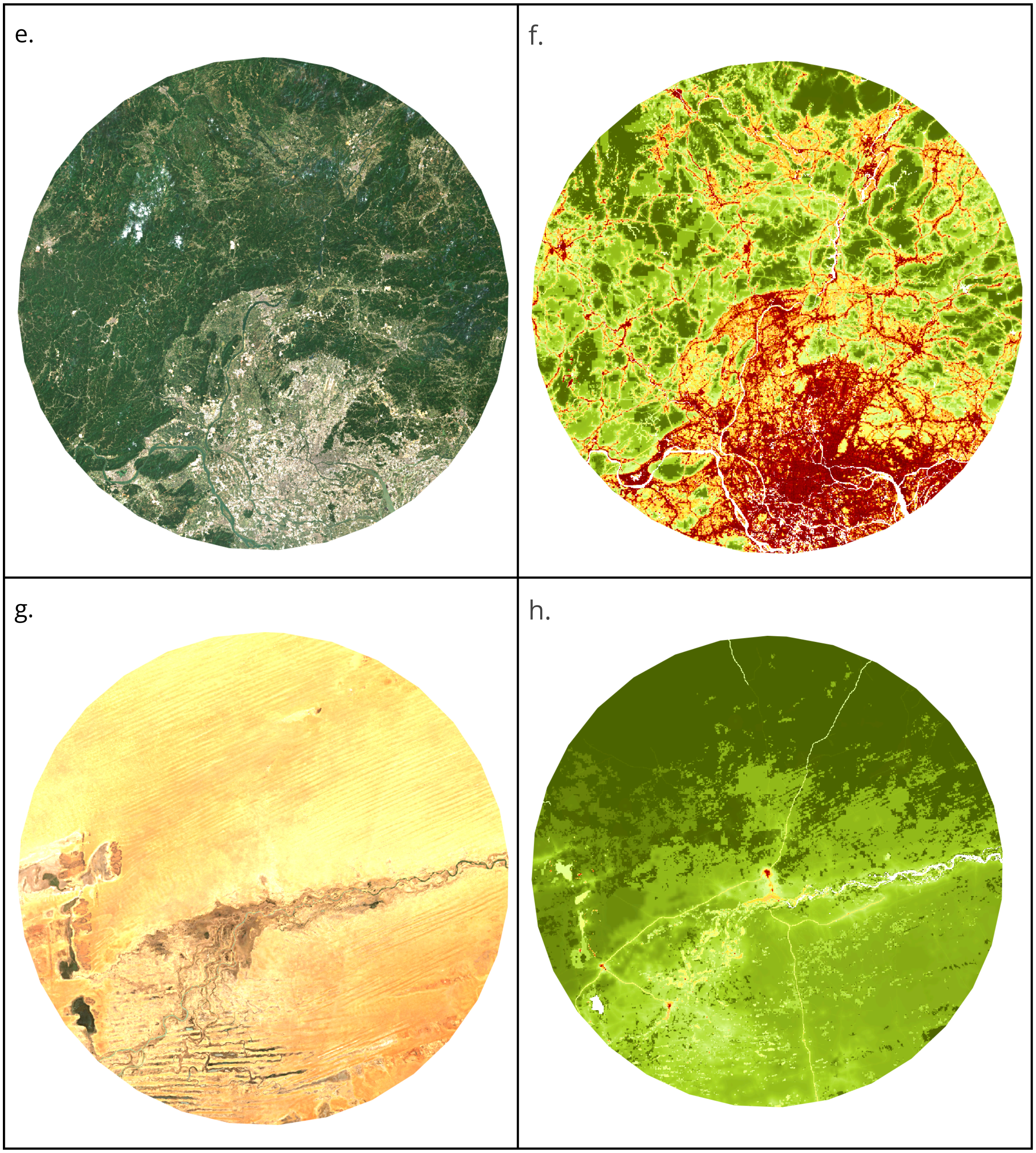
e.
f.
g.
h.

Supplemental Figure 3. Maps of human modification at four example landscapes and their associated H for 2022 within 100 km radius: (a) Asuncion, Paraguay at 300 m & (b) Asuncion, Paraguay at 90 m; (c & d) Calgary, Canada; (c) Qing yuan, China; and Timbuktu, Mali. (Note: datasets can be visualized at: [https://hm-30x30.projects.earthengine.app/view/hm-v3](https://hm-30x30.projects.earthengine.app/view/hm-v3).

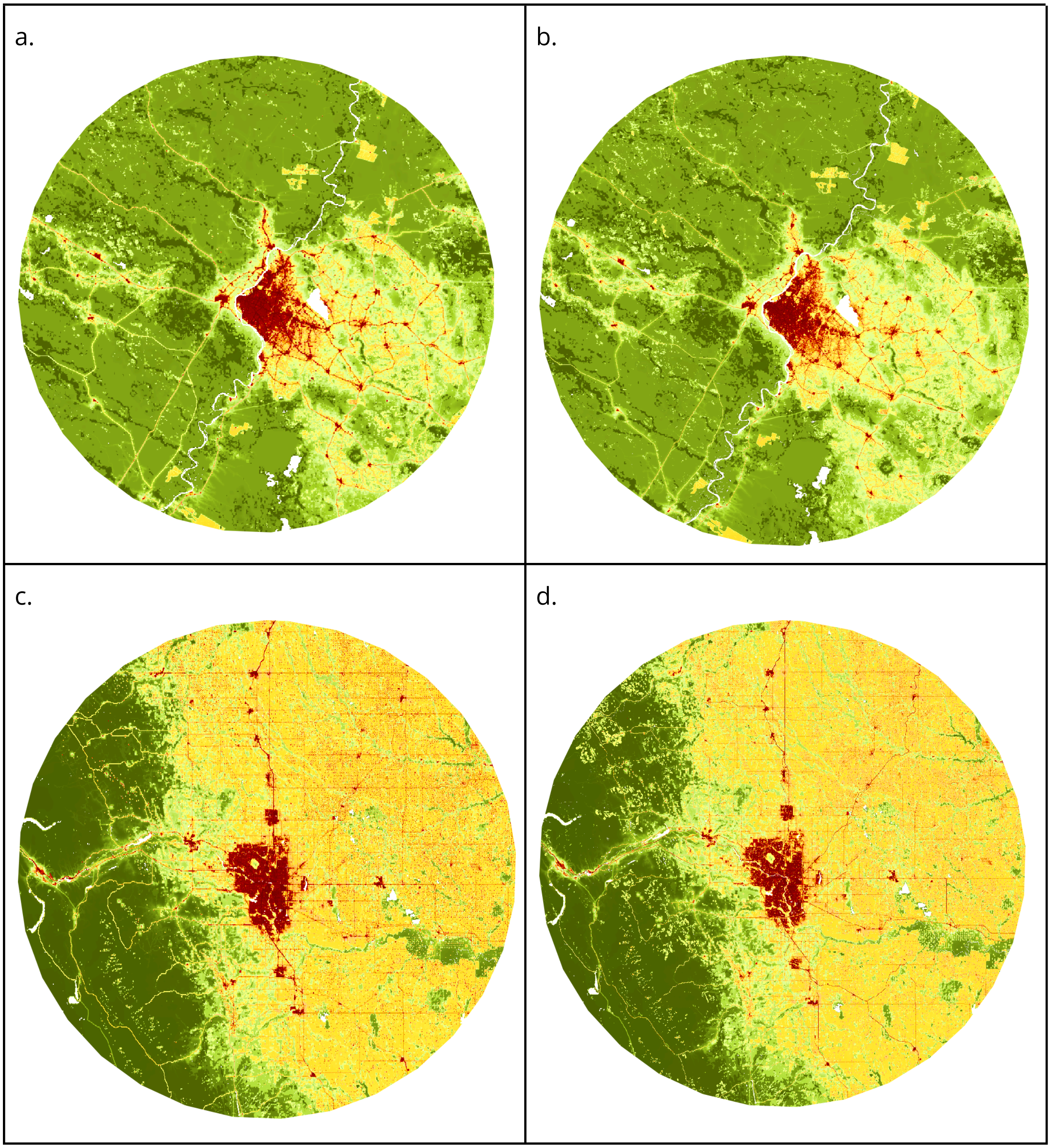

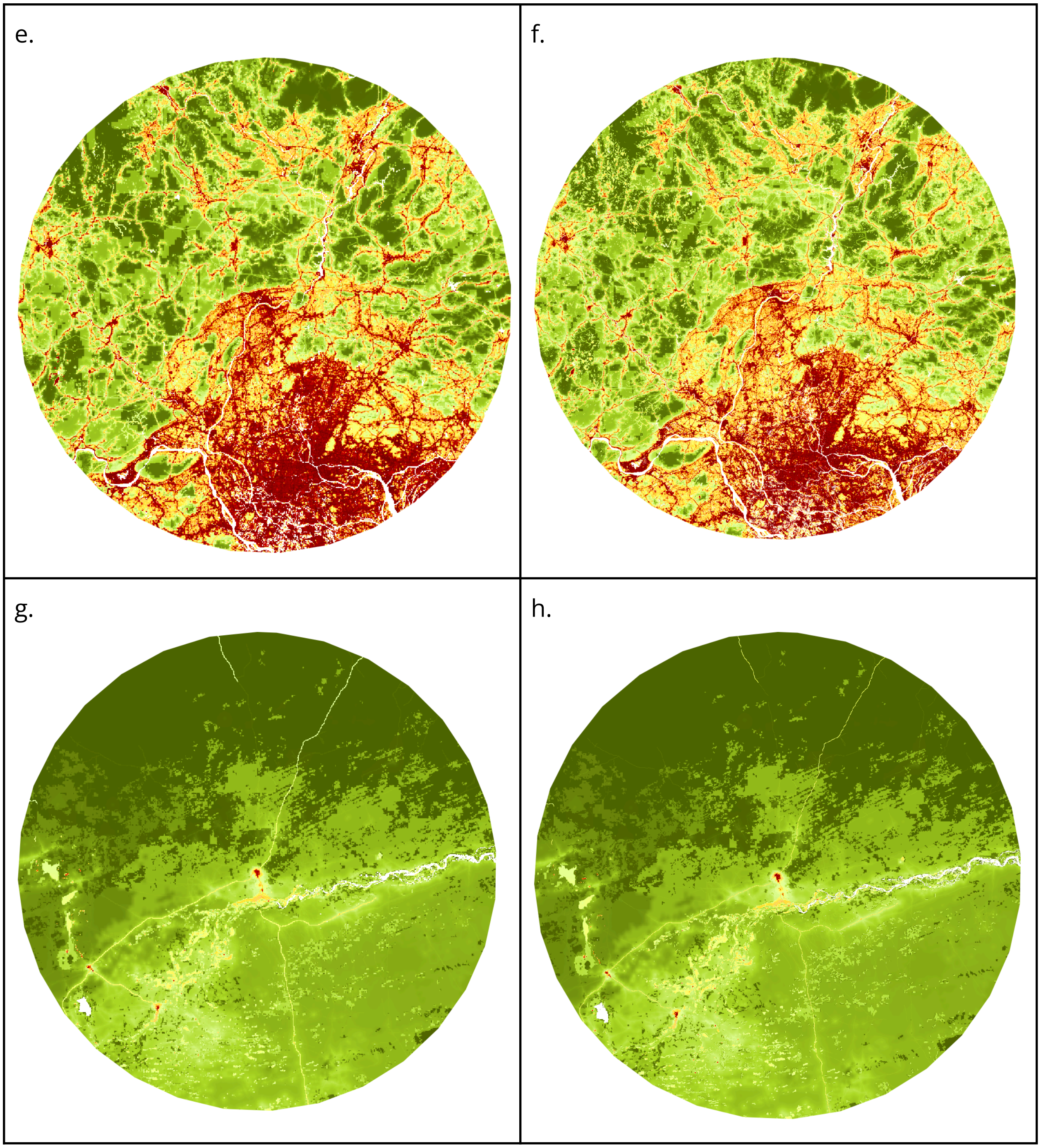
e.
f.
g.
h.

Supplemental Figure 4. Maps of human modification within 100 km radius of Asuncion, Paraguay for 1990, 2000, 2010, and 2020 (Note: datasets can be visualized at: [https://hm-30x30.projects.earthengine.app/view/hm-v3](https://hm-30x30.projects.earthengine.app/view/hm-v3).

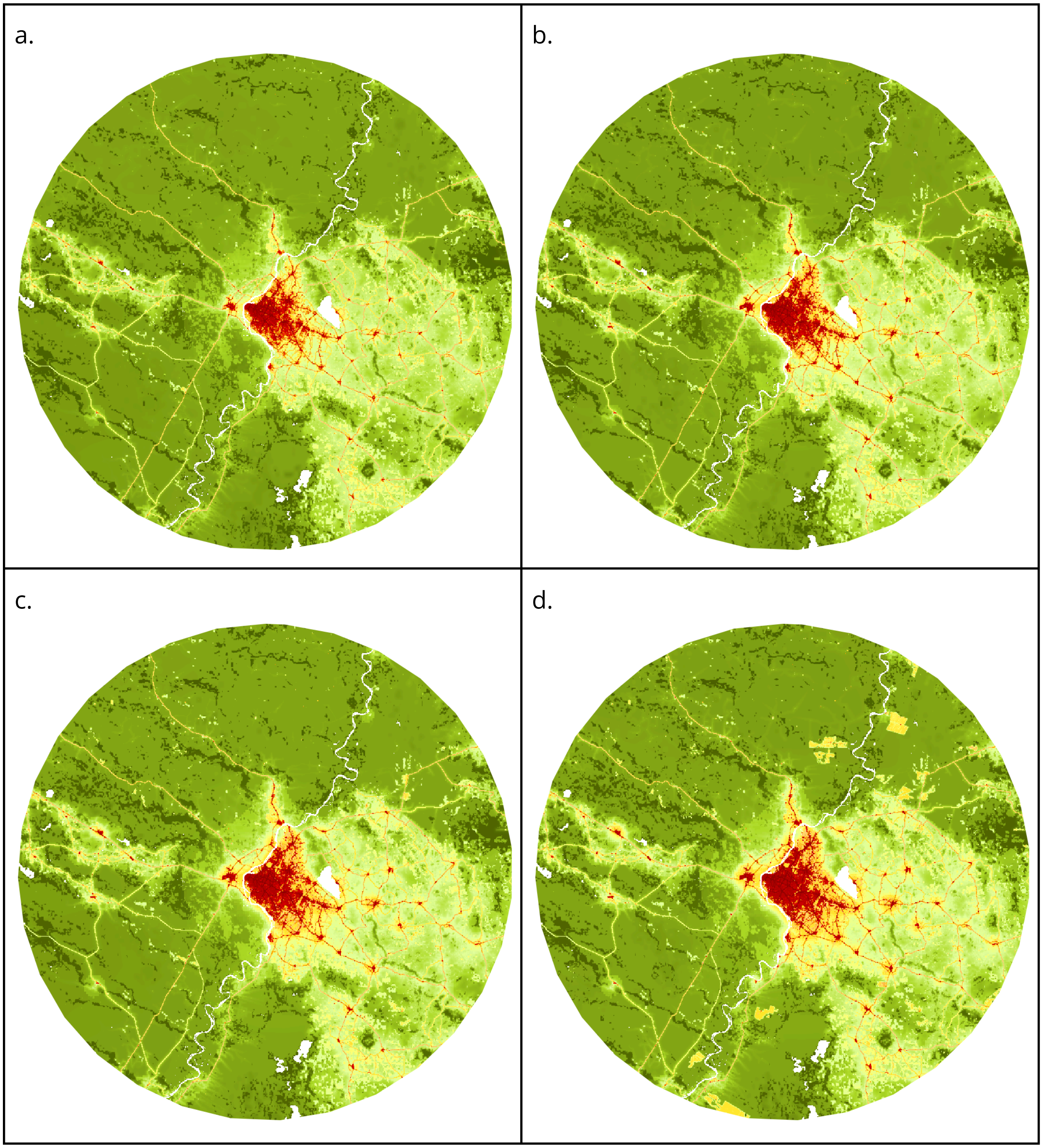

Supplemental Figure 5. Maps of human modification within 100 km radius of Calgary, Canada for 1990, 2000, 2010, and 2020. (Note: datasets can be visualized at: [https://hm-30x30.projects.earthengine.app/view/hm-v3](https://hm-30x30.projects.earthengine.app/view/hm-v3).

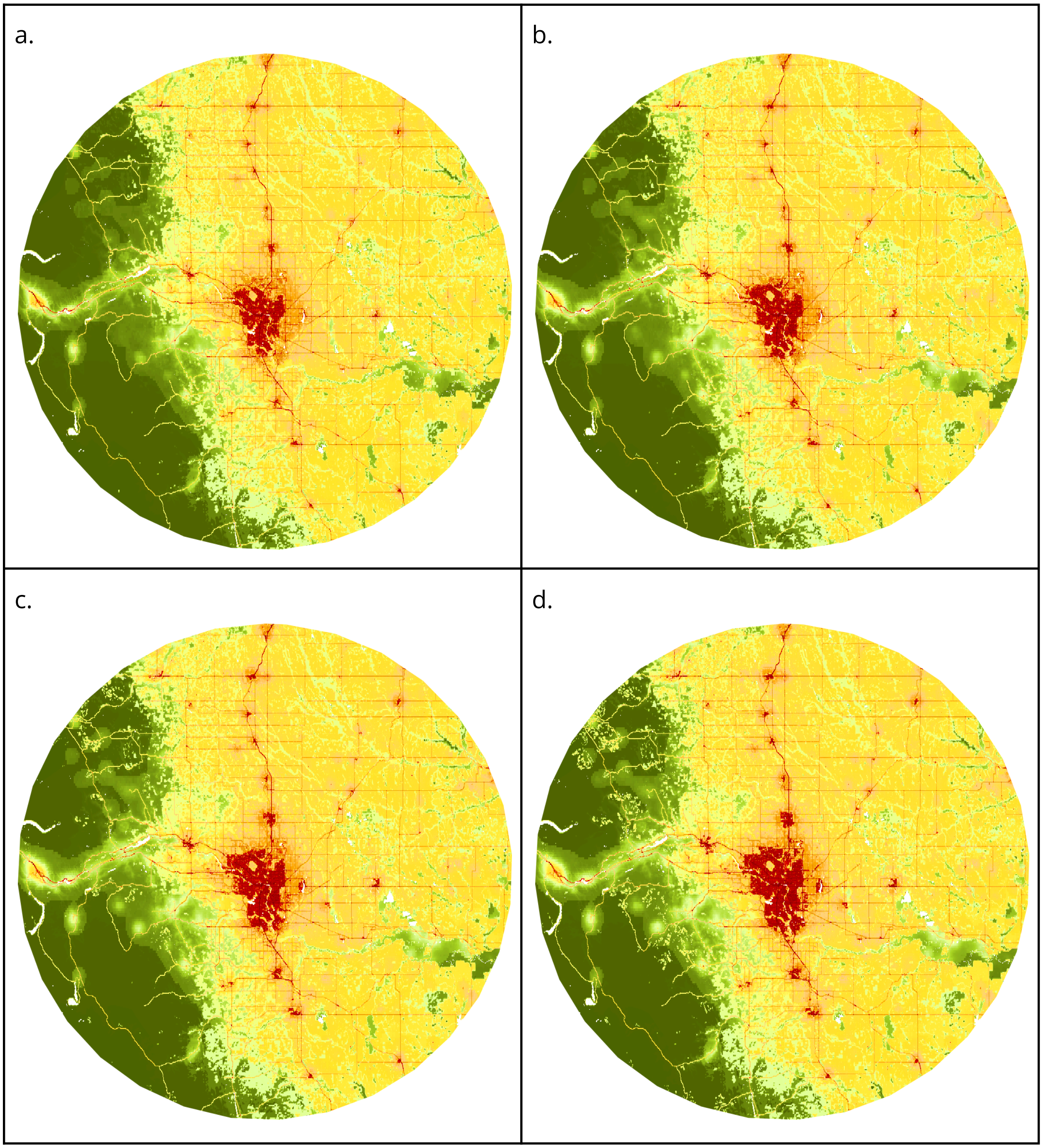

Supplemental Figure 6. Maps of human modification within 100 km radius of Qing Yuan, China for 1990, 2000, 2010, and 2020. Note: datasets can be visualized at: https://hm-30x30.projects.earthengine.app/view/hm-v3.

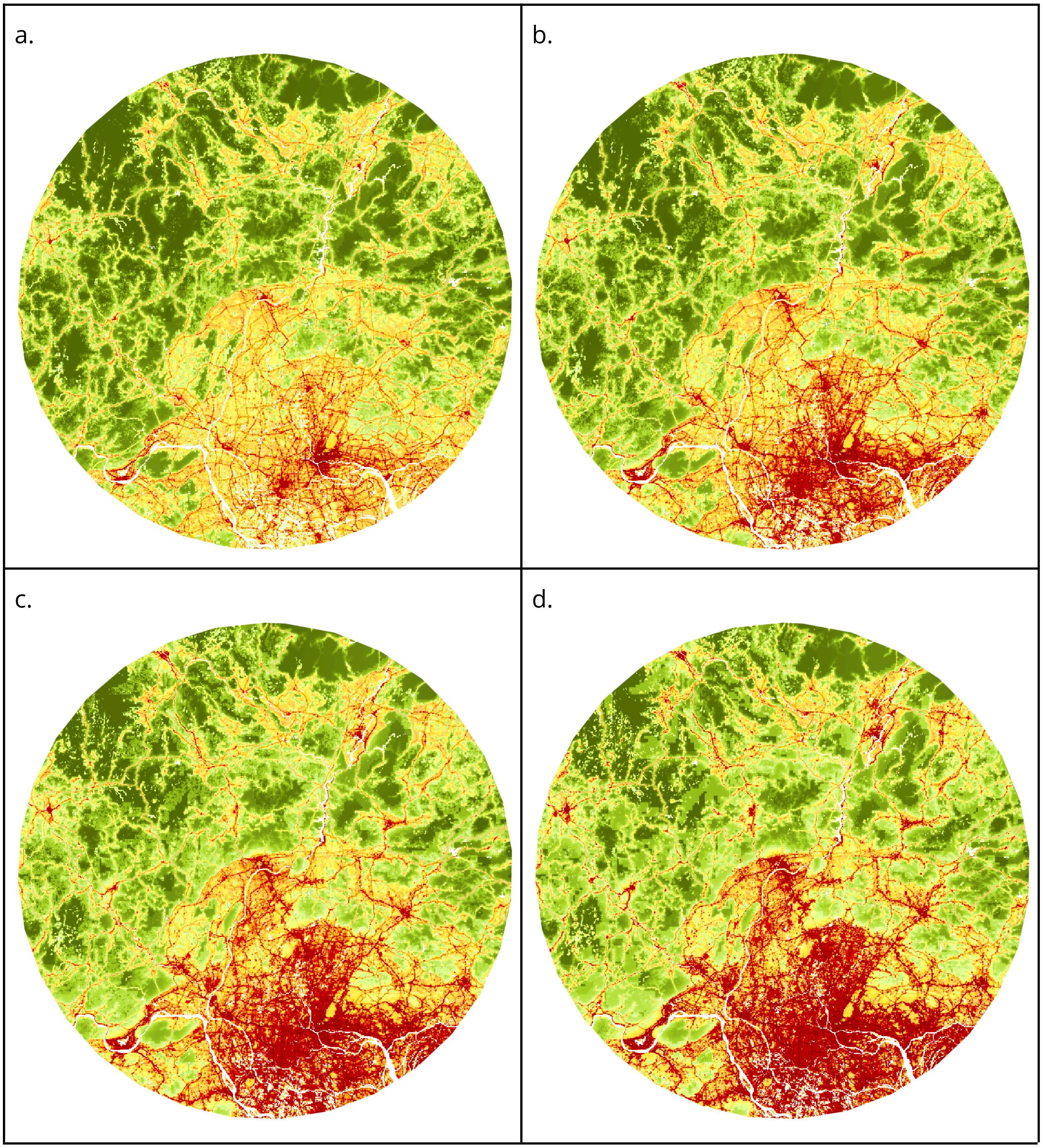

Supplemental Figure 7. Maps of human modification within 100 km radius of Timbuktu, Mali for 1990, 2000, 2010, and 2020. Note: datasets can be visualized at: [https://hm-30x30.projects.earthengine.app/view/hm-v3](https://hm-30x30.projects.earthengine.app/view/hm-v3).

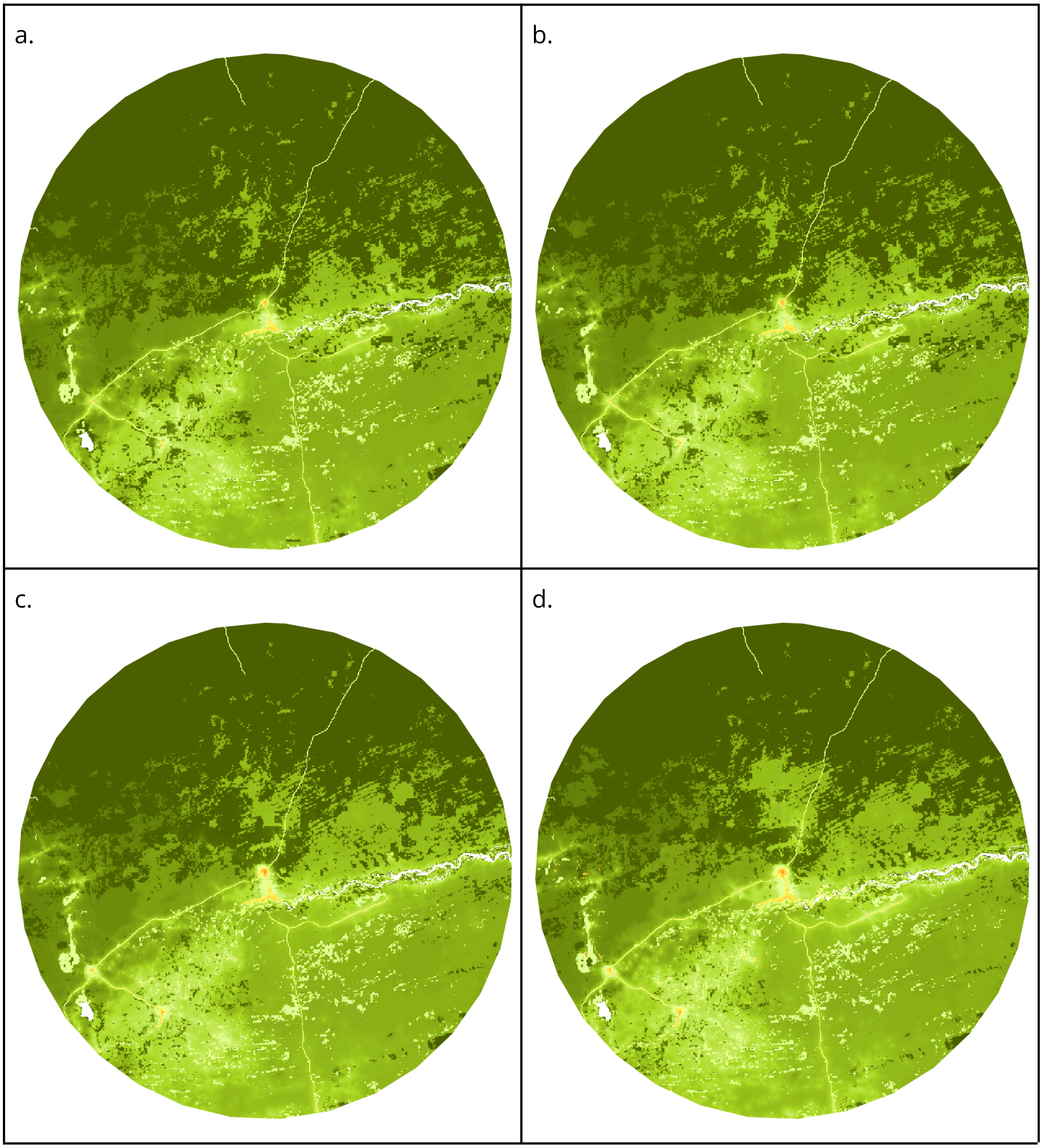

Supplemental Figure 8. Comparison of human modification and related datasets within 100 km radius of Asuncion, Paraguay for: (a) reference image from Sentinel 2; (b) HMv3 (300 m); (c) HM v2 (Theobald et al. 2020, 300 m); (d) HM v1 (Kennedy et al. 2024, 1 km); (e) HF for 2009 (Venter et al. 2016, 1 km); (f) HF for 2013 (Williams et al. 2020, 300 km); (g) HF for 2020 (Mu et al. 2022, 100 m), and (h) HF 100 m (Gassert et al. 2022). Note: datasets can be visualized globally at: [https://hm-30x30.projects.earthengine.app/view/hm-v3](https://hm-30x30.projects.earthengine.app/view/hm-v3).

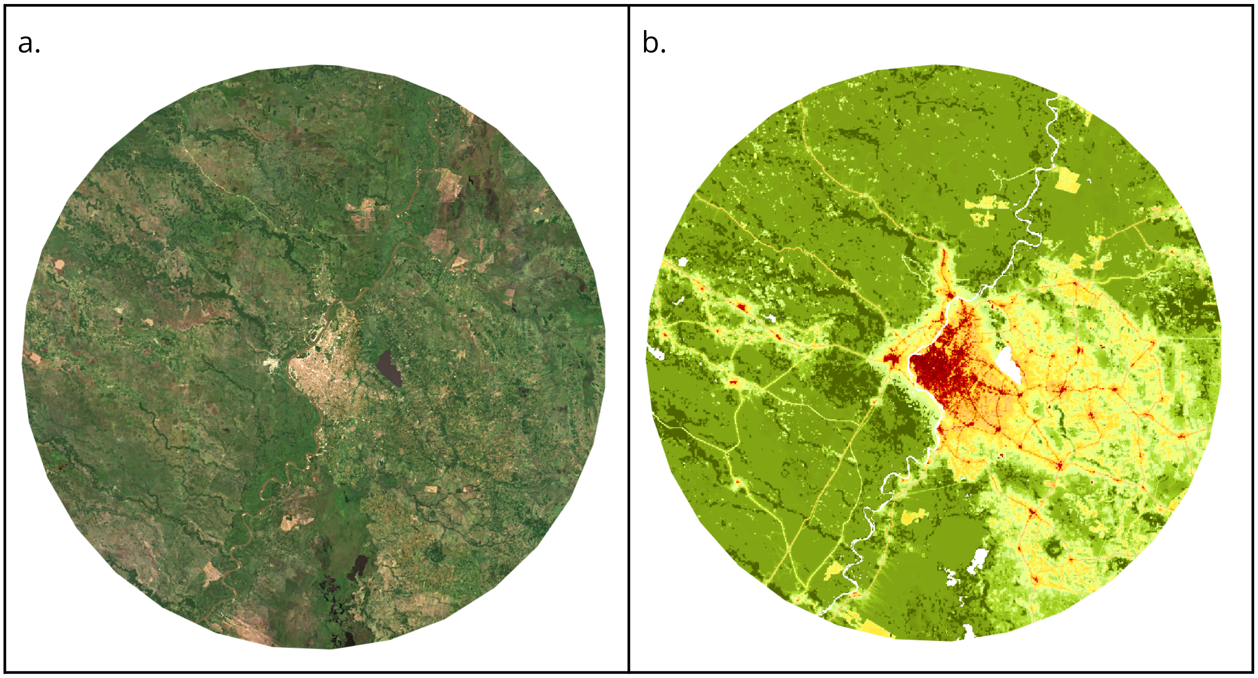

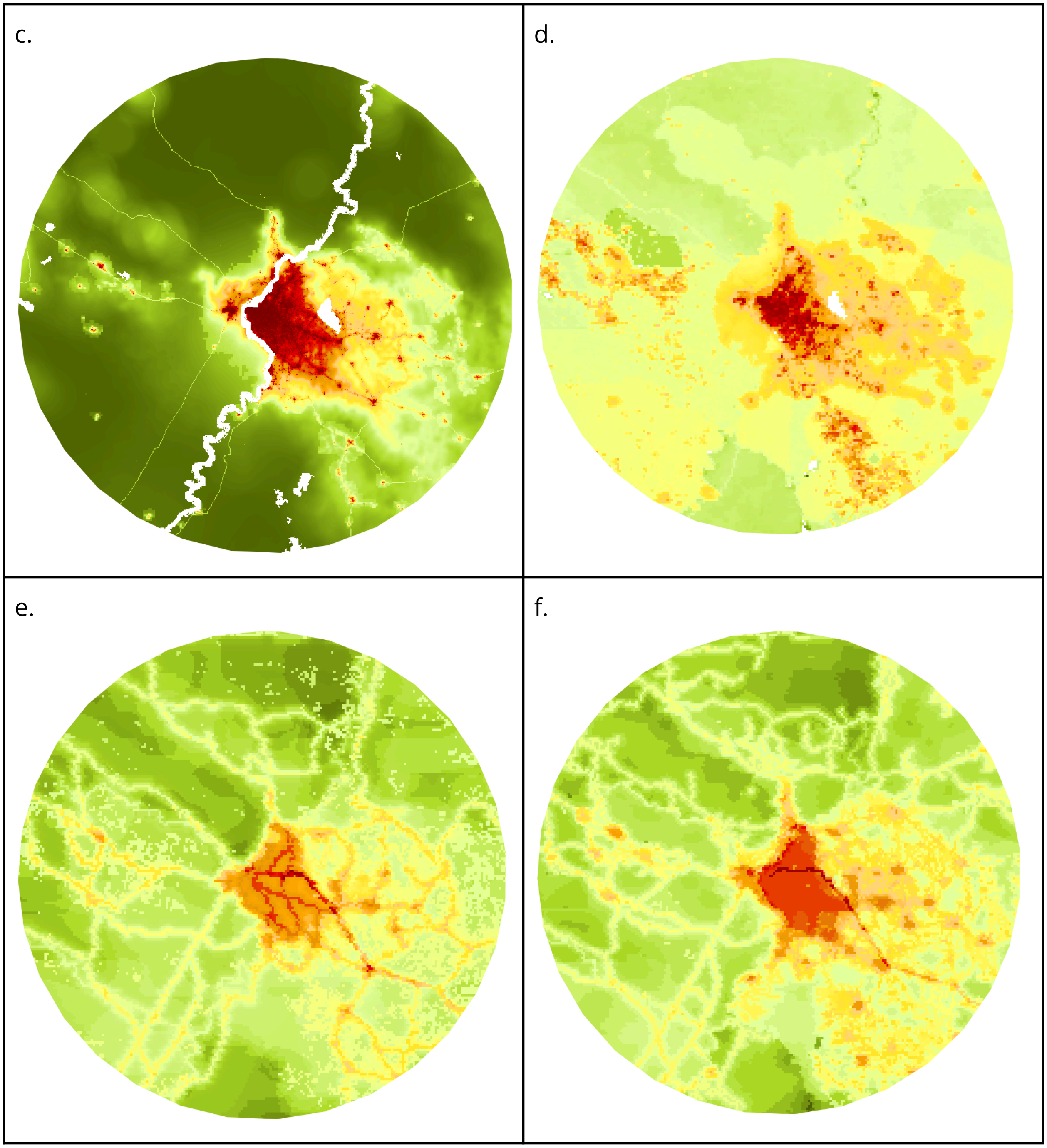
c.
d.
e.
f.

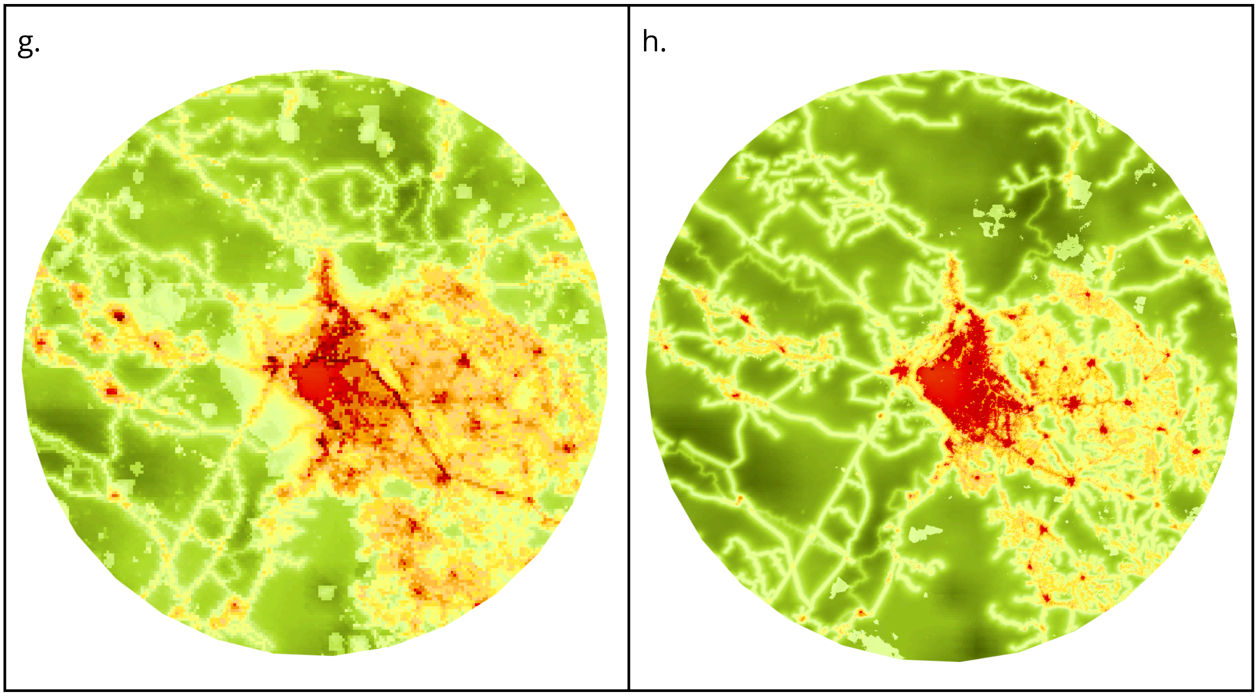
g.
h.

Supplemental Figure 9. Comparison of human modification and related datasets within 100 km radius of Calgary, Canada for: (a) reference image from Sentinel 2; (b) HMv3 (300 m); (c) HM v2 (Theobald et al. 2020, 300 m); (d) HM v1 (Kennedy et al. 2024, 1 km); (e) HF for 2009 (Venter et al. 2016, 1 km); (f) HF for 2013 (Williams et al. 2020, 300 km); (g) HF for 2020 (Mu et al. 2022, 100 m), and (h) HF 100 m (Gassert et al. 2022). Note: datasets can be visualized globally at: [https://hm-30x30.projects.earthengine.app/view/hm-v3](https://hm-30x30.projects.earthengine.app/view/hm-v3).

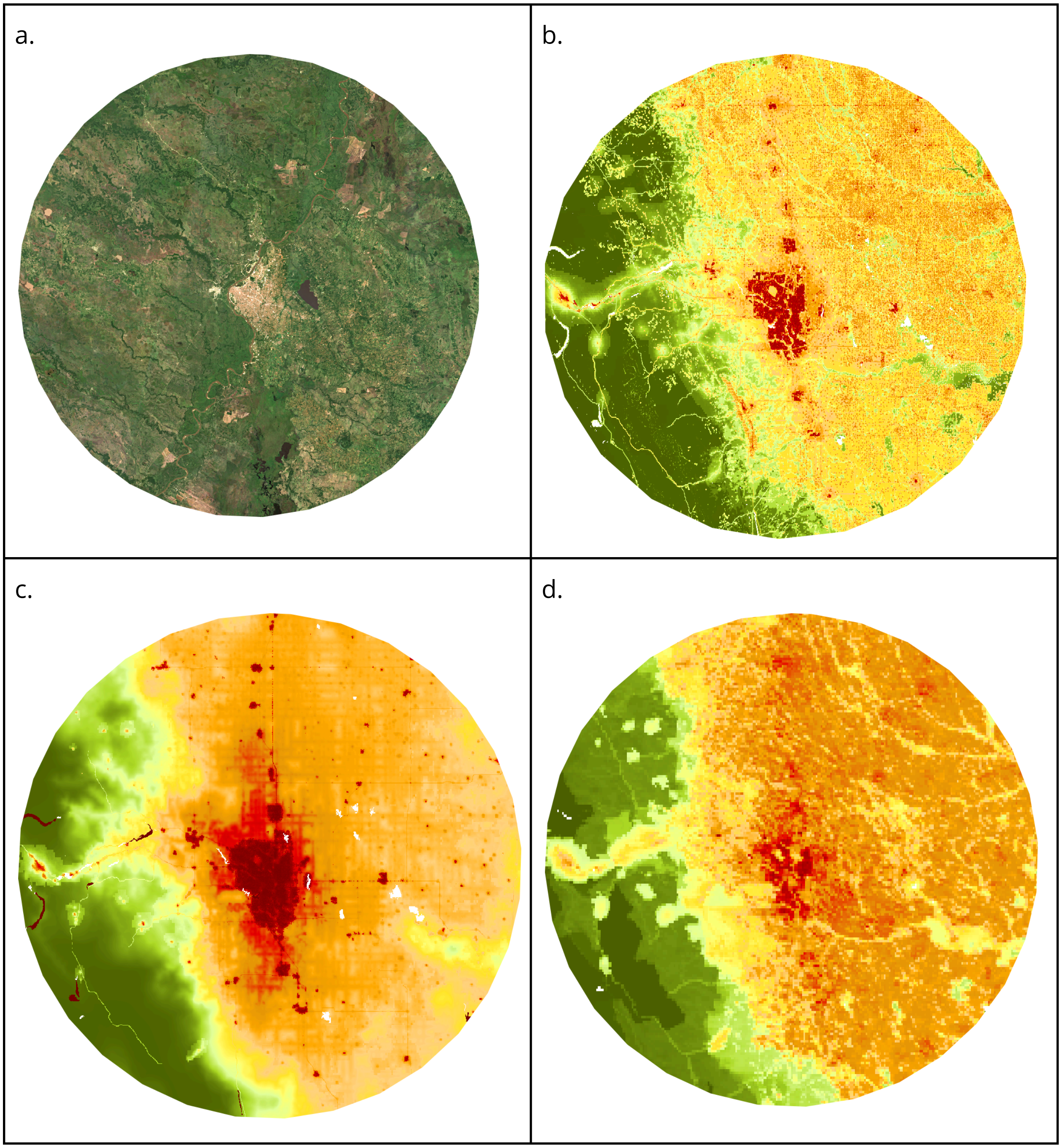

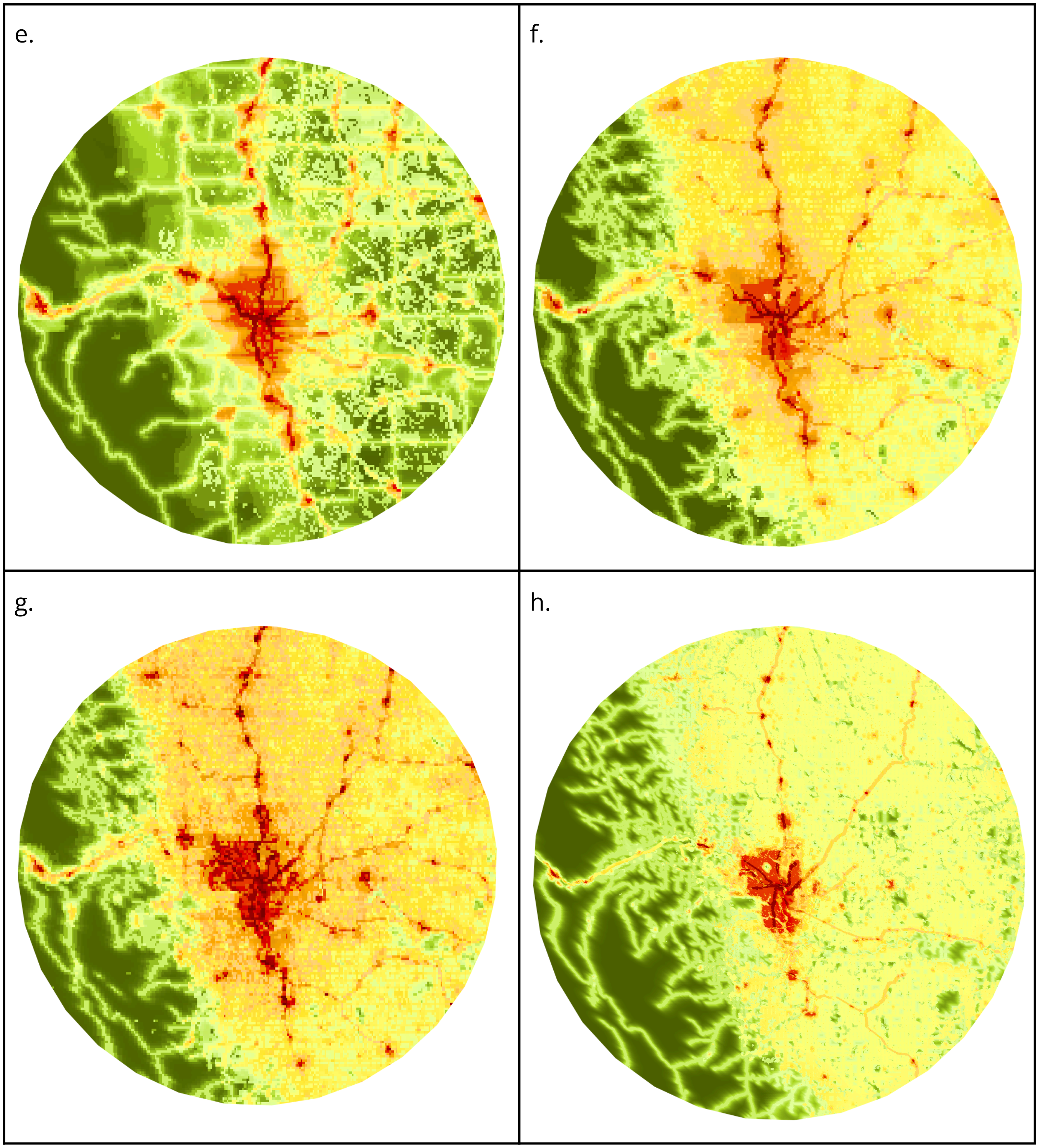
e.
f.
g.
h.

Supplemental Figure 10. Comparison of human modification and related datasets within 100 km radius of Qing Yuan, China for: (a) reference image from Sentinel 2; (b) HMv3 (300 m); (c) HM v2 (Theobald et al. 2020, 300 m); (d) HM v1 (Kennedy et al. 2024, 1 km); (e) HF for 2009 (Venter et al. 2016, 1 km); (f) HF for 2013 (Williams et al. 2020, 300 km); (g) HF for 2020 (Mu et al. 2022, 100 m), and (h) HF 100 m (Gassert et al. 2022). Note: datasets can be visualized globally at: [https://hm-30x30.projects.earthengine.app/view/hm-v3](https://hm-30x30.projects.earthengine.app/view/hm-v3).

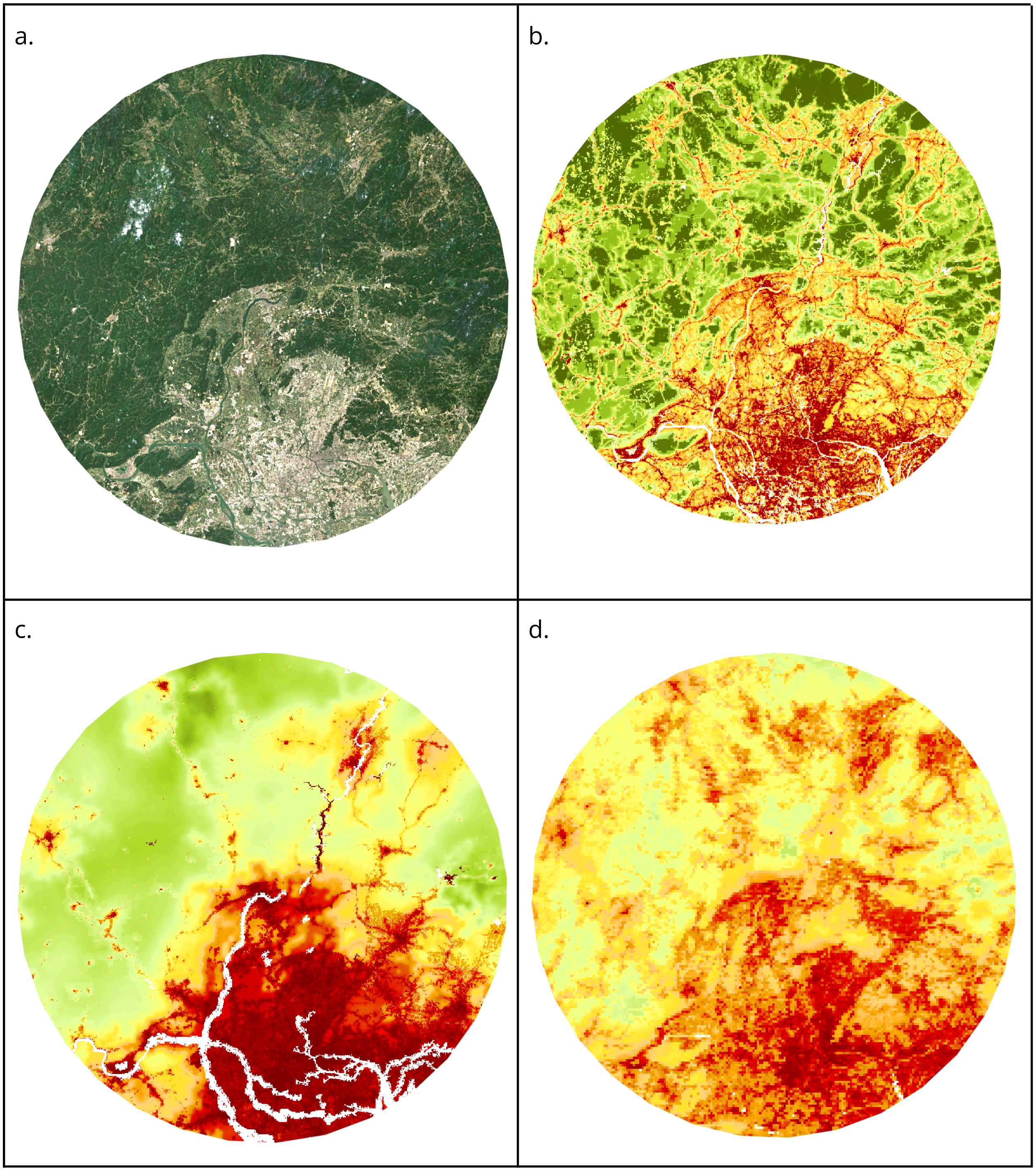

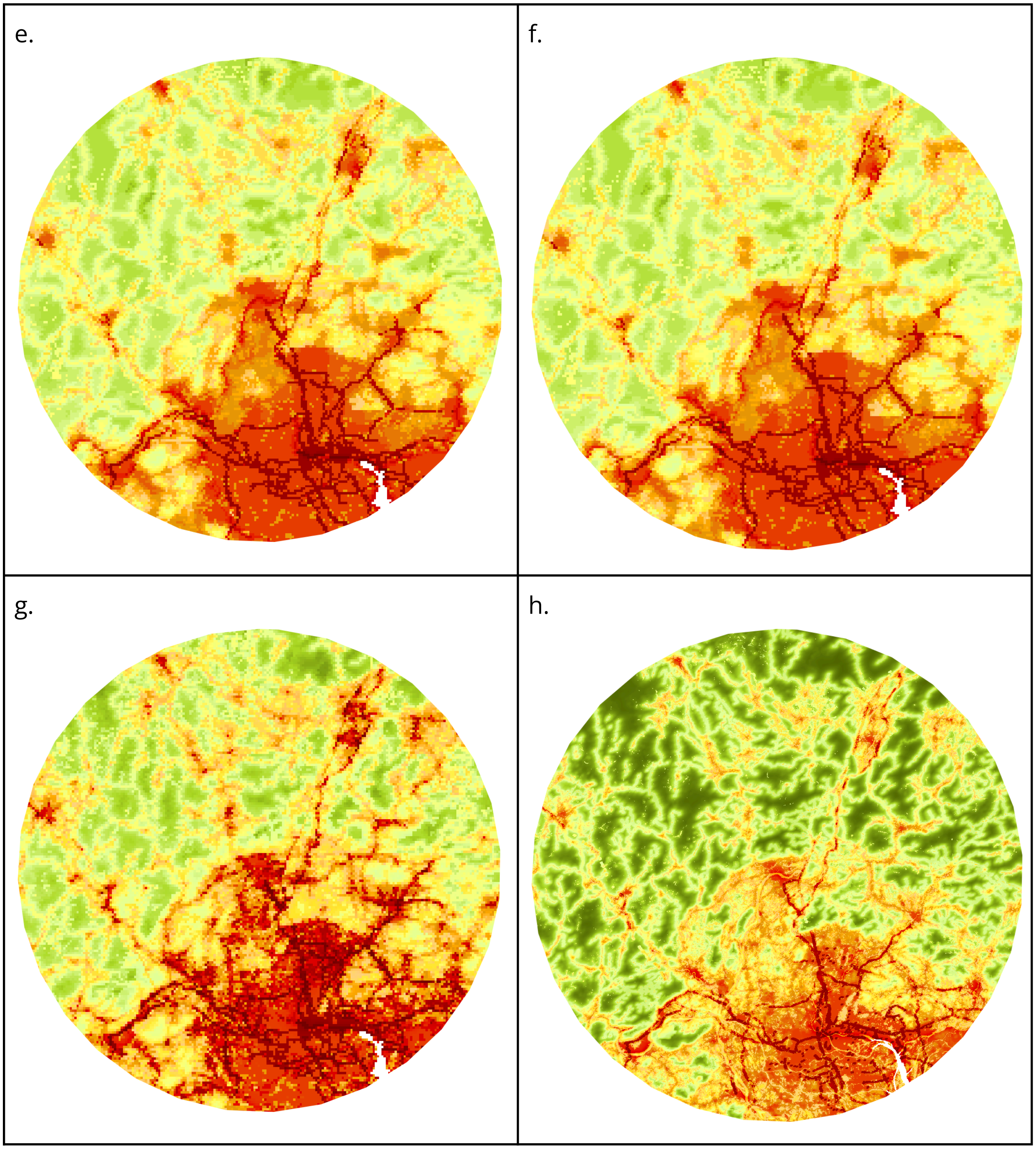
e.
f.
g.
h.

Supplemental Figure 11. Comparison of human modification and related datasets within 100 km radius of Timbuktu, Mali for: (a) reference image from Sentinel 2; (b) HMv3 (300 m); (c) HM v2 (Theobald et al. 2020, 300 m); (d) HM v1 (Kennedy et al. 2024, 1 km); (e) HF for 2009 (Venter et al. 2016, 1 km); (f) HF for 2013 (Williams et al. 2020, 300 km); (g) HF for 2020 (Mu et al. 2022, 100 m), and (h) HF 100 m (Gassert et al. 2022). Note: datasets can be visualized globally at: [https://hm-30x30.projects.earthengine.app/view/hm-v3](https://hm-30x30.projects.earthengine.app/view/hm-v3).

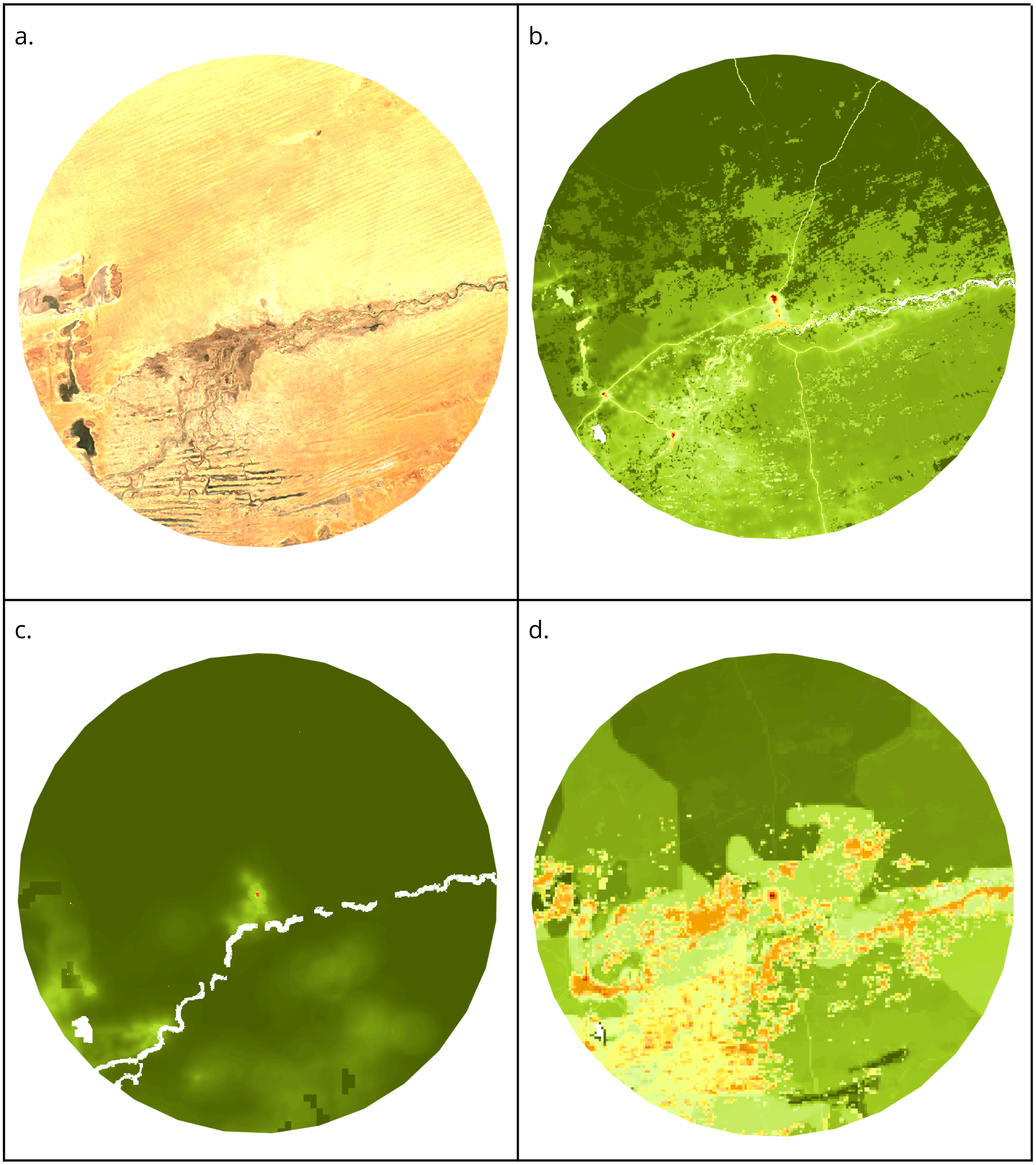

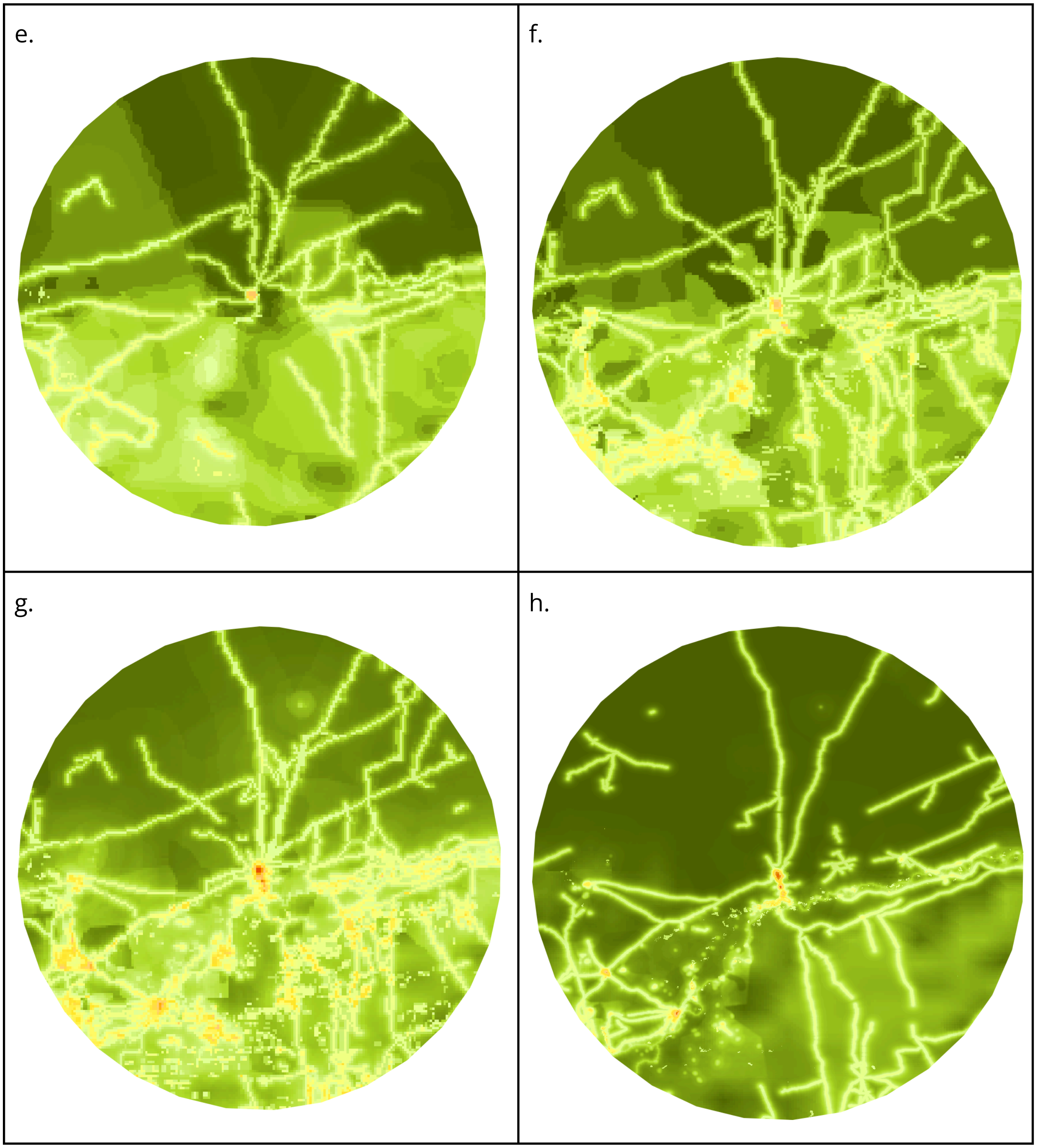
e.
f.
g.
h.

Supplemental Figure 12. Map showing which of the threat classes was “dominant” (i.e. had the highest *H* value, that exceeded 0.01), globally (top) and for the central part of North America.

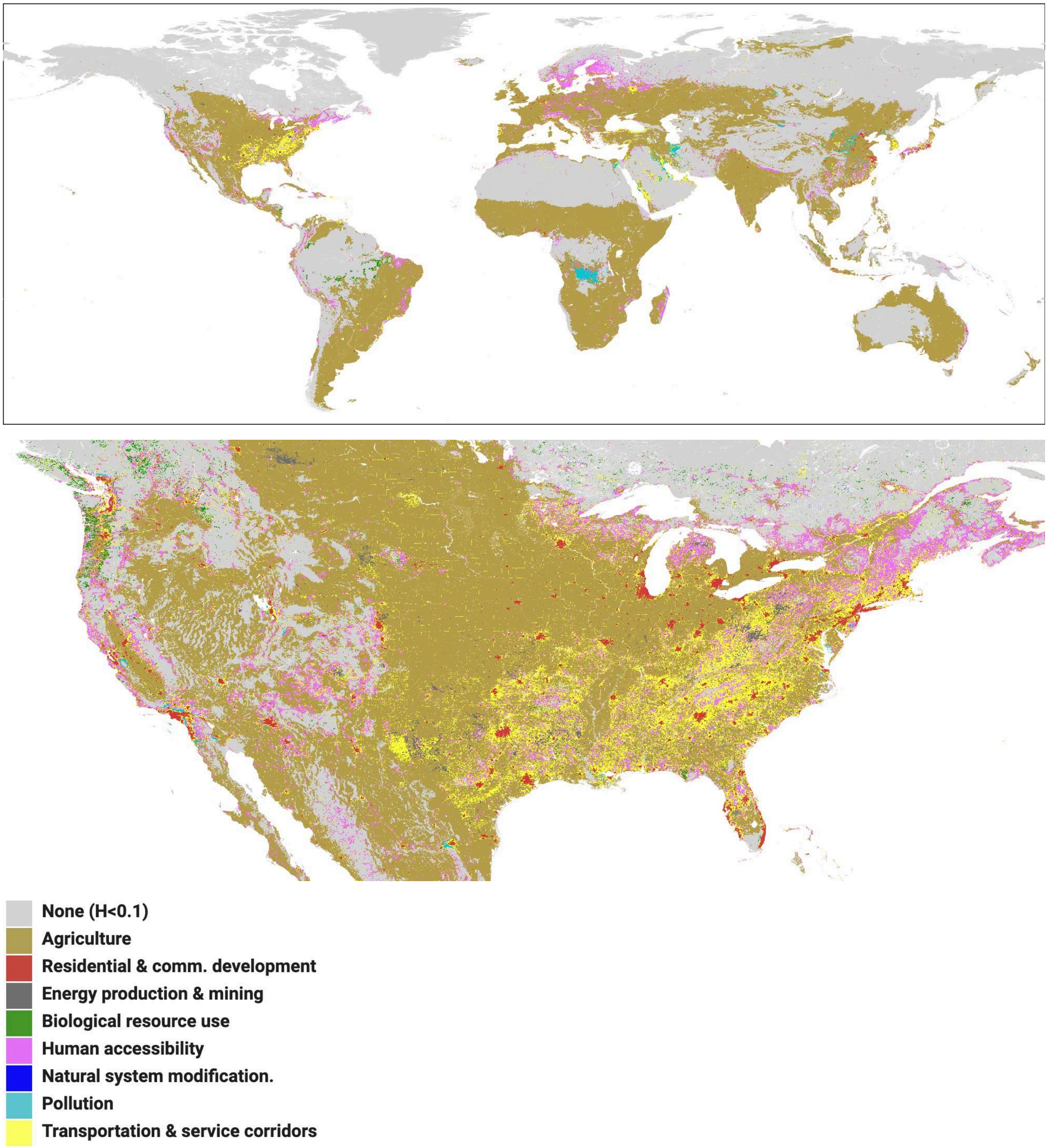

Supplemental Figure 13. Scattergram of countries depicting the percent protected vs. the increase in human modification from 1990-2020.

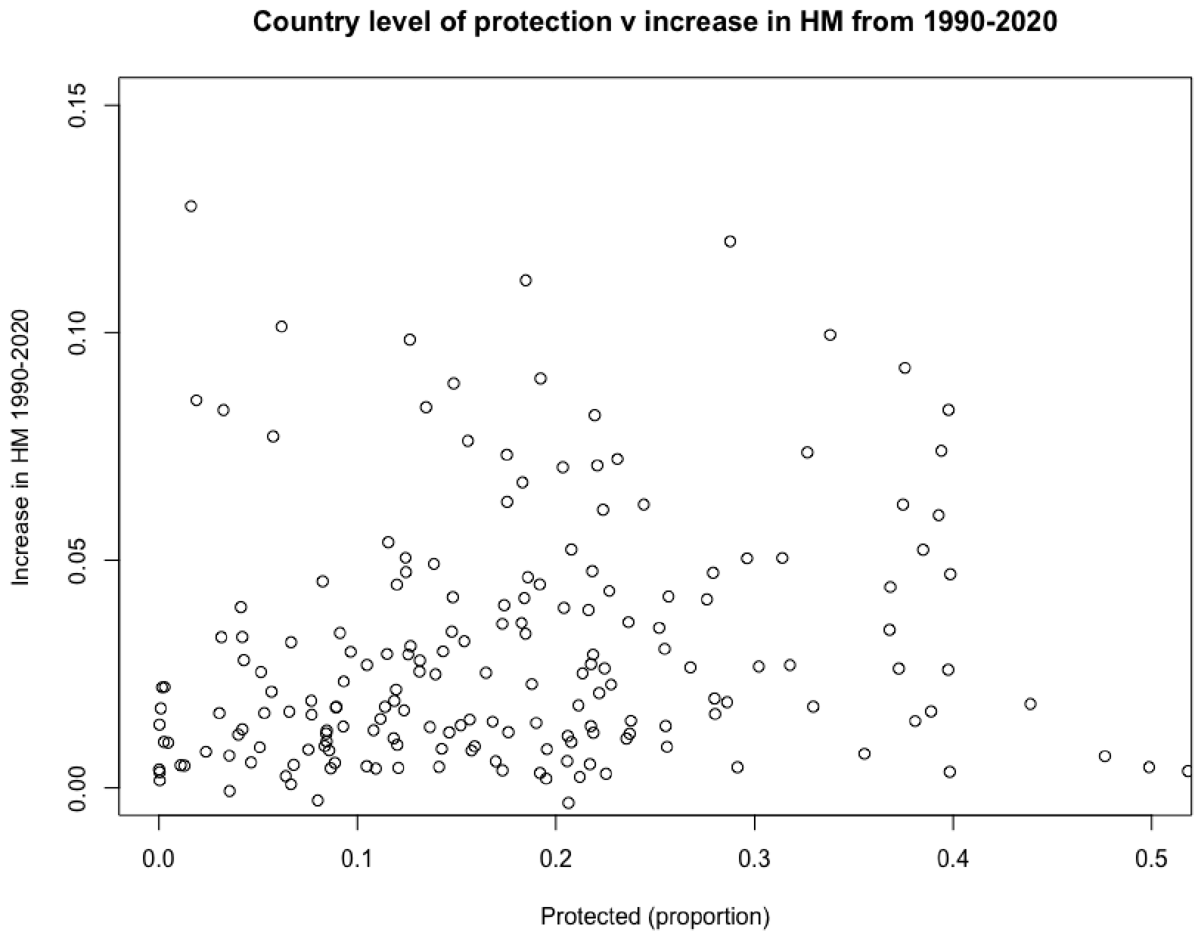

Supplemental Figure 14. Scattergram of ecoregions depicting the proportion protected vs. the increase in human modification from 1990-2020.

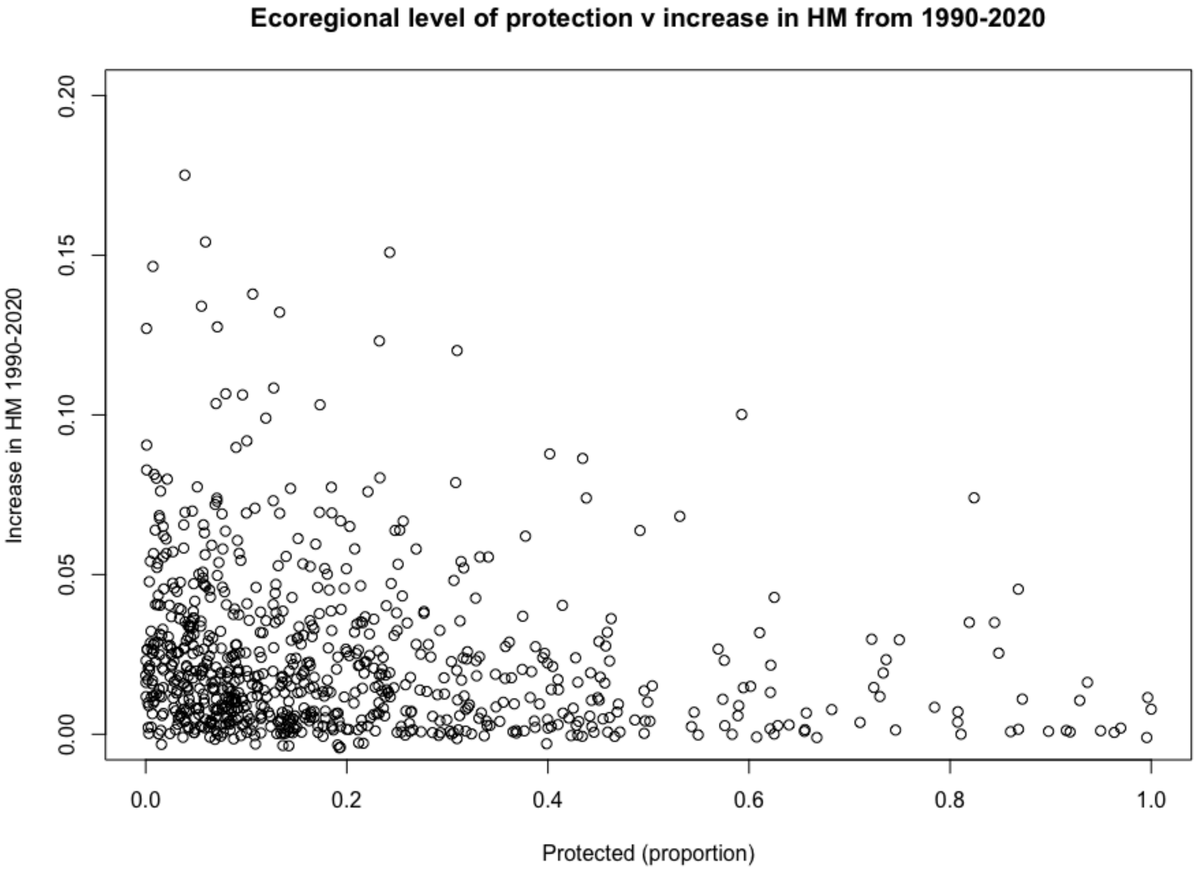

Supplemental Figure 15. A bivariate map showing the percentage protected in 2024 vs. the change in human modification ($HM_{2020}$ - $HM_{1990}$).

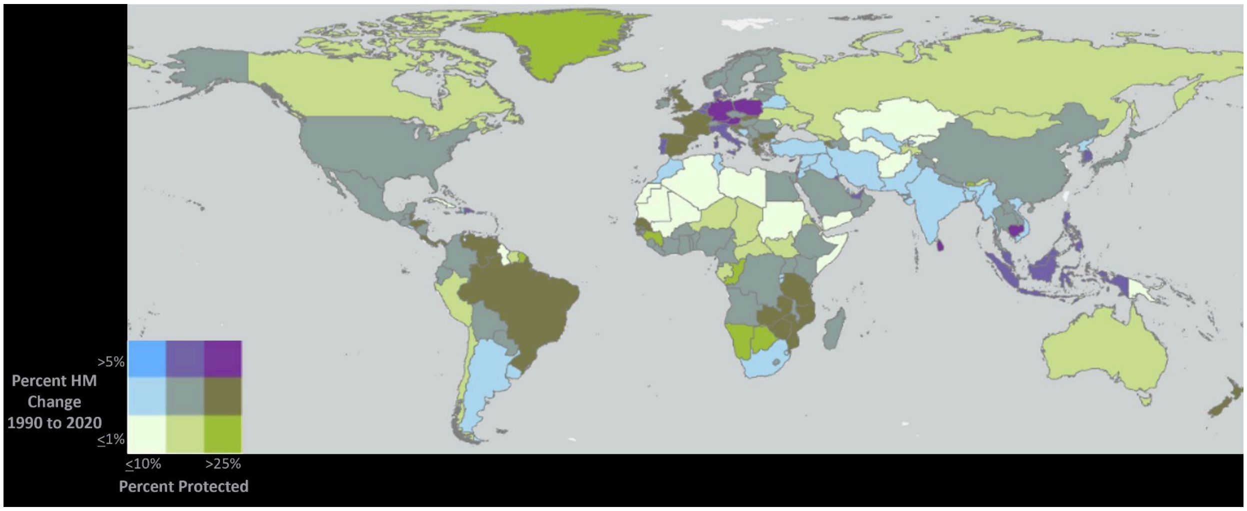

Supplementary Table 1. Derivation of human modification intensity values ($I_s$) by land use classification from the landscape development intensity index (LDI), calculated as max-normalized empower densities[30].

| Land use | LDI | $I_s$ |
|---|---|---|
| Natural system | 1 | 0.000 |
| Natural open water | 1 | 0.000 |
| Pine plantation | 1.58 | 0.064 |
| Rec/open space low-intensity | 1.83 | 0.092 |
| Woodland pasture (w/livestock) | 2.02 | 0.113 |
| Improved past (w/o livestock) | 2.77 | 0.197 |
| Improved past - low intensity (w/livestock) | 3.41 | 0.268 |
| Citrus plantation | 3.68 | 0.298 |
| Improved pasture high-intensity (w/livestock) | 3.74 | 0.304 |
| Row crops | 4.54 | 0.393 |
| Single family low density | 6.9 | 0.656 |
| Rec/open space high-intensity | 6.92 | 0.658 |
| Ag high intensity | 7 | 0.667 |
| Single family medium density | 7.57 | 0.730 |
| Single family high density | 7.55 | 0.728 |
| Mobile home medium density | 7.7 | 0.744 |
| Highway 2 lane | 7.81 | 0.757 |
| Low intensity commercial | 8 | 0.778 |
| Institutional | 8.07 | 0.786 |
| Highway 4 lane | 8.28 | 0.809 |
| Mobile home high density | 8.29 | 0.810 |
| Industrial | 8.32 | 0.813 |
| Multi-family res low-rise | 8.66 | 0.851 |
| High intensity commercial | 9.18 | 0.909 |
| Multi-family res high-rise | 9.19 | 0.910 |
| Commercial/business district, 2 stories | 9.42 | 0.936 |
| Commercial/business district, 4 stories | 10 | 1.000 |

Supplementary Table 2. ESA CCI Land cover classes examined ("class name") and used to define pattern of grazing patches, andused to estimate grazing footprint from the livestock density for bovines, goats, and sheep (LU/km2) from the Gridded Livestock of the World[51] (GLW). LU= livestock units. Grazing classes were first identified by identifying cover classes typically associated with grazing from a biological perspective (column "biological"). Adjusted classes (column "adjusted") reflect subsequent evaluation based on the empirical distribution of livestock grazing for each class, and the percent of land in each class, as well as visual inspection of the spatial overlapping pattern of the grazing density against the land cover map.

| | | **Livestock units/km2 value at percentile** | | | **Grazing classes** | | |
|---|---|---|---|---|---|---|---|
| **Value** | **Class name** | **50%** | **90%** | **95%** | **Biological** | **Adjusted** | **% of weighted area** |
| 0 | No data | | | | | | |
| 10 | Cropland, rainfed | 27.9 | 156.0 | 220.0 | | | 11.765% |
| 11 | Cropland, rainfed herbaceous | 11.7 | 78.1 | 110.3 | | | 9.662% |
| 12 | Cropland, rainfed tree or shrub | 23.0 | 103.8 | 168.3 | | | 0.242% |
| 20 | Cropland, irrigated | 92.0 | 340.0 | 444.0 | | | 4.438% |
| 30 | Mosaic cropland (>50%) | 23.5 | 103.4 | 135.6 | Y | Y | 4.801% |
| 40 | Mosaic cropland (<50%) | 23.3 | 87.2 | 135.7 | Y | Y | 4.238% |
| 50 | Tree (>15%), broadleaved, evergreen | 2.1 | 27.7 | 51.8 | | | 6.860% |
| 60 | Tree (>15%), broadleaved, deciduous | 5.9 | 44.4 | 44.4 | | Y | 4.277% |
| 61 | Tree (>40%), broadleaved, deciduous | 11.9 | 35.7 | 51.6 | | | 0.678% |
| 62 | Tree (>15-40%), broadleaved, deciduous | 11.8 | 59.8 | 99.9 | | Y | 3.628% |
| 70 | Tree (>15%), needleleaved, evergreen | 1.0 | 17.9 | 34.0 | | | 2.607% |
| 71 | Tree (>40%), needleleaved, evergreen | 0.3 | 0.3 | 0.3 | | | 0.086% |
| 72 | Tree (>15-40%), needleleaved, evergreen | 0.2 | 0.3 | 1.4 | | | 0.000% |
| 80 | Tree (>15%, needleleaved, deciduous | 0.8 | 0.8 | 0.8 | | | 0.412% |
| 81 | Tree (>40%), needle leaved, deciduous | 51.2 | 183.1 | 249.3 | | Y | 0.006% |
| 82 | Tree (>15-40%), needleleaved, deciduous | 127.0 | 127.0 | 127.0 | | Y | 0.000% |
| 90 | Tree, mixed leaf type | 3.0 | 3.0 | 3.0 | | | 0.724% |
| 100 | Mosaic tree/shrub (>50%) | 7.7 | 45.4 | 77.6 | Y | Y | 2.805% |
| 110 | Mosaic herbaceous (>50%) | 11.6 | 67.9 | 107.7 | Y | Y | 1.011% |
| 120 | Shrubland | 8.5 | 44.6 | 77.5 | Y | Y | 10.135% |

| | | | | | | | |
|---|---|---|---|---|---|---|---|
| 121 | Shrubland, evergreen | 9.9 | 34.0 | 54.0 | Y | Y | 0.179% |
| 122 | Shrubland, deciduous | 2.2 | 35.9 | 67.7 | Y | Y | 1.147% |
| 130 | Grassland | 5.4 | 71.3 | 103.6 | Y | Y | 13.217% |
| 140 | Lichens and mosses | 0.3 | 0.3 | 0.3 | | | 0.094% |
| 150 | Sparse vegetation (<15%) | 2.4 | 23.0 | 55.3 | Y | Y | 4.303% |
| 151 | Sparse tree (<15%) | 17.1 | 23.8 | 23.8 | | | 0.000% |
| 152 | Sparse shrub (<15%) | 41.9 | 106.0 | 122.1 | Y | Y | 0.066% |
| 153 | Sparse herbaceous (<15%) | 42.0 | 110.0 | 126.2 | Y | Y | 0.469% |
| 160 | Tree, flooded | 1.0 | 1.0 | 1.0 | | | 0.238% |
| 170 | Tree, flooded saline | 2.4 | 52.0 | 83.3 | | | 0.110% |
| 180 | Shrub/herbaceous flooded | 6.1 | 43.6 | 77.4 | Y | Y | 0.923% |
| 190 | Urban areas | 5.9 | 103.7 | 167.2 | | | 0.859% |
| 200 | Bare | 2.4 | 35.7 | 51.8 | | | 8.638% |
| 201 | Bare areas | 0.6 | 74.0 | 122.0 | | | NA |
| 202 | Bare areas | 5.4 | 21.6 | 41.4 | | | NA |
| 210 | Water | 1.3 | 35.8 | 76.0 | | | 1.138% |
| 220 | Permanent snow & ice | 0.3 | 4.9 | 6.9 | | | 0.107% |

Supplementary Table 3. The area and percentage of lands dominated by threat classes.

| Threat class | Percentage | Area (M km2) |
|---|---|---|
| HM<0.01 | 46.98% | 61.74 |
| Agriculture | 39.60% | 52.05 |
| Residential & commercial devel. | 1.59% | 2.10 |
| Energy production & mining | 0.22% | 0.29 |
| Biological resource use | 1.00% | 1.32 |
| Human accessibility | 5.10% | 6.70 |
| Natural process modification | 0.04% | 0.05 |
| Pollution | 0.59% | 0.77 |
| Transportation & service corridors | 4.87% | 6.40 |

Supplementary Table 4. Time-series of mean human modification values (HMv3 300 m) from 1990 to 2020 for the four example areas: Asuncion, Paraguay; Calgary, Canada; Qing Yuan, China; and Timbuktu, Mali.

| **HM v3 300 m** | **Asuncion, Paraguay** | **Calgary, Canada** | **Qing Muay, China** | **Timbuktu, Mali** |
|---|---|---|---|---|
| **1990** | 0.1081 | 0.2765 | 0.2416 | 0.0470 |
| **1995** | 0.1129 | 0.2800 | 0.2599 | 0.0472 |
| **2000** | 0.1157 | 0.2834 | 0.2812 | 0.0480 |
| **2005** | 0.1163 | 0.2852 | 0.3011 | 0.0489 |
| **2010** | 0.1220 | 0.2890 | 0.3252 | 0.0531 |
| **2015** | 0.1257 | 0.2927 | 0.3422 | 0.0543 |
| **2020** | 0.1289 | 0.2947 | 0.3482 | 0.0571 |

Supplementary Table 5. Overview comparison of human modification to other human pressure methods.

| Name | Human modification v3 (2022) | Human modification 2017 (HM2017) | Human modification 2016 (HM2016) | Low impact areas (LIA) | Human footprint (HFP-100) | Human footprint (HF 2000-2013) | Human footprint (HF 1993-2009) | Human Influence Index (HII) |
|---|---|---|---|---|---|---|---|---|
| **Citation** | (this paper) | Theobald et al. (2020) | Kennedy et al. (2019); Theobald (2013) | Jacobsen et al. (2019) | Gassert et al. (2023) | Williams et al. (2020) | Venter et al. (2016) | Sanderson et al. (2002) |
| **Conceptual framework** | IUCN Direct Threats Classification v2 (Salafsky et al., 2008); Theobald et al. (2020) | Direct Threats Classification v2 (Salafsky et al., 2008) | Direct Threats Classification v2 (Salafsky et al., 2008) | Classed overlays (as primarily modified or used by humans) | Based on Sanderson et al. (2002) framework, a "simpler" approach than HM | Based on Venter et al. (2016). | Based on Sanderson et al. (2002) framework | Ad hoc |
| **Footprint/intensity** | Intensity values based on land development index (LDI; Brown and Vivas, 2005); Kennedy et al. 2019 SI; McBride et al. 2012 elicitation procedure | Intensity values based on land development index (LDI; Brown and Vivas, 2005); Kennedy et al. 2019 SI; McBride et al. 2012 elicitation procedure | Intensity values based on land development index (LDI; Brown and Vivas, 2005); Kennedy et al. 2019 SI; McBride et al. 2012 elicitation procedure | - | Authors estimated 0-10 index for each pressure. | Authors estimated 0-10 index for each pressure | Authors estimated 0-10 index for each pressure | Authors estimated 0-10 index for each pressure |
| **Resolution** | 90, 300 m | 0.3 km | 1 km | 1 km | 100 m | 1 km | 1 km | 1 km |
| **Year(s)** | ~2022<br>1990-2020 (5 years) | ~2017<br>1990-2015 | ~2016 | ~2016 | 2015-2019, 2020 (v1.2 2017-2021) | 2013<br>2000-2013 | 2009<br>1993-2009 | ~2000 |
| **Threat classes** | | | | | | | | |

| | | | | | | | | |
|---|---|---|---|---|---|---|---|---|
| **Agriculture** | ESA LCCS land cover (300 m) ESA LCCS land cover (100 m) Global cropland database (30 m) Plantations (30 m) Grazing (GLW v2; 1km) | Cropland & pastureland for 1990, 2015 (ESA CCI; 300 m) and cropland intensity (GLS; 1 km); Unified Cropland Layer (UCL; 1 km); grazing (GLW; 10 km; 1 | Unified Cropland Layer (UCL; 1 km); grazing (GLW v2; 1 km; LU km-2 | ESA LCCS land cover: urban & cropland Grazing Gridded livestock World and LandScan Population (>1->16 people/km2) | Cropland from Copernicus CGLS-LC100 (for 2015-2019) and ESRI 2020 Global Land UseLand Cover (for 2020) Pastureland from Ramankutty et al. 2008 (10 km resolution* not 1 km downsampled as stated). | Cropland (ESA CCI land cover 300 m for 2000 and 2013) Pastureland for 2000 from Ramankutty et al. (2008), 10 km resolution, downsampled to 1 km) | Cropland (University of Maryland for 1990 and GlobCover for 2009); pastureland (2000), 10 km | Land cover 1993 |
| **Residential & commercial development** | Global Human Settlement Layer vR2022A (GHSL; 100 m); GHSL 10 m; Microsoft Building Footprints Impervious surface (10 m) Land cover (10 m) | GHSL vR2019 300 m | GHSL v2016 1 km; Population density (GPW v4 2015; 1 km) | Nighttime lights (VIIRS > 0) | Built-up class from Copernicus CGLS-LC100 (for 2015-2019) and ESRI 2020 Global Land UseLand Cover (for 2020) Human population density from WorldPop 100 m resolution (not 100 m2 as erroneously stated) | Nighttime lights (DMSP OLS >20; 1 km; 1994–2012); Population density (CIESIN v3; 4 km; 1990, 2010) | Nighttime lights (DMSP OLS >20; 1 km; 1994–2012); population density (CIESIN v3; 4 km; 1990, 2010) | Gridded Population of the World (1995) Land cover 1993 Nightlights (DMSP/OLS; 1994) |
| **Energy production & mining** | Oil & gas production (gas flares; DMSP OLS and VIIRS); renewable and nonrenewable power plants | Oil & gas production (gas flares; DMSP OLS and VIIRS); renewable and nonrenewable power plants | Oil & gas wells, wind turbines, mines (OSM, 2016) | - | | - | - | - |

| | | | | | | | | |
|---|---|---|---|---|---|---|---|---|
| | (WRI); large mining operations (Maus) | (WRI); large mining operations (S&P) | | | | | | |
| **Biological resource use** | Forest loss (Hansen et al., 2013; 0.03–1 km; 2000–2023) | Forest loss (Hansen et al., 2013; Curtis et al., 2018; 0.03–1 km; 2000–2017) | - | Forest loss (Hansen et al., 2013); Burned area extent (MODIS) | - | - | - | - |
| **Human accessibility (intrustions)** | Human intrusion (HUE, 1990–2020; 1 km) - accessibility along transportation network and navigable rivers/coastlines, weighted by settlement population | Human intrusion (HUE, 1990–2015; 1 km) - accessibility along transportation network and navigable rivers/coastlines, weighted by settlement population | - | - | Navigable waterways (using VMAP) | Navigable waterways (using VMAP) | Navigable waterways (using VMAP) | Navigable waterways (using VMAP) |
| **Natural system mod.** | Reservoirs (GRanD, 1990–2017; 0.03 km; GeoDAR 30 m; RealSAT 30 m) | Reservoirs (GRanD, 1990–2017; 0.03 km) | - | - | - | - | - | - |
| **Pollution** | Nitrous oxide pollution (Sentinel 5P 0.5 km; EDGAR, 1990–2012; 100 km) | Nitrous oxide pollution (EDGAR, 1990–2012; 100 km) | - | - | - | - | - | - |

| | | | | | | | | |
|---|---|---|---|---|---|---|---|---|
| **Transportation** | Road (highway, minor, two-track; OSM, 2024); railways (OSM, 2024) land use (OSM, 2024); power lines (OSM, 2024); electrical power infrastructure (harmonized DMSP and VIIRS, 1992–2023) | Road (highway, minor, two-track; OSM, 2019); railways (OSM, 2019) power lines (OSM, 2019); electrical power infrastructure (harmonized DMSP and VIIRS, 1992–2018) | Road (highway, minor, two-track; OSM, 2016; gROADS-2000); railways (OSM, 2016; VMAP-2000), power lines (OSM, 2016); electric infrastructure (nighttime lights; DMSP OLS, 2013) | - | Roads and railways (no differentiation of type) from OSM (2021 source). Mapped as 80% with 0.5 buffer VIIRS nightime lights (using 25 as maximum) | Roads (gROADS, 1980–2000) and Open Streetmap (OSM) railways (VMAP0-2000). Mapped as 40% 1 to 15 km decay. | Roads (gROADS, 1980–2000) railways (VMAP0-2000). Mapped as 40% 1 to 15 km decay. | Roads (VMAP 1990s) |
| **Analyses** | | | | | | | | |
| **Integration formula** | Increasive to 1.0 using fuzzy sum (averaging multiple datasets into individual threats) | Increasive to 1.0 using fuzzy sum (averaging multiple datasets in a threat) | Increasive to 1.0 using fuzzy sum | Overlayed excluded areas (added WDPA Cat I-IV) | Summed (additive) | Summed (additive) | Summed (additive) | Summed (additive) |
| **Metric** | H; 0–1.0 continuous value | H; 0–1.0 continuous value | H; 0–1.0 continuous value | 0/1 | Ordinal value | Ordinal value | Ordinal value | - |
| **"Wild"** | H<0.084 (weighted mean H in WDPA Cat I-II) | H<0.084 (mean H in WDPA Cat I-II) | H<0.084 (mean H in WDPA Cat I-II) | 0/1 | HF < 4 | HF < 4 | HF < 4 | 10% cutoff (HF 3-18) |
| **Uncertainty analysis** | Calculates per-pixel variance due to estimates of intensity values, randomized (n=100). Calculate | Calculates per-pixel variance due to estimates of intensity values, randomized (n=50) | Calculates per-pixel variance due to estimates of intensity values, randomized (n=100) | - | - | - | - | - |

| | | | | | | | | |
|---|---|---|---|---|---|---|---|---|
| | uncertainty associated with primary land user/cover datasets. | | | | | | | |
| **Validation** | Tested using independent validation dataset that included ~10 000 subplots within ~1000 0.36 km2 sample plots stratified by urban, exurban, and rural areas. | Tested using independent validation dataset that included ~10 000 subplots within ~1000 0.36 km2 sample plots | Tested using independent validation dataset that included ~10 000 subplots within ~1000 0.36 km2 sample plots | Tested using independent validation, random distribution of 3,460 1 km2 sample plots | Tested using independent validation, random distribution of 3,460 1 km2 sample plots | Tested using independent validation, random distribution of 3,460 1 km2 sample plots | Tested using independent validation, random distribution of 3,460 1 km2 sample plots | - |
| **Assumption of edge-effects** | Not modeled directly, and so provides foundation for subsequent fragmentation and connectivity analyses | Not modeled directly, and so provides foundation for subsequent fragmentation and connectivity analyses | Not modeled directly, and so provides foundation for subsequent fragmentation and connectivity analyses | - | Buffer 15 km from roads (declining) | Buffer 12 km from roads (declining); 10 km smoothing on cropland/pasture ? | Buffer 12 km from roads (declining); 10 km smoothing on cropland/pasture ? | Buffer 12 km (declining) |
